# Monomeric agonist peptide/MHCII complexes activate T-cells in an autonomous fashion

**DOI:** 10.1101/2023.03.13.532401

**Authors:** René Platzer, Joschka Hellmeier, Janett Göhring, Iago Doel Perez, Philipp Schatzlmaier, Clara Bodner, Gerhard J. Schütz, Eva Sevcsik, Hannes Stockinger, Mario Brameshuber, Johannes B. Huppa

**Affiliations:** Medical University of Vienna, Center for Pathophysiology, Infectiology, Immunology, Institute for Hygiene and Applied Immunology, Vienna, Austria; TU Wien, Institute of Applied Physics, Vienna, Austria

**Keywords:** MHC class II, T-cell antigen recognition, molecular imaging

## Abstract

Molecular crowding of agonist peptide/MHC class II complexes (pMHCIIs) with structurally similar, yet *per se* non-stimulatory endogenous pMHCIIs has been postulated to sensitize T-cells for the recognition of single antigens on the surface of dendritic cells and B-cells. When testing this premise with the use of advanced live cell microscopy, we observed pMHCIIs as monomeric, randomly distributed entities diffusing rapidly after entering the APC surface. Synaptic TCR-engagement of highly abundant endogenous pMHCIIs was low or non-existent and affected neither TCR-engagement of rare agonist pMHCII in early and advanced synapses nor agonist-induced TCR-proximal signaling. Our findings highlight the capacity of single freely diffusing agonist pMHCIIs to elicit the full T-cell response in an autonomous and peptide-specific fashion with consequences for adaptive immunity and immunotherapeutic approaches.

**SHORT SUMMARY:** Platzer et al. revealed via highly quantitative and single molecule live cell microscopy the nature of peptide-loaded MHC class II molecules (pMHCII) as monomeric, densely populating, randomly distributed and predominantly rapidly diffusing entities on the surface of B-cells and dendritic cells. Low abundant stimulatory agonist pMHCII acted as autonomous units with the highest chance of T-cell detection when equally spread on APCs. The presence of bystander-pMHCII previously termed “co-agonist pMHC” affected neither synaptic agonist -TCR-binding nor efficiencies of T-cell recognition. “Co-agonist”-TCR-binding resembled random molecular collisions. Findings inform the design of T-cell-based immunotherapies.

## INTRODUCTION

The recognition of antigenic peptides by T-cells relies on transient interactions with antigen presenting cells or target cells (APC). The formation of this specialized cell contact – termed the immunological synapse – is driven by specific T-cell antigen receptors (TCRs) on the T-cell side engaging peptide-loaded MHC complex molecules (pMHCs) (Bromley et al., 2001; Huppa and Davis, 2003). In this setting T-cells can detect the presence of even a single stimulatory pMHC molecule amongst thousands of structurally similar yet non-stimulatory or endogenous pMHCs on a densely packed APC surface (Huang et al., 2013; Irvine et al., 2002; Li et al., 2004; Purbhoo et al., 2004). The molecular, biophysical, and cellular mechanisms underlying this phenomenon are still unclear, in particular when considering that most T-cell antigen receptors (TCR)-engage their nominal ligands with a low to moderate affinity (1–100 µM), at least when measured *in vitro* (Corr et al., 1994; Davis et al., 1998; Matsui et al., 1991). Previous studies have suggested that ligand recognition may not only depend on the intrinsic binding properties of TCRs and pMHCs but also on the quality and quantity of accessory interactions (i.e. CD28/B7-1, CD2/CD48, LFA-1/ICAM-1, CD4 and CD8 co-receptors), as well as cell biological parameters such as TCR-subunit or pMHC stoichiometry, local molecular crowding and cytoskeletal forces (reviewed in (Göhring et al., 2022; Platzer and Huppa, 2020)).

Key to understanding the molecular foundation of T-cell antigen recognition is to examine TCR-pMHC engagement *in situ*, as conditions for binding within the synapse differ in many regards from those found in solution (Huang et al., 2010; Huppa et al., 2010). If positioned in sufficient proximity, TCRs and pMHCs engage each other on opposed plasma membranes in a pre-oriented fashion which appears to accelerate the binding on-rate (Huang et al., 2013; Huppa et al., 2010; Weikl et al., 2016). Synaptic TCR-pMHC lifetimes are furthermore affected by simultaneous engagement of co-receptors with the same pMHC (Huppa et al., 2010; Jiang et al., 2011) as well as cellular forces imposed on the binding partners (Feng et al., 2017; Gohring et al., 2021; Liu et al., 2014; Pettmann et al., 2022; Sibener et al., 2018). Furthermore, both pMHCI and pMHCII have been suggested to be organized in clusters (Lu et al., 2012) where they may undergo lateral interactions, which could alter binding and signaling dynamics. In fact, nanoscale MHCII clustering through interaction with tetraspanins or partitioning into glycosphingolipid rafts has been considered critical for accelerated T-cell scanning and sensitized antigen recognition (Anderson et al., 2000; Bosch et al., 2013; Zuidscherwoude et al., 2015). pMHCII enrichment has furthermore been suggested to drive the formation of pMHCII higher order structures, in particular in view of the tendency of some pMHCIIs to crystallize as dimers (Brown, 1993; Fremont et al., 2002). Consistent with this, soluble agonist pMHCIIs activate antigen-experienced T-cells only when tethered to a second agonist pMHCII or to an endogenous pMHCII capable of TCR binding (Cochran et al., 2000; Krogsgaard et al., 2005). Rather than confounding the process of antigen recognition, highly abundant endogenous pMHCs have hence been proposed to facilitate the detection of rare agonist pMHCIIs in a so-termed “pseudodimer” arrangement (Anikeeva et al., 2012; Krogsgaard et al., 2005; Wulfing et al., 2002) whereby such pMHCs may act – without being able to stimulate T-cells on their own – as so-termed “co-agonists”.

We have recently shown via molecular live cell imaging that TCRs are organized prior to pMHCII engagement in a randomized fashion (Platzer et al., 2020; Rossboth et al., 2018) and function as monomeric antigen-binding and signaling entities (Brameshuber et al., 2018). Therefore, any co-agonistic effect mediated by a pseudodimer arrangement would have to result from agonist pMHCIIs being either displayed as part of the same higher-order MHCII structure or densely packed in membrane nanodomains. However, to this date neither the presence of MHCII-enriched membrane domains, nor the existence of pMHCII dimers or higher order structures has been demonstrated on the surface of living APCs, and hence the relevance of pMHCII-clustering for T-cell sensitization remains to be determined (Anikeeva et al., 2012; Gascoigne, 2008; Hoerter et al., 2013; Ma et al., 2008; Nagy et al., 2001).

To do so we have conducted minimally invasive single-molecule imaging on living professional APCs with sub-diffraction resolution. We failed to obtain any evidence for the existence or formation of higher-order pMHCII structures prior to or in the course of antigen recognition and ruled out any relevance of such entities for antigen recognition if generated experimentally. Instead, we observed pMHCIIs as monomeric entities randomly distributed at high densities across the plasma membrane and diffusing rapidly immediately after entering the surface of bone marrow-derived dendritic cells and B-cells. When measuring synaptic TCR-pMHCII interactions with the use of a Förster Resonance Energy Transfer (FRET)-based assay we did not find the presence of highly abundant endogenous bystander-pMHCIIs to be relevant for the TCR-engagement of rare agonist pMHCII at early and later stages of synapse formation and the ensuing TCR-proximal signaling. Instead, we witnessed low level calcium signaling in T-cells facing “co-agonistic” altered peptide ligands (APLs) at supraphysiological densities (above 1000 molecules per µm^2^), yet without contributing to T-cell sensitization towards rare antigenic pMHCIIs. In summary, our findings challenge the concept of pMHCII co-agonism and underscore the stimulatory potency of single, freely diffusing agonist pMHCIIs, which activate cognate T-cells in a peptide-specific manner without any significant contribution from endogenous bystander-pMHCIIs.

## RESULTS

### Dendritic cells and B-cells express pMHCIIs at high numbers and densities

We first determined the number of pMHCIIs that T-cells can in principle encounter while scanning the surface of professional APCs. For precise quantitation we employed a monovalent site-specifically labeled and in selected cases also biotinylated recombinant single chain antibody fragment (scF_V_) derived from the I-E^k^ -reactive 14.4.4S mAb (**Supp. Fig. 1A-E,** for details see Methods section). As verified by flow cytometry, 14.4.4 scF_V_ binding to I-E^k^ expressed on CH27 B-cells was specific, efficient and stable: when employed at a concentration of 5 µg ml^-1^ 14.4.4 scF_V_ conjugated to Alexa Fluor 647- or Alexa Fluor 488 (14.4.4 scF_V_-AF647 or AF488) decorated more than 95% of surface expressed I-E^k^ with half-lives of staining amounting to 41 minutes at 37°C, 80 minutes at 25°C and 16 hours on ice (**Supp. Fig. 1C-E**). To further confirm target-specificity of the 14.4.4 scF_V_, we co-cultured 5c.c7 TCR-transgenic T-cells with moth cytochrome c (MCC 88-103) peptide-pulsed BMDCs at 37 °C in the presence of 20 µg ml^-1^ 14.4.4 scF_V_. T-cells no longer secreted interferon-*γ*(IFN-*γ*) and tumor necrosis factor (TNF-*α*). EC50 values derived from a peptide titration series in the presence or absence of the 14.4.4 scF_V_ indicated an epitope coverage of 98.6% over a period of 4 hours. (**Fig. 1A** **and Supp. Fig 2A).**

**Figure 1:**
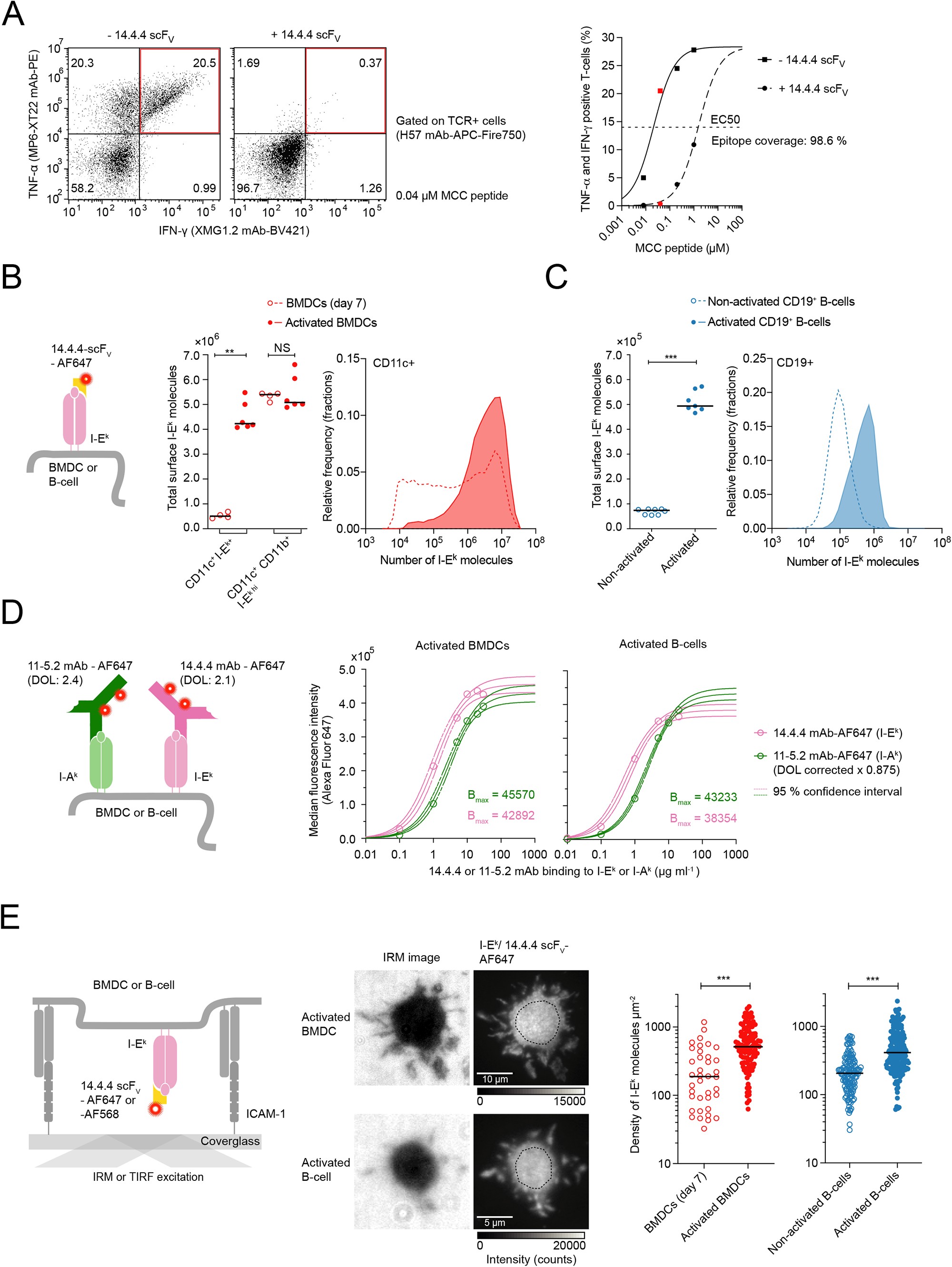
Dendritic cells and B-cells express pMHCIIs in high numbers and high surface densities. **(A)** Efficient blockade of T-cell antigen recognition by the 14.4.4 scF_V._ Shown is the flow cytometric analysis of antigen-induced TNF-*α* and IFN-*γ* expression in 5c.c7 TCR-transgenic T-cells co-cultured with BMDCs which had been pre-pulsed with MCC peptide antigen (from 0 to 1 µM) in the absence or presence of the 14.4.4 scF_V_ with exemplary dot plot depicted on the left. **(B)** I-E^k^ is expressed at high numbers on activated CD11c^+^ CD11b^+^ BMDCs. **Left panel**: staining procedure employed for quantitation. **Middle panel**: the median number of I-E^k^ molecules displayed on the surface of non-activated and activated BMDCs, gated as CD11c^+^ I-E^k+^ or CD11c^+^ CD11b^+^ I-E^k hi^, as analyzed through quantitative flow cytometry (Mann-Whitney-U-test). Data was pooled from two independent recordings. **Right panel**: histogram depicting the distribution of surface-expressed I-E^k^ molecules on non-activated (dashed) and activated (solid) CD11c^+^ BMDCs (binning = 0.15 (log scale), n = 14585 non-activated BMDCs, n = 24214 activated BMDCs). Data are representative of two independent recordings. **(C)** I-E^k^ is expressed at high numbers on non-activated and activated CD19^+^ B-cells. **Left panel**: the median number of I-E^k^ molecules displayed on the surface of non-activated and activated CD19^+^ B-cells as analyzed through quantitative flow cytometry (Mann-Whitney-U-test). Data was pooled from three independent recordings. **Right panel**: histogram depicting the distribution of I-E^k^ molecules expressed on non-activated and activated CD19^+^ B-cells (binning = 0.15 (log scale), n = 14032 non-activated CD19+ B-cells, n = 10010 activated CD19+ B-cells). Data are representative of three independent recordings. **(D)** I-A^k^ and I-E^k^ molecules are expressed on the surface of activated CD11c^+^ BMDCs and activated CD19^+^ B-cells at similar levels. **Left panel**: staining procedure employed for quantitation. **Right panel**: I-A^k^- and I-E^k^-expression levels as determined on the basis of I-E^k^ surface-expression after correcting for the degree of fluorophore conjugation of the antibodies employed. The mAb titration curves were fitted to a one-site specific binding model to estimate B_max_ and the 95% confidence interval. Goodness of fit R^2^ > 0.99. **(E)** High I-E^k^ densities as measured via TIRF-microscopy on the surface of activated BMDCs and B-cells. **Left panel**: Scheme illustrating 14.4.4 scF_V_-AF647-based staining as well as Interference Reflection Microscopy (IRM)-and Total Internal Reflection Fluorescence (TIRF)-based microscopy readout to confirm attachment and spreading of cells on the ICAM-1-coated glass surface and density quantitation, respectively. **Middle panel**: Example of IRM and TIRF images of indicated cells on ICAM-1-functionalized glass slides. **Right panel**: Dot plot of I-E^k^ surface densities on non-activated BMDCs (median, n = 38 cells) and activated BMDCs (median, n = 113 cells) and non-activated B-cells (median, n = 141 cells) and activated B-cells (median, n = 188 cells). Data were pooled from three (BMDCs) and five (B-cells) independent experiments. Statistics: median, unpaired two-tailed student’s t-test. P value format: P > 0.05 (ns), P ≤ 0.05 (*), P ≤ 0.01 (**), P ≤ 0.001 (***).

Total I-E^k^ surface-expression was quantitated based on flow cytometric median fluorescence intensities (MFIs) of 14.4.4 scF_V_-AF647 stained BMDCs and B-cells, which we then normalized to measured fluorescence intensities of AF647 quantification beads serving as reference (Supp. **Fig. 2B**). As shown in **Fig. 1B**, non-activated day 6 to 7 CD11c^+^ I-E^k+^ BMDCs expressed an average of 0.5 million copies of I-E^k^ at the surface per cell, which increased to 4-5 million copies per BMDC upon activation with lipopolysaccharide (LPS). Indicative of partial BMDC maturation prior to LPS exposure, I-E^k^ expression levels of non-activated BMDCs (7 days after bone marrow isolation) were a highly heterogeneous (Supp. **Fig. 3A**), with CD11c^+^ CD11b^+^ I-E^k hi^ BMDCs representing mainly mature DCs (Helft et al., 2015). At least in part owing to their significantly smaller size, CD19^+^ B-cells displayed about ten-to 50-fold fewer copies of I-E^k^, i.e. on average ∼1×10^5^ molecules per non-activated and ∼5x10^5^ molecules per B-cell stimulated with LPS for 48 hours (**Fig. 1C**, **Supp. Fig. 3B-C**).

**Figure 2:**
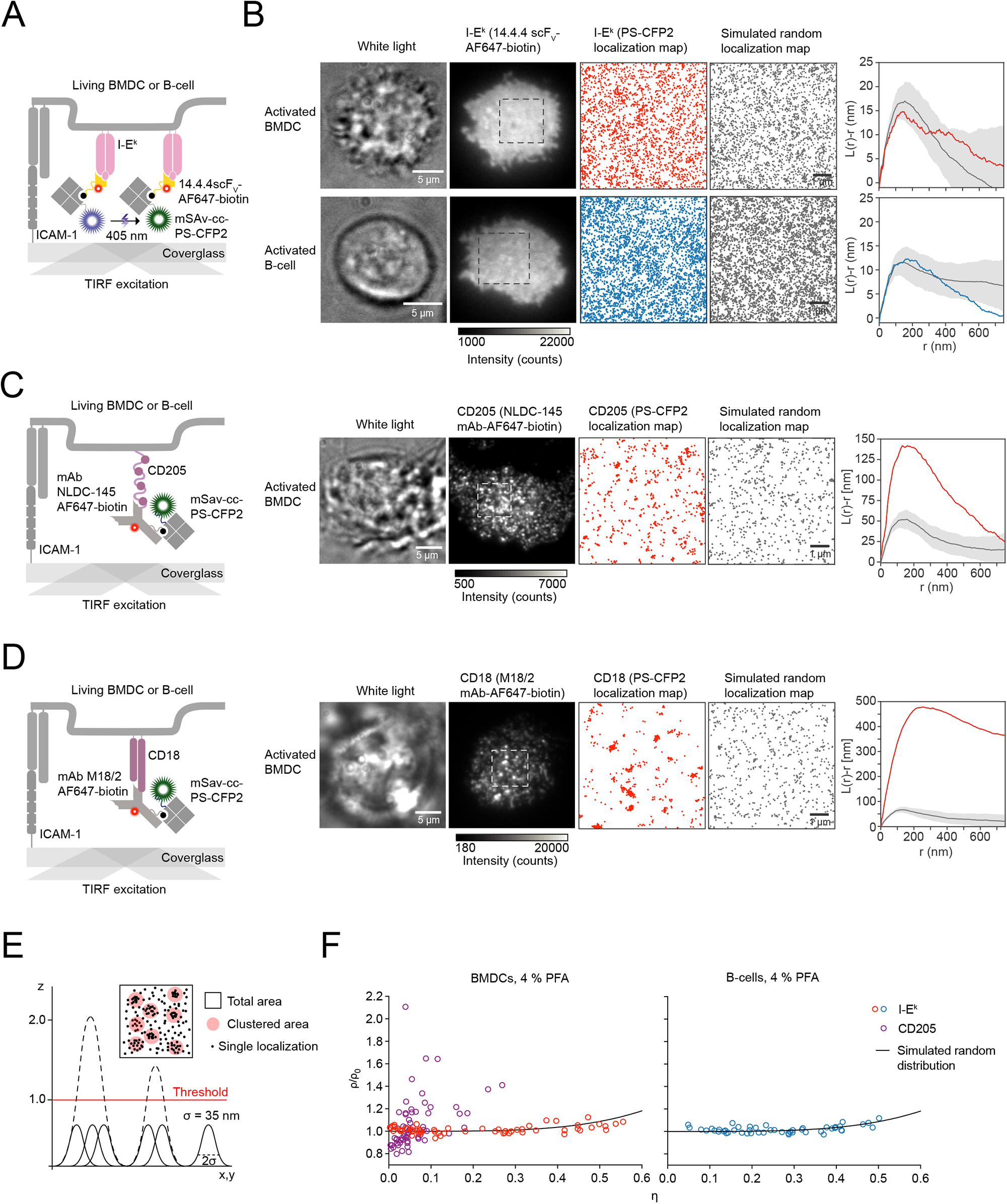
Surface I-E^k^ molecules randomly distributed on the surface of activated BMDCs and B-cells. **(A)** Staining procedure to image the surface distribution of I-E^k^ on living professional APCs below the diffraction limit of visible light using hsPALM. **(B)** BMDCs and B-cells were stained with 14.4.4 scF_V_-AF647-biotin and mSav-cc-PS-CFP2 and seeded onto ICAM-1 coated glass slides for imaging in TIRF mode. The presence of each cell was verified through acquisition of a white light image (first image, scale bar 5 µm), a single AF647 ensemble fluorescence image (second image), followed by 3050 image frames displaying single PS-CFP2 fluorescence events within 7 seconds to be assembled as a localization map (third image), which was compared to a simulated random I-E^k^ localization map (fourth image, scale bar 1 µm). Ripley’s K analysis was used to compare BMDC-derived I-E^k^ distributions (red) or B-cell-derived I-E^k^ distributions (blue) to ten simulations of randomized distributions with corresponding molecular densities, diffusion parameters and blinking statistics (gray). Statistics: mean ± standard deviation. **(C, D)** Quantification of the surface-distribution of (**C**) DEC205 and (**D**) CD18 on activated BMDCs. **Left panels**: Staining and imaging procedure. **Right panels**: BMDCs were stained with the indicated mAbs (NLDC145 mAb-AF647-biotin or M18/2 mAb-AF647-biotin) and mSav-cc-PS-CFP2 for hsPALM imaging. Ripley’s K analysis was used to compare cell-derived DEC205 distributions (red) to ten simulations of randomized distributions with corresponding molecular densities, diffusion parameters and blinking statistics (grey). Statistics: mean ± standard deviation. **(E)** Principle of label density variation analysis. A theoretical spatial uncertainty (*σ*) of 35 nanometers was assigned to each PS-CFP2 localization and overlapping Gaussian profiles were merged (and intensities were added up) to arrive at three-dimensional cluster maps (X, Y position, intensity). Different thresholds were assigned to probe derived cluster maps and calculate ρ (the density of localizations in the clustered area per µm^2^) and η (clustered area divided by total area). **(F)** Label density variation analysis is consistent with a random surface distribution of I-E^k^ on activated BMDCs and B-cells and a clustered surface distribution of DEC205 on activated BMDCs, serving as positive control. Quantification of the relative clustered area (η) and the density of localizations per clustered area (ρ) for indicated molecular species at a threshold of 1.0. The black line indicates the reference curve for a random distribution.

**Figure 3:**
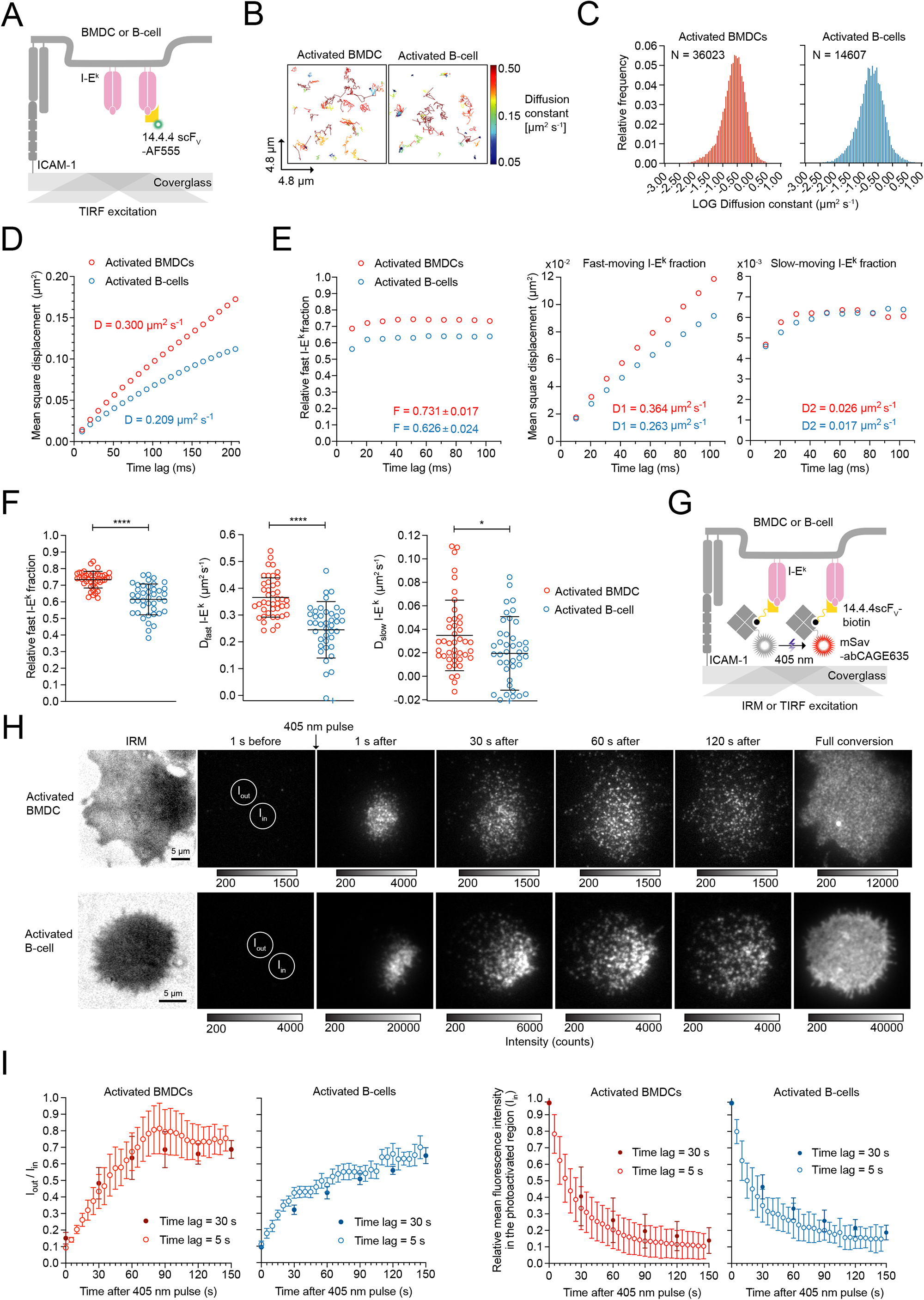
Surface resident pMHCIIs are to 99% mobile with 60-75% displaying fast diffusion immediately after entering the cell surface. **(A)** Schematic illustration of a single molecule tracking experiment to monitor the diffusion of single I-E^k^ molecules within the plasma membrane of activated BMDCs and B-cells at 37°C. Cells were stained with an I-E^k^-specific AF555-conjugated 14.4.4 scF_V_ and seeded on ICAM-1-coated glass slides. Single I-E^k^ molecules were tracked over time with a time lag of 10.2 or 10.5 milliseconds to study the mobility of I-E^k^. **(B)** Visualization of single I-E^k^ trajectories on the surface of representative BMDCs and B-cells. Trajectories were color-coded according to their lateral speed. **(C)** Histogram of diffusion constants derived from individual I-E^k^ trajectories on BMDCs and B-cells. **(D)** Mean square displacement plot of recorded I-E^k^ trajectories. D = diffusion constant (fitted through the first two data points). Statistics: mean and standard deviation; number of trajectories n = 36023 (BMDCs); n = 14607 (activated B-cells) **(E)** Binary diffusion model to describe heterogenous I-E^k^-diffusion. **Left panel**: Relative fraction of fast- and slowly-moving I-E^k^ molecules on activated BMDCs and B-cells were calculated by fitting the recorded I-E^k^ trajectories to a binary diffusion model. **Middle and right panel**: Mean square displacement plot of fast-moving I-E^k^ molecules (middle panel) and slow-moving I-E^k^ molecules (right panel). Statistics: mean and standard deviation. F = fraction of fast-moving molecules. D1 = diffusion constant of fast-moving molecules. D2 = diffusion constant of slow-moving molecules. Diffusion constants were extracted by fitting the first two data points. Fractions were calculated by fitting the first ten data points. **(F)** Quantification of the binary diffusion data on individual activated BMDCs and B-cells. Each circle represents the mean of all trajectories of a single cell after being categorized by the binary fit. Data points outside the axis limits are marked with a plus (+). Statistics: mean and standard deviation; unpaired two-tailed student’s t-test. P value format: P > 0.05 (ns), P ≤ 0.05 (*), P ≤ 0.01 (**), P ≤ 0.001 (***), P ≤ 0.0001 (****). **(G, H, I)** I-E^k^ spread evenly with high velocity on the cell surface of activated BMDCs and B-cells. **(G)** Scheme illustrating the use of inverse Fluorescence Recovery after Photo-bleaching (iFRAP) to assess cell surface spreading of I-E^k^. Activated BMDCs were stained with site-specifically biotinylated 14.4.4 scF_V_ (14.4.4 scF_V_-biotin) for detection with site-specifically conjugated mSav-abCAGE635, a photoswitchable and photostable bright fluorophore that allowed for tracking of individual molecules from a particular region of interest after photoswitching with a short 405 nm laser pulse. Stained BMDCs were seeded onto ICAM-1-coated glass slides for recordings in TIRF mode at 37°C. **(H)** Shown are representative time lapse montages of I-E^k^-conjugated mSav-abCAGE635 photo-converted within the indicated circular ROI (I_in_) on the surface of an activated BMDC and B-cell. **(I)** Over 70-90% of I-E^k^-bound mSav-CAGE635 dispersed over the entire APC surface within 150s after photo-conversion of a small region of interest. **Left panel:** Shown is the ratio between the background-subtracted mean intensity of a dark cell region (I_out_) and the photoconverted region (I_in_) recorded either with a long (30 seconds) or a short (5 seconds) time delay. Statistics: mean and standard error of the mean of 10 to 14 cells (for each delay) from one experiment. **Right panel:** Decline of the fluorescence intensity within the photoactivated region (I_in_). Shown is mean fluorescence intensity which had been background-subtracted and corrected for photobleaching (3% per image). Statistics: mean and standard deviation of 10 to 14 cells (for each delay) from one experiment.

To account for all surface-resident pMHCIIs that are in principle available for T-cell scanning, we also determined the total number of displayed I-A^k^-copies indirectly through comparison of MFIs obtained from APCs stained with saturating quantities of the 14.4.4 mAb and the I-A^k^ reactive 11-5.2 mAb (**Fig. 1D**). After normalizing MFIs for degrees of antibody labeling (DOL; 14.4.4 mAb = 2.1; 11-5.2 mAb = 2.4) we arrived at similar numbers of I-A^k^ and I-E^k^ that were accessible to antibody-staining on the surface of BMDCs and B-cells.

To assess whether endogenous pMHCIIs exerted any influence on the detection of rare agonist ligands, we sought to determine the number of bystander-pMHCII in close vicinity to any scanning or ligated TCR. Of note, both BMDCs and B-cells gain considerably in surface area upon activation, while B-cells increase in overall size (blast formation), the surface of BMDCs changes from microvilli to veils (Verdijk et al., 2004). As a result, total cellular expression levels cannot easily be converted into molecular densities due to varying cell size and plasma membrane convolutions. We hence stained BMDCs and B-cells with saturating amounts of 14.4.4 scF_V_ site-specifically conjugated with Alexa Fluor 647 or Alexa Fluor 568, and then allowed the cells to spread on Intercellular Adhesion Molecule 1 (ICAM-1)-coated glass slides as a means to directly measure surface I-E^k^ densities using total internal reflection fluorescence (TIRF) microscopy (**Fig. 1E**). Image acquisition was conducted at room temperature (22.5°C) to minimize probe dissociation (Supp. Fig. 1D). Tight alignment of the APC plasma membrane with the glass slide was verified by Interference Reflection Microscopy (IRM). We then determined average I-E^k^ densities within a central region of interest (ROI) randomly picked from the cell-glass contact region. For this we divided the integrated background-subtracted fluorescence intensity by the mean intensity of a single molecule fluorescent signal and the area of the ROI (**Fig. 1E**, for more details refer to Material and Methods). We arrived at a median I-E^k^ surface density of 188 molecules per µm^2^ (minimum: 33 µm^-2^, maximum: 1179 µm^-2^) on non-activated BMDCs and 206 molecules per µm^2^ (minimum: 30 µm^-2^, maximum: 723 µm^-2^) on non-activated B-cells (**Fig. 1E**). After LPS treatment we noticed an approximately two-fold increase in I-E^k^ surface densities on BMDCs (median: 514 µm^-2^, minimum: 63 µm^-2^, maximum: 1991 µm^-2^) and B-cells (median 413 µm^-2^, minimum: 61 µm^-2^, maximum: 2346 µm^-2^).

### I-E^k^ molecules fail to associate with detergent-resistant membranes and are randomly distributed on the surface of activated BMDCs and B-cells

pMHCII have previously been described to be sequestered to so-termed lipid rafts, a controversially-debated class of small and highly dynamic membrane domains consisting of cholesterol, sphingolipids, saturated fatty acids, glycosylphosphatidylinositol (GPI)–anchored and cysteine-palmitoylated proteins (Anderson et al., 2000; Kusumi et al., 2012; Sevcsik et al., 2015). To assess possible mechanisms underlying lipid raft-dependent T-cell sensitization as proposed by others (Anderson et al., 2000; Bosch et al., 2013), we quantitated the extent to which pMHCIIs on the surface of BMDCs and B-cells are found in so-termed detergent-resistant membranes (DRMs). Protein presence in DRMs has previously been regarded as the biochemical equivalent of lipid raft residence. To this end we conducted lysis gradient centrifugation (LGC) affording comprehensive results with a sample size of fewer than 10,000 cells when combined with a flow cytometric readout (Schatzlmaier et al., 2015). The LGC methodology rests on quantitative co-isolation of DRMs and their associated proteins with the detergent-resistant nuclei of their parental detergent-treated cells (**Supp. Fig. 4A**). As shown in **Supp. Fig. 4B-C** we analyzed surface-resident I-E^k^ or I-A^k^ molecules on non-activated and activated BMDCs and B-cells as well as the GPI-anchored DRM markers CD44 and CD55 as positive control, and the non-DRM resident CD71 as negative control. As expected, CD44 and CD55 decorated with fluorescently labeled mAbs were predominantly present within the DRM fraction while CD71 was absent, testifying to the technical integrity of the experiment. Strikingly, I-E^k^ labeled with either the 14.4.4-scF_V_-AF647 or 14.4.4 mAb-AF647 was hardly detectable in DRMs (< 1%, see **Supp. Fig. 4C**), a behavior we could also associate with I-A^k^ decorated with the 11-5.2 mAb-AF647 on the surface of both BMDCs and B-cells. Contrasting previous reports, these results indicate that MHCII, at least when present on the surface of primary B-cells and BMDCs, are predominantly absent from or only very weakly associated with DRMs (< 1%).

To examine whether pMHCIIs are enriched in micro- or nanoclusters other than lipid rafts, we stained living activated BMDCs and B-cells with randomly AF647-conjugated I-E^k^ and I-A^k^-reactive mAbs or site-specifically AF647-conjugated 14.4.4 scF_V_s and analyzed cellular mid sections via fluorescence microscopy.

We continued our study by focusing primarily on activated BMDCs given the pivotal role in T-cell priming. For comparative analysis, we also included activated B-cells in our study. Using the 14.4.4 scF_V_ or 14.4.4 mAb (or 11-5.2 mAb) as probe, deconvolved images revealed a uniform staining of I-E^k^ or I-A^k^. As is consistent with a previous study (Bosch et al., 2013) we observed a punctate staining pattern on BMDCs after exposing surface-resident mAb-engaged I-E^k^ molecules with secondary AF647-conjugated IgG mAbs. Taken together, we concluded that pMHCIIs are not organized in plasma membrane microdomains that are large enough to be resolvable with diffraction-limited microscopy (**Supp. Fig. 5A-B**).

To determine whether pMHCIIs are enriched in plasma membrane nanodomains, which can no longer be identified by conventional fluorescence microscopy, we analyzed the surface distribution of I-E^k^ not only on living (**Fig. 2**) but also on paraformaldehyde (PFA)-fixed (**Supp. Fig. 8**) BMDCs and B-cells with sub-diffraction resolution applying high-speed Photoactivation Localization Microscopy (hsPALM). We included cell fixation in half of our analyses since the high lateral diffusion of pMHCII may compromise cluster detection via hsPALM. Of note, PALM relies on the stochastic photoswitching of suitable fluorophores between a blue-shifted fluorescent or dark state and a red-shifted or bright fluorescent state and the subsequent reconstruction of diffraction-limited but well-separated single molecule events into a localization map (Betzig et al., 2006, Hess et al., 2006). For visualization of I-E^k^, we site-specifically conjugated the 14.4.4 scF_V_ with AF647 and biotin (14.4.4 scF_V_-AF647-biotin, **Fig. 2A** **and Supp. Fig. 1B**). Cell surface-resident 14.4.4 scF_V_-AF647-biotin was then detected in hsPALM with the use of monovalent streptavidin (mSav) site-specifically conjugated with recombinant PS-CFP2 (mSav-cc-PS-CFP2, **Fig. 2A** **and Supp. Fig. 6A-E)**, a photoswitchable fluorophore with a moderate repetitive blinking behavior (Platzer et al., 2020).

To allow for noise-reduced imaging in TIRF mode we seeded living BMDCs and B-cells labeled with 14.4.4 scF_V_-AF647-biotin and mSav-cc-PS-CFP2 on ICAM-1-coated glass-slides. Prior to conducting hsPALM, attachment of each cell was verified through acquisition of a single AF647 ensemble fluorescence image, which also served to assess surface I-E^k^ distribution at a diffraction-limited level (**Fig. 2B**). For hsPALM we recorded 3050 image frames which displayed single diffraction-limited PS-CFP2 fluorescence events within a time frame of seven seconds. Diffraction-limited single molecule signals were then fitted with a Gaussian intensity profile accounting for the point spread function, spatially positioned with an accuracy of 15 to 108 (mean value 51 ± 12) nanometers (**Supp. Fig. 6F**) and then superimposed to give rise to a composite localization map.

Of note, the interpretation of single-molecule localization microscopy (SMLM) data is typically complicated by overcounting and undercounting artefacts, which result from insufficient photobleaching (on-off blinking) and variability in psCFP-maturation. Fluorophore blinking can give rise to false-positive nanocluster detections in the reconstructed localization map (Annibale et al., 2011; Baumgart et al., 2016). To distinguish true protein nanoclusters from overcounting artifacts, we first employed an imaging platform to quantitatively determine the blinking signature of PS-CFP2 prior to PALM measurements on cells (Platzer et al., 2020). To promote the appearance of PS-CFP2 in a single imaging frame only, we employed a 488 nm laser power density of 3.5 kW cm^-2^ and recorded the images in stream mode (488 nm laser is continuously illuminating) with an illumination time of two milliseconds. (**Supp. Fig. 7A**). Applying a Monte Carlo simulation-based approach we quantitatively related cell-derived PALM images to localization maps, which had been computed from simulated random distributions featuring cell-derived molecular densities, the diffusion parameters of I-E^k^ at 25°C (**Supp. Fig. 7B**) and the respective blinking signature of PS-CFP2 (for more details see Material and Methods). As shown in **Fig. 2B**, comparing respective Ripley’s K maxima of recorded I-E^k^ localization maps with those derived from simulated random localization maps did not reveal any significant differences, which let us conclude that pMHCIIs are homogenously distributed on the surface of activated BMDCs and B-cells, at least when judged within the cluster detection sensitivity of our approach (**Supp.** **Fig. 7A**). Importantly and in contrast, we could detect local surface enrichments for the internalization receptor DEC205 and for the integrin beta chain CD18 (**Fig. 2C-D** **and Supp. Fig. 7C-D)**. Confirming the sensitivity and robustness of our imaging and analysis methodologies, background localizations registered only in negligible numbers in the absence of PS-CFP2 (**Supp. Fig. 7E**). To circumvent potential complications arising from molecular diffusion and cellular motility, we performed analogous imaging and analysis procedures on PFA-fixed activated BMDCs and B-cells. As shown in **Supp. Fig. 8A-B**, we identified inhomogeneities and thus small deviations from a complete random distribution on PFA-fixed cells, which could also arise from the fixation procedure itself.

We hence carried out artefact-free label density variation SMLM (Baumgart et al., 2016) to quantitate the extent to which I-E^k^ is organized in nanoscale entities within I-E^k^ localization maps derived from PFA-fixed activated BMDCs and B-cells. To this end and based on image analysis, we assigned a spatial uncertainty (*σ*) of 35 nanometers to each PS-CFP2 localization and merged overlapping Gaussian profiles to three-dimensional cluster maps (X, Y position, intensity, **Fig. 2E**). We calculated localization densities within an assigned clustered area (ρ) and the relative clustered area to the total area (η) for a given threshold (1.0) and plotted the density of localizations in the clustered area per µm^2^ (ρ) against η (clustered area divided by total area). Results correlated best with a random distribution of I-E^k^ on activated PFA-fixed BMDCs and B-cells, in particular because the localization density within the clustered area did not significantly deviate with higher staining degree from a simulated random distribution (solid line), unlike what would be expected from a clustered distribution (**Fig. 2F**). In contrast and consistent with previous studies, density variation SMLM revealed clustering of DEC205 on BMDCs, with an increase of the localization density within clusters at higher expression or staining levels (**Fig. 2F**).

Lastly, to avoid any issues potentially arising from SMLM and data interpretation, we employed stimulated emission depletion (STED) microscopy. STED is not susceptible to artifacts resulting from overcounting, as it resolves structures below the diffraction limit of visible light by reducing the width of the point spread function to several tens of nanometers through high powered stimulated emission depletion (Sahl et al., 2017). For this purpose, we stained activated BMDCs and B-cells with a 14.4.4-scF_V_ which had been site-specifically conjugated with abSTAR635P (14.4.4-scF_V_-abSTAR635P), a STED-compatible fluorophore, and allowed the cells to spread on ICAM-1-coated glass slides prior to PFA-mediated fixation (**Supp. Fig. 9A-B and Supp. Fig. 10**). To assess the degree of pMHCII clustering in STED microscopy images, we compared the STED recordings with simulated images of randomly distributed molecules obtained with the fitted parameters of single fluorescence emitters derived from activated BMDCs and B-cells labelled with single 14.4.4 scF_V_-abSTAR635P molecules (**Supp. Fig. 9A**). To this end, we employed image-autocorrelation function (ACF) analysis, which calculates the probability that two pixels separated by a distance r have a similar brightness, and thus enables quantification of the spatial distribution of fluorophores in the recorded STED images (Robertson and George, 2012). The ACF amplitude scales with the inverse of the number of molecules per pixel, thus, small surface densities as well as clustering of molecules yielded a pronounced increase of the ACF amplitude (Rossboth et al., 2018; Sengupta et al., 2011). ACF analysis of the recorded STED images of fixed activated BMDCs (**Supp. Fig. 9B)** and B-cells (**Supp. Fig. 10)** decorated with 14.4.4 scF_V_-abSTAR635P did not reveal any difference in the ACF curves when compared to analyses of simulated random pMHCII-distributions.

In summary, the use of two independent super-resolution methodologies and two autonomous cluster analysis tools failed to provide any evidence for a clustered surface distribution of I-E^k^ on the surface of living as well as PFA-fixed activated BMDCs and B-cells.

### Surface resident pMHCIIs are to 99% mobile with 60-70% displaying fast diffusion immediately after entering the cell surface

The lateral mobility of pMHCII may influence T-cell antigen detection as it affects (i) the speed with which antigens become distributed on the APC surface, (ii) the frequency of serial TCR engagement, (iii) the dynamics of lateral pMHCII interactions, and also (iv) the mechanical forces exerted on pMHCII-engaged TCRs (Gohring et al., 2021) with possible consequences for the ensuing TCR-proximal signaling response (Hsu et al., 2012). To measure the lateral mobility of I-E^k^ by single molecule tracking, we decorated I-E^k^ molecules in low abundance on non-activated as well as activated BMDCs and B-cells with Alexa Fluor 555-conjugated 14.4.4 scF_V_ (14.4.4 scF_V_-AF555) prior to allowing the cells to spread on ICAM-1-coated glass slides for TIRF imaging at 37 °C (**Fig. 3A, Supp. Fig. 11A-D**). We recorded traces of single I-E^k^ molecules over several hundred milliseconds with a delay time of about ten milliseconds. Trajectories containing two to 50 steps were used to calculate the mean square displacement and diffusion constants for I-E^k^ (**Fig. 3B-C**). Overall, I-E^k^ diffusion was significantly faster on activated BMDCs than on activated B-cells with average diffusion constants of D = 0.30 µm^2^ s^-1^ and D = 0.21 µm^2^ s^-1^, respectively (**Fig. 3D**). Since we witnessed a broad distribution of diffusion constants for individual I-E^k^ traces, we fitted all recorded trajectories to a binary diffusion model (see Material and Methods) and identified a fast (73.1% on activated BMDC, 62.6% on activated B-cells) and a more slowly diffusing fraction as well as a small fraction of immobile I-E^k^ (< 1%). Based on mean square displacement analysis, the rapidly diffusing I-E^k^ molecules appeared to follow free Brownian motion on activated BMDCs (D = 0.364 µm^2^ s^-1^) and activated B-cells (D = 0.263 µm^2^ s^-1^) while the slowly moving I-E^k^ fraction (D = 0.026 µm^2^ s^-1^, activated BMDCs; D = 0.017 µm^2^ s^-1^, activated B-cells, **Fig. 3E**) showed signs of confinement with a radius smaller than 75 nm (Wieser and Schutz, 2008). We arrived at similar I-E^k^ diffusion parameters when comparing the binary data fit performed on individual cells (**Fig. 3F**). Following 24 to 48 hours of LPS-treatment, the fraction of rapidly diffusing I-E^k^ entities increased marginally in B-cells, but not in BMDCs and the mobility of rapidly moving I-E^k^ molecules increased slightly on both activated BMDCs and B-cells (**Supp. Fig. 11A-F**). Furthermore, the average diffusion constant of the slowly moving fraction did not increase after LPS treatment and was similar for BMDC-and B-cell-resident I-E^k^ molecules (**Supp. Fig. 11B and 11D**).

We next visualized the dynamics with which I-E^k^ molecules dispersed at any given time on the APC surface. For this we decorated activated BMDCs with a site-specifically biotinylated 14.4.4 scF_V_ (14.4.4 scF_V_-biotin) for the detection with mSav. The latter had been site-specifically conjugated with abCAGE635, a photo-switchable and photo-stable bright fluorophore (**Fig. 3G** **and Supp. Fig. 6C**). Stained BMDCs and B-cells were seeded on ICAM-1 coated glass slides for imaging at 37 °C in TIRF mode. As shown in **Fig. 3H-I** and consistent with high lateral mobility, we witnessed a rapid dispersion of I-E^k^-associated photo-uncaged fluorescence over the entire surface of BMDCs and B-cells within 150 seconds after setting the UV-pulse to photo-converge a small, selected area.

Extracellular antigens are present at high concentrations in the MIIC or endosomal peptide-loading compartments (Pierre et al., 1997; Turley et al., 2000). Antigen proximity could in principle be spatially and temporally conserved on the cell surface after exosome fusion with the plasma membrane. If true, this may in turn enhance T-cell antigen detection. To test whether pMHCII are locally enriched within the first minutes after entering the cell surface, we loaded I-E^k^ with a traceable peptide. To this end, we exposed non-activated BMDCs (day 6) and activated B-cells (day 2) for 0 to 36 hours to a C-terminally biotinylated MCC peptide 88-103 derivative (MCC-PEG_2_-biotin) in the presence of LPS. We first determined the proportion of surface-resident MCC-PEG_2_-biotin-loaded I-E^k^ molecules by flow cytometry of activated BMDCs (up to 17%) and B-cells (up to 25%) with the use of site-specifically AF647-conjugated monovalent streptavidin (**Fig. 4A-B** **and Supp. Fig. 12A**). We next tracked surface I-E^k^/MCC-PEG_2_-biotin complexes six to twelve hours after addition of the peptide to the APC culture, i.e. 0-12 hours prior to maximal I-E^k^/MCC-PEG_2_-biotin surface expression (**Fig. 4A**). To discriminate newly arriving I-E^k^/MCC-PEG_2_-biotin complexes from all other previously arrived I-E^k^/MCC-PEG_2_-biotin complexes, we first blocked surface-exposed-biotin residues quantitatively using an excess of unlabeled monovalent mSav (500 µg ml^-1^, for more details please refer to Methods section). After removing unbound mSav species as part of a rapid washing step conducted at 4°C to avoid fusion of MHCII-loaded anterograde transport vesicles with the plasma membrane, we allowed newly assembled I-E^k^/MCC-PEG_2_-biotin complexes to arrive at the cell surface for ten minutes at 37 °C, where they were directly conjugated to site-specifically abCAGE635-labeled mSav (or AF555-labeled mSav) and subsequently tracked as single molecules after photo-uncaging by TIRF microscopy at 37 °C (**Fig. 4C**). After pre-blocking and staining at 4 °C, only a small number of I-E^k^/MCC-PEG_2_-biotin complexes were still accessible for staining with abCAGE635-labeled mSav (with an average of 70 trajectories per BMDC and 15 trajectories per B-cell). This number increased around ten-fold after incubating the cells at 37 °C, indicating that more than 90% of the observable I-E^k^/MCC-PEG_2_-biotin complexes had arrived at the cell surface within the recovery time of ten minutes (**Supp. Fig. 12B-C**).

**Figure 4:**
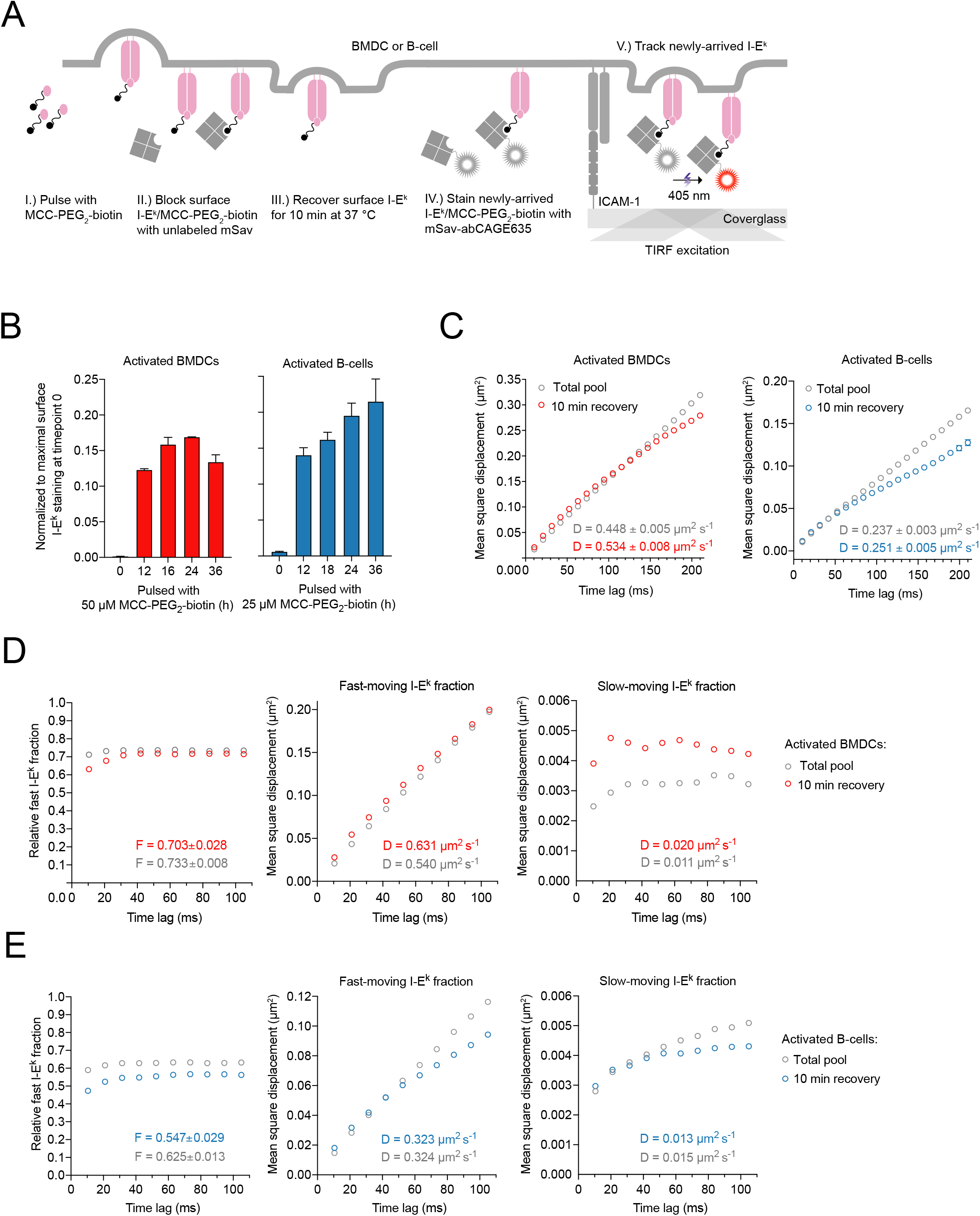
Newly-arriving I-E^k^ molecules are not confined and feature a similar diffusion behavior as the total pool of I-E^k^ molecules. **(A)** Scheme illustrating the experimental procedure applied. (i) Activated BMDCs and B-cells were pulsed with MCC-PEG_2_-biotin. (ii) Resulting I-E^k^/MCC-PEG_2_-biotin molecules on the cell surface were blocked with unlabeled mSAv after the initial peptide pulse (6-12 hours). (iii) After a recovery lasting 10 minutes at 37°C, newly arrived and surface-exposed I-E^k^/MCC-PEG_2_-biotin molecules were stained with mSav-abCAGE635 at room temperature, washed and (iv) directly subjected to single molecule tracking analysis. **(B)** To determine the duration of peptide loading most adequate for tracking experiments (analysis in **D, E**) we quantitated the degree of I-E^k^-peptide-loading on activated BMDCs and B-cells by flow cytometric analysis. Cells were pulsed with MCC-PEG_2_-biotin for up to 36 hours and then stained with either 14.4.4 scF_V_-AF647 (reflecting all surface resident I-E^k^ molecules) or mSav-AF647 (marking I-E^k^ loaded with MCC-PEG_2_-biotin). Ratios were built from MFIs measured for either scF_V_-AF647- or mSav-AF647-decorated cells to determine the proportion of surface-accessible I-E^k^ molecules loaded with MCC-PEG_2_-biotin. Statistics: mean and standard deviation of two technical replicates. **(C)** Activated BMDCs and B-cells pulsed with MCC-PEG_2_-biotin for twelve and six hours, respectively, were stained with mSav-abCAGE635 and seeded on ICAM-1 coated glass slides for imaging in TIRF mode at 37°C. Single I-E^k^/MCC-PEG_2_-biotin molecules were tracked over time with a time lag of 10.5 milliseconds. The graphs demonstrate the mean square displacement of the total pool and newly arrived I-E^k^ molecules on activated BMDCs (10051 trajectories on 14 cells, total pool; 8607 trajectories on 13 cells, 10 min recovery) and activated B-cells (11312 trajectories on 16 cells, total pool; 8974 trajectories on 15 cells, 10 min recovery). Data are representative of one (BMDCs) and two (B-cells) experiments. Statistics: mean and standard deviation. **(D, E)** Fast-moving fraction of newly-arriving and the total pool of I-E^k^ molecules on activated BMDCs and activated B-cells (**E**) were calculated by fitting the recorded I-E^k^ trajectories to a binary diffusion model (**left panel**). Mean square displacement plot of fast-moving I-E^k^ molecules (**middle panel**) and slowly moving I-E^k^ molecules (**right panel**) are plotted for the indicated time lags employed in the tracking experiments. Statistics: mean and standard deviation. F = fraction of fast-moving molecules. D1 = diffusion constant of fast-moving fraction. D2 = diffusion constant of slow-moving fraction. Diffusion constants were calculated by fitting through the first two data points. Fractions were calculated by fitting the first ten data points.

As shown in **Fig. 4C**, newly arriving I-E^k^ molecules exhibited a similar, if not even slightly elevated lateral mobility when compared to all other I-E^k^ molecules on mature BMDCs (D_new_ = 0.534 ± 0.005 µm^2^ s^-1^; D_pool_ = 0.448 ± 0.005 µm^2^ s^-1^) and activated B-cells (D_new_ = 0.237 ± 0.003 µm^2^ s^-1^; D_pool_ = 0.251 ± 0.005 µm^2^ s^-1^). The proportion of newly-arrived rapidly moving I-E^k^ molecules remained unchanged on activated BMDCs (**Fig. 4D** **and Supp. Fig. 12D)** and was marginally decreased on activated B-cells (**Fig. 4E** **and Supp. Fig. 12E)**. Our single molecule tracking analysis rendered a scenario, in which newly arriving antigen-loaded MHCIIs conserve their spatial proximity observed in MIIC compartments and anterograde transport vesicles, rather unlikely, at least within a physiologically meaningful timeframe. Still to be considered is however the possibility that such molecules undergo confined diffusion over a prolonged period of time together with other colocalizing antigen-loaded MHCIIs derived from the same vesicle. Yet inconsistent with this notion is the observation that the labeling procedure employed gave exclusively rise to diffraction-limited fluorescence events which featured brightness values matching those of single fluorophores.

### Surface resident pMHCIIs are monomeric

In large part motivated by our above findings we next sought to assess in a definitive manner the possibility that pMHCII form higher order structures on the surface of living APCs. In fact, the observation that some pMHCII variants crystalize as dimeric entities (Brown, 1993) could in principle result from an inherent propensity of pMHCII to assemble into complexes of larger size. If true, such behavior may sensitize T-cell antigen detection, as had been previously proposed by the “TCR-pseudodimer” model, which postulates simultaneous binding of two adjacent TCRs to a pMHCII agonist and a physically linked pMHCII co-agonist (Krogsgaard et al., 2005).

I-E^k^ surface-densities on living activated BMDCs (i.e. median 514 molecule per µm^2^) and activated B-cells (median 413 molecule per µm^2^) are too high to allow for nanoscopic assessment of individual I-E^k^ mono- or multimers via diffraction limited microscopy. To score for the presence of I-E^k^ oligomers on living APCs, we hence resorted to the “Thinning Out Clusters While Conserving the Stoichiometry of Labeling” (TOCCSL) methodology, a live cell-compatible single-molecule fluorescence recovery after photobleaching (FRAP) approach, which affords identification and quantification of laterally mobile higher order entities at high surface densities (Brameshuber et al., 2018; Brameshuber et al., 2010; Moertelmaier et al., 2005). For TOCCSL measurements, we stained activated BMDCs and B-cells with saturating quantities of the 14.4.4 scF_V_-AF647 and then allowed the cells to spread on ICAM-1-coated glass slides for imaging in TIRF mode. To ensure quantitative decoration of cell surface-resident I-E^k^ with the 14.4.4-scF_V_, we conducted TOCCSL experiments at 25 °C (a temperature at which 14.4.4-scF_V_ exhibited a half-life of binding of ∼80 minutes) and only between two and 20 minutes after the cells had landed on the glass slide. As is illustrated in **Fig. 5A**, we subjected cells to a laser-based TOCCSL illumination sequence, which included a high-powered laser pulse (750-1000 ms) to quantitatively ablate the AF647 fluorescence of a spatially defined area, to be succeeded by the acquisition of a control image 100-750 milliseconds after bleaching for verification of complete AF647-photobleaching (**Fig. 5A**, center panel). Following a recovery phase of two to 20s, we recorded in a so-termed TOCCSL image isolated diffraction-limited spots. These originated from the masked area of the cell surface as laterally mobile I-E^k^ entities and had therefore not undergone bleaching (**Fig. 5A**, right panel). To enumerate the number of 14.4.4-scF_V_-AF647-decorated I-E^k^ molecules per diffraction-limited signal, we compared the obtained distribution of AF647-fluorescence brightness values ρ(B) with the brightness values obtained from recorded cells decorated with AF647-conjugated and unconjugated 14.4.4-scF_V_ pre-mixed at a ratio of 15:1 as monomer control (ρ1). A linear combination of multimeric n-mer contributions, ρn, derived from ρ1 and applied to fit the distribution ρ(B) yielded exclusively I-E^k^ monomers (**Fig. 5B**). To determine the overall capacity of the TOCCSL procedure to reveal the existence of diffraction limited entities exhibiting more than one fluorophore, we imaged cells which we had decorated with 14.4.4-scF_V_-AF647-biotin pre-conjugated with mSav-AF647. This regimen gave invariably rise to events associated with two AF647 dyes, and as shown in **Fig. 5B**, 19% and 30% of the events recorded in this fashion on activated BMDCs and activated B-cells, respectively, reflected dimers. The moderate detection efficiency especially in the case of activated BMDCs resulted in all likelihood from fast lateral I-E^k^ diffusion during the application of the TOCCSL bleach pulse, which appeared to have led to a relatively high number of partially ablated dimeric events at the interface of the bleached and the masked area, and which we confirmed in Monte Carlo-based TOCCSL simulations (**Supp. Fig. 13A**). Similarly, crosslinking of I-E^k^-associated biotinylated 14.4.4 scF_V_-AF647 with 125 nM divalent streptavidin revealed a dimer fraction of 20% on activated BMDCs (**Fig. 5C**).

**Figure 5:**
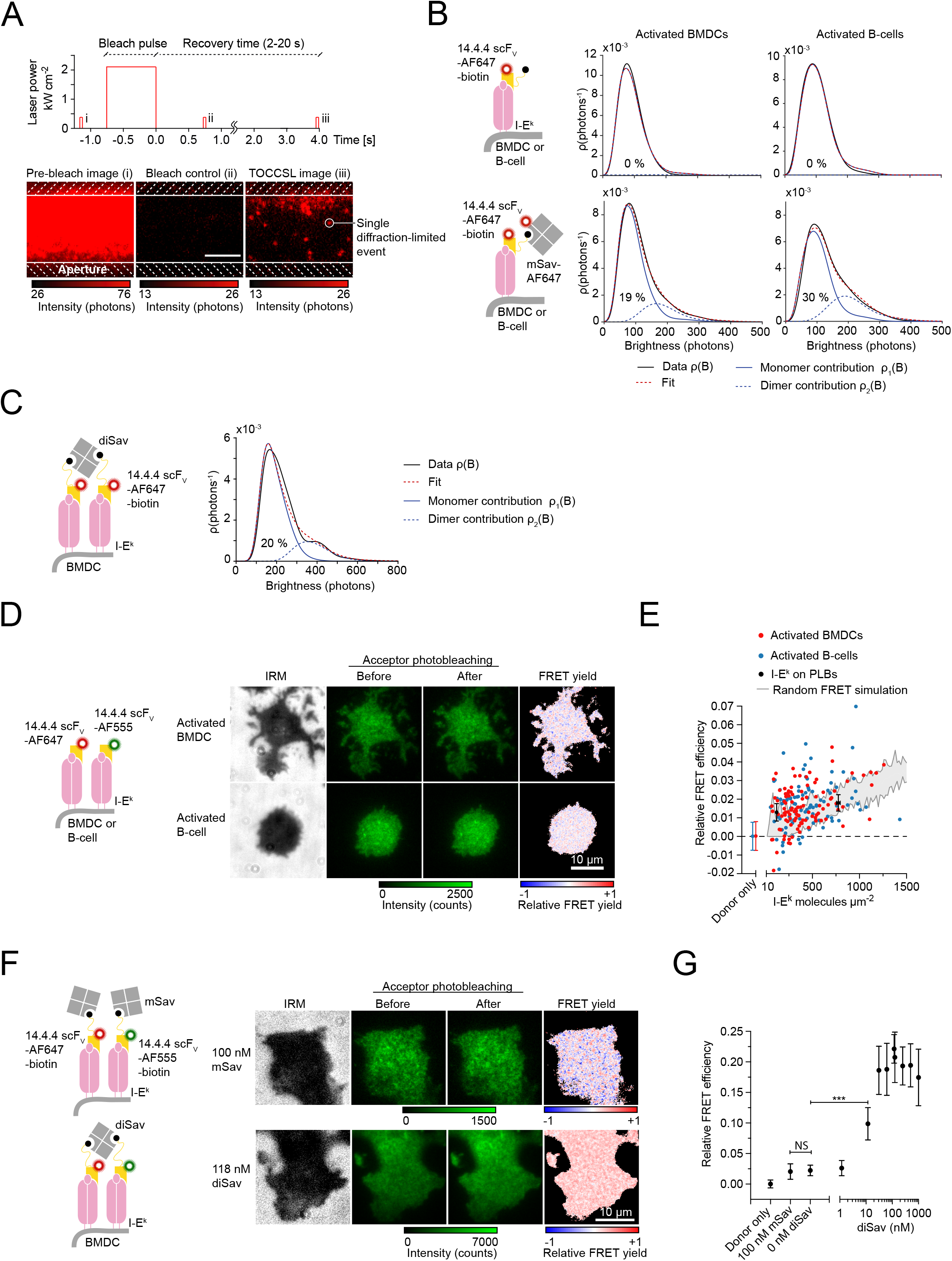
Surface resident pMHCIIs are monomeric. **(A)** Principle of TOCCSL to study the oligomeric state of I-E^k^ on living BMDCs and B-cells. Cells were quantitatively labeled with 14.4.4 scF_V_, placed on an ICAM-1-coated glass slide for imaging and subjected to an illumination sequence of (i) a pre-bleach image followed by a high-powered laser pulse to quantitatively ablate the AF647 fluorescence of a particular cell area (as defined by the slit aperture). (ii) Complete photobleaching of AF647 was verified by a control image 100 milliseconds after the bleach pulse. (iii) Following a recovery phase of 2-10 seconds, we next recorded TOCCSL images to detect diffraction limited single molecule entities resulting from laterally mobile I-E^k^ molecules, which had moved via lateral diffusion from the masked into the unmasked area. Finally, the brightness of individual fluorescence events was determined to assess I-E^k^-complex stoichiometry (shown in **(B, C)**). Scale bar: 5 µm. **(B)** Quantitative brightness analysis revealed a monomeric contribution of recovered I-E^k^ molecules on BMDCs (n = 471 molecules on 59 cells) and B-cells (n = 187 molecules on 52 cells). As monomer control, we stained the cells with a cocktail of 14.4.4 scF_V_-AF647-biotin and 14.4.4 scF_V_-biotin (mixed 1:15) and recorded n = 441 molecules on 50 BMDCs and n = 263 molecules on 57 B-cells. As dimer control we stained BMDCs and B-cells with 14.4.4 scF_V_-AF647-biotin and mSav-AF647 which increased the dimeric contribution of recorded fluorescent events to 19% on activated BMDCs (n = 541 molecules on 64 cells) and 30% on activated B-cells (n = 334 molecules on 61 cells). Data are representative for one (B-cells) and three experiments (BMDCs). **C)** TOCCSL measurement of I-E^k^-bound 14.4.4 scF_V_-AF647-biotin crosslinked with 125 nM divalent streptavidin on activated BMDCs. n = 301 molecules (monomer control) and n = 294 molecules (dimer control) derived from over 50 cells each of one experiment. **(D)** Activated B-cells and BMDCs were stained with a 1:1 premix of the 14.4.4 scF_V_-AF555 and 14.4.4 scF_V_-AF647 and seeded onto ICAM-1-coated glass slides for imaging in TIRF mode to probe molecular proximities of APC-surface expressed pMHCIIs via FRET imaging. FRET donor intensity (green, background subtracted) was monitored before and after FRET DRAAP. FRET efficiencies were calculated on a pixel-by-pixel basis for fully attached membrane regions (see IRM image; scale bar: 10 µm). **(E)** FRET measurements of I-E^k^ on activated BMDCs, B-cells and PLBs. Activated B-cells and BMDCs were stained with a 1:1 premix of the 14.4.4 scF_V_-AF555 and 14.4.4 scF_V_-AF647. PLBs featuring I-E^k^ at indicated densities were decorated with I-E^k^/MCC-AF555 and I-E^k^/MCC-AF647 premixed 1:1. Density-dependent FRET efficiencies estimated from randomized molecular pMHCII encounters through Monte Carlo simulations are shown in grey for FRET donor and acceptors present at a 1:1 ratio. Each dot represents the mean FRET value determined for an individual cell. Error bars represent mean and standard deviation of n=71 BMDCs, n=131 B-cells (donor only) and n = 30-34 PLB positions. Upper and lower boundaries of the grey area represent range for n = 20 simulations per data point. Data were pooled from three (BMDCs) and two (B-cells) experiments. **(F, G)** Activated BMDCs were stained with 14.4.4 scF_V_-AF555-biotin and 14.4.4 scF_V_-AF647-biotin premixed at a 1:1 ratio. Dimerization of biotinylated scF_V_s was induced with divalent streptavidin at increasing concentrations and quantified via FRET donor recovery after FRET acceptor photobleaching. The addition of monovalent streptavidin (mSav) did not crosslink the 14.4.4-scF_V_-biotin molecules. Data were pooled from two independent experiments. Statistics: mean and standard deviation and unpaired two-tailed student’s t-test. P value format: P > 0.05 (ns), P ≤ 0.05 (*), P ≤ 0.01 (**), P ≤ 0.001 (***).

To also account for the less abundant yet more slowly diffusing pMHCIIs, which might have escaped detection by the TOCCSL method, we employed FRET as a readout for spatial proximity between surface-resident pMHCII. To this end we quantitatively decorated I-E^k^ molecules on BMDCs, B-cells and functionalized planar glass-supported lipid bilayers (PLBs) with 14.4.4 scF_V_s, which had been site-specifically conjugated with either AF555 or AF647 serving as FRET donor and acceptor, respectively. For most direct FRET yield quantitation, we recorded FRET donor-recovery after FRET acceptor-bleaching. We anticipated FRET yields to augment with increasing FRET acceptor abundance, yet also had factor in decreasing signal-to-noise ratios with decreasing FRET donor surface densities. We therefore applied in staining procedures AF555- and AF647-conjugated 14.4.4 scF_V_s at molar ratios of 1:1, 1:2, 1:3 and 1:4 with expected FRET detection efficiencies of 66.7%, 80%, 85.7% and 88.9% and signal recovery of 50%, 44.4%, 37.5% and 32%, respectively (**Fig. 5D-G** **and Supp. Fig. 13B-D**). To quantitate the I-E^k^ density on BMDCs and B-cells with and without the use of FRET acceptor probe (14.4.4 scF_V_-AF647), we measured the density of I-E^k^ bound to the FRET donor probe (14.4.4 scF_V_-AF555) for every scenario and related loss of the FRET donor intensity with the gain in FRET acceptor intensity. As shown in **Supp. Fig. 13B,** the use of FRET donor and FRET acceptor probes premixed at a ratio of 1:1 led to a twofold reduction in FRET donor intensity (as determined after FRET acceptor ablation), confirming that the density of all I-E^k^ was twofold higher than the density measured for FRET donor probe-associated I-E^k^.

We next compared FRET efficiencies measured on cells and PLBs via FRET donor recovery after acceptor photobleaching with simulated FRET efficiencies we had calculated from monomeric and randomly distributed molecules present at comparable densities. As shown in **Fig. 5D-E** **and Supp. Fig. 13C-D,** we failed to observe in our cell-based measurements significant deviations from FRET efficiencies as expected from random molecular collisions. Yet validating the use of our FRET-based approach to score for higher order I-E^k^ entities, we witnessed a considerable increase in FRET efficiency after cross-linking I-E^k^-bound biotinylated 14.4.4 scF_V_ (14.4.4 scF_V_-AF555-biotin and 14.4.4 scF_V_-AF647-biotin premixed in a 1:1 ratio) on activated BMDCs via divalent streptavidin (**Fig. 5F-G**).

In summary, both TOCCSL- and FRET-based analyses rendered the existence of mobile and immobile I-E^k^ higher order structures, at least at frequencies that would be relevant for T-cell detection, highly unlikely.

### TCR interactions by endogenous bystander-pMHCIIs are indistinguishable from random molecular collisions

We conjectured that, if they exist, co-agonist pMHCIIs engage TCRs within the immunological synapse as a means to sensitize T-cells for rare antigen. To test this, we quantitated synaptic TCR-engagement of previously reported co-agonist pMHCs with the use of a FRET-based imaging approach previously employed to deduce the molecular dynamics of TCR-pMHC engagement *in situ* at the ensemble and single molecule level (Huppa et al., 2010). For this we confronted primary TCR-transgenic T-cells with protein-functionalized PLBs, which served as surrogate APCs and recapitulated scenarios of free diffusion and random distribution of pMHCs, in line with our observations concerning living APCs described above. Of note, the use of PLBs did not only provide full control over the identity and quantity of synaptic T-cell binding partners, but was also compatible with the use of TIRF microscopy, which afforded considerably improved signal to noise ratios when recording individual synaptic TCR-pMHCII interactions via single molecule FRET (Huppa et al., 2010).

For FRET measurements we first decorated 5c.c7 TCR-transgenic T-cells with a site-specifically AF555-conjugated TCRβ-reactive H57-scF_V_ serving as FRET donor and confronted them with PLBs functionalized with the costimulatory molecule B7-1, the adhesion molecule ICAM-1 and I-E^k^ complexed with either the agonist moth cytochrome C peptide 88-103 (MCC), MCC-derivatives or endogenous peptides for T-cell recognition (**Fig. 6A**, first panel). Prior to MHC-loading, presented peptides had been C-terminally conjugated with AF647 to act as FRET acceptor (for details see to Material and Methods). With an AF555/AF647-Förster radius of ∼5.1 nm, the inter-dye distance giving rise to half-maximal energy transfer, we reasoned that our labeling procedure resulted in measurable FRET only when TCR and pMHCII were engaged with one another (**Fig. 6A**, second panel). Given the calculated inter-dye distance of approximately 4.1 nm within the 5c.c7 TCR-I-E^k^/peptide complex (**Fig. 6A**), we expected TCR-pMHC engagement to result in a FRET yield of about 78% (**Fig. 6B, Supp. Fig. 14A-B**), which would be sufficiently high to visualize synaptic TCR-pMHC interactions and quantitate their lifetime at the single molecule level (**Supp. Fig. 14A-C)**. In fact, ablation of AF647 gave indeed rise to an increase AF555 FRET donor fluorescence as a direct measure for the FRET yield (**Fig. 6B** and **Supp. Fig. 14B**).The latter is proportional to the TCR occupancy, the proportion of pMHC within the synapse that are bound to TCRs (**Supp. Fig. 14D-H** and (Axmann et al., 2015)).

**Figure 6:**
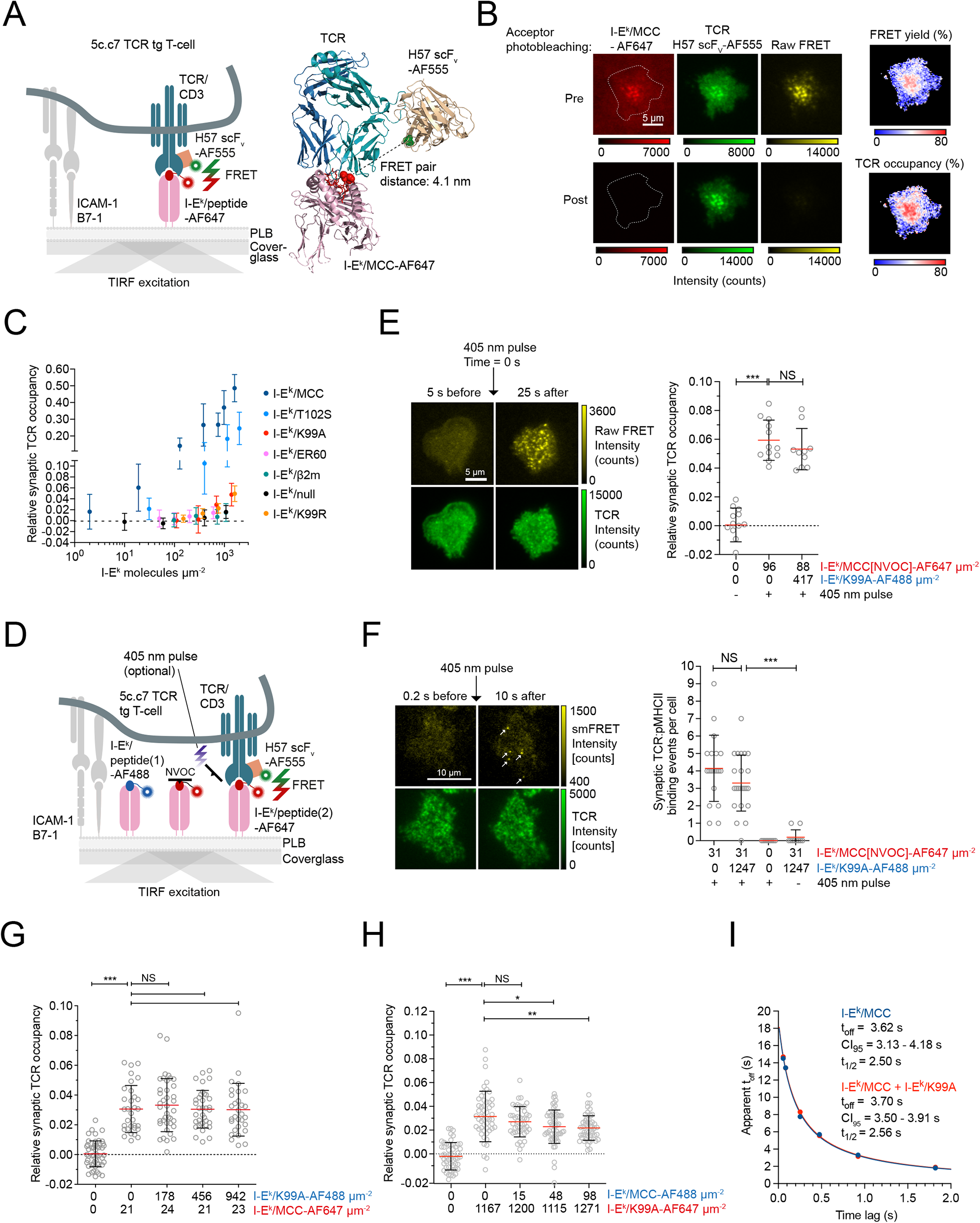
TCR-agonist binding is not affected by the presence of the reported co-agonist I-E^k^/K99A. **(A)** Scheme of a FRET-based assay to visualize TCR-pMHCII interactions *in situ*. **(B)** FRET yields resulting from 5c.c7 TCR-transgenic T-cells stained with H57 scF_V_-AF555 interacting with PLBs featuring I-E^k^/MCC-AF647 as determined via FRET DRAAP and FRET-sensitized emission. The FRET yield is directly proportional to the TCR occupancy, the ratio of bound TCRs to total TCRs. Scale bar: 5 µm. **(C)** Synaptic TCR-ligand engagement was quantified at 25°C via FRET observed for H57 scF_V_-AF555-decorated 5c.c7 TCR-transgenic T-cells engaging PLBs featuring agonist (MCC), weak agonist (T102S), antagonist (K99R), potential co-agonist (K99A, *β*2m, ER60) and non-binding (null) pMHCIIs displayed at increasing densities. ICAM-1 and B7-1 were present at densities of ∼100 molecules per µm^2^. The TCR occupancy of the entire synapse was calculated according to TCR occupancy = E_FRET_ x1.21. Data represent mean and standard deviation of at least 30 T-cell synapses recorded between two and 12 minutes after establishing PLB contact. Statistics: mean and standard deviation. **(D)** Scheme of a FRET-based assay to analyze I-E^k^/MCC-TCR-binding events in the presence or absence of bystander-pMHCIIs with and without the use of a caged peptide (MCC[NVOC]). **(E)** Relative synaptic TCR occupancy of T-cells approaching functionalized PLBs displaying the agonist I-E^k^/MCC[NVOC]-AF647 together with I-E^k^/K99A-AF488 as well as ICAM-1 and B7-1 (both 100 molecules µm^-2^). Binding of the TCR to I-E^k^/MCC-AF647 was quantitated 25 seconds after photo-uncaging via FRET donor recovery after acceptor photobleaching. Scale bars: 10 µm. Statistics: mean and standard deviation and unpaired two-tailed student’s t test. P value format: P > 0.05 (ns), P ≤ 0.05 (*), P ≤ 0.01 (**), P ≤ 0.001 (***). Each grey circle represents a T-cell synapse. This data represents one experiment. **(F)** Single molecule FRET analysis of T-cells approaching PLBs functionalized with the photoactivatable agonist I-E^k^/MCC[NVOC]-AF647 in low abundance and I-E^k^/K99A-AF488 in high abundance together with ICAM-1 and B7-1 (100 µm^-1^). Ten seconds after the uncaging pulse, single molecule FRET events were recorded and counted as individual TCR-I-E^k^/MCC binding events (white arrow) if they had disappeared within one frame during recording. Scale bar: 10 µm. Statistics: mean, standard deviation and unpaired two-tailed student’s t test. This data represents two independent experiments. P value format: P > 0.05 (ns), P ≤ 0.05 (*), P ≤ 0.01 (**), P ≤ 0.001 (***). **(G)** Synaptic TCR occupancy of T-cells engaging I-E^k^/MCC-AF647 in the presence of increasing densities of I-E^k^/K99A-AF488. Statistics: mean, standard deviation and unpaired two-tailed student’s t test. This data is representative for two experiments. P value format: P > 0.05 (ns), P ≤ 0.05 (*), P ≤ 0.01 (**), P ≤ 0.001 (***). **(H)** Synaptic TCR occupancy of T-cells engaging I-E^k^/K99A-AF647 in the presence of increasing I-E^k^/MCC-AF488 densities. Statistics: mean, standard deviation and unpaired two-tailed student’s t test. This data is representative for two experiments. Statistics: mean, standard deviation and unpaired two-tailed student’s t test. P value format: P > 0.05 (ns), P ≤ 0.05 (*), P ≤ 0.01 (**), P ≤ 0.001 (***). **(I)** Synaptic off-rate of I-E^k^/MCC-TCR interactions in the presence or absence of 1060 I-E^k^/K99A-AF488 µm^-2^ as measured via single molecule FRET imaging at 26 °C.

**Fig. 6C** displays TCR occupancies as they had been measured for 5c.c7 TCR transgenic T-cells recognizing I-E^k^ loaded with either the agonist MCC peptide, the weak agonistic T102S, the antagonistic K99R or the reported co-agonist K99A MCC peptide variant (Matsui et al., 1994; Wulfing et al., 1997; Wulfing et al., 2002). We also included measurements involving the recognition of I-E^k^ complexed with the endogenous peptides ER60 and β2m (Ebert et al., 2009). I-E^k^ displaying the triple null MCC mutant, which no longer binds the 5c.c7 TCR (ANERAELIAYLTQAAK, null peptide), was employed as a negative control to determine FRET yields resulting solely from molecular crowding.

Consistent with their stimulatory potency and the TCR occupancies measured, the agonist I-E^k^/MCC and weak agonist I-E^k^/T102S rapidly accumulated underneath T-cells, which was followed by their recruitment to the central supramolecular activation cluster. In contrast, the antagonist I-E^k^/K99R became only slightly enriched, while non-stimulatory I-E^k^/K99A, I-E^k^/ER60, I-E^k^/β2m and I-E^k^/null failed altogether to accumulate in the synapse and to induce the formation of a mature immune synapse (**Supp. Fig. 15A-B**).

As shown in **Fig. 6C**, synaptic TCRs became ligand-engaged already at low densities of I-E^k^/MCC and, to a lesser extent, I-E^k^/ T102S. Expectedly, we observed much reduced synaptic binding of I-E^k^/K99R and especially of I-E^k^/K99A, with median TCR occupancies of ∼5% at high densities (i.e. 1390 I-E^k^/K99A or 1611 I-E^k^/K99R per µm^2^). Of note, we failed to detect any synaptic binding of the endogenous ligands I-E^k^/ER60 and I-E^k^/β2m even when these had been present in numbers comparable to pMHCII expression levels on activated BMDCs and B-cells (median 400-500 I-E^k^ per µm^2^, see **Fig. 1E**). Challenging the notion of any co-agonistic potency associated with I-E^k^/ER60 and I-E^k^/β2m, measured FRET levels could merely be attributed to molecular crowding rather than TCR-binding, as they were indistinguishable from those determined for I-E^k^/null.

### Competition rather than cooperativity among agonist, ‘co-agonist’ and weak agonist ligands defines synaptic pMHCII-TCR binding

To further test the conceptual validity of co-agonism, we assessed whether co-presentation of more weakly binding TCR-ligands promotes or rather suppresses TCR-binding of low abundant agonist ligands.

Among all investigated co-agonist candidates, only I-E^k^/K99A engaged a measurable fraction of synaptic TCRs, at least when present at ∼ 1400 molecules per µm^2^ or higher. To evaluate whether I-E^k^/K99A affected initial TCR-binding to the lowly abundant agonist I-E^k^/MCC, we loaded I-E^k^ with an AF647-conjugated derivative of the MCC peptide, which had been modified at the central lysine residue to contain a (4,5-dimethoxy-2-nitrophenyl)methyl carbonochloridate (NVOC) moiety (DeMond et al., 2006; Huse et al., 2007). As schematically illustrated in **Fig. 6D**, NVOC sterically hinders I-E^k^ binding to the 5c.c7 TCR, unless it becomes ablated on demand through a brief violet (λ = 405 nm) laser pulse to release I-E^k^/MCC for TCR engagement (DeMond et al., 2006). As is demonstrated in **Fig. 6D-F**, T-cells did not engage the provided I-E^k^/MCC[NVOC] until photo-uncaging had occurred.

To assess any influence of co-agonists on the TCR-engagement by agonist ligands, we titrated I-E^k^/K99A-AF488 in high abundance (417 +/-11 molecules per µm^2^) to I-E^k^/MCC[NVOC]-AF647 (92 +/-4 molecules per µm^2^) and quantified 5c.c7 TCR:I-E^k^/MCC engagement by means of FRET donor recovery after acceptor photobleaching 25 seconds after application of the uncaging 405 nm pulse. As can be appreciated from **Fig. 6E**, the presence of I-E^k^/K99A-AF488 did not increase TCR-agonist interactions during synapse formation. To assay earliest TCR-pMHCII binding events, we titrated I-E^k^/K99A-AF488 in high abundance (1247 +/-12 molecules per µm^2^) to I-E^k^/MCC[NVOC]-AF647 (31 +/-0 molecules per µm^2^) and recorded the appearance of single molecule FRET events after photo-uncaging. As shown in **Fig. 6F**, the presence of I-E^k^/K99A-AF488 did also not significantly affect the frequency of initial TCR-agonist binding (i.e. detected single TCR-pMHC binding events via single molecule FRET imaging), rendering scenarios in which bystander-pMHCIIs boost the detection of rare agonist ligands in early synapses highly unlikely.

To assess whether bystander-pMHCIIs influenced synaptic TCR binding of rare agonist pMHCIIs at later stages of synapse formation (i.e. between two and 12 minutes), we measured via FRET TCR-agonist binding on PLBs featuring no or increasing levels of I-E^k^/K99A. As is shown in **Fig. 6G**, synaptic 5c.c7 TCR-I-E^k^/MCC binding was not affected by the presence of up to 942 I-E^k^/K99A µm^-2^, even when agonist ligands were outnumbered by the co-agonist candidates up to 40-fold. Vice versa, 5c.c7 TCR-binding to I-E^k^/K99A-AF647 present in high numbers (1188 +/-65 molecules per µm^2^) dropped appreciably with increasing numbers of co-presented agonist I-E^k^/MCC-AF488 (**Fig. 6H**). This proved also true for synaptic engagement of the weak agonist I-E^k^/T102S-AF647 (present at a density of 1053 +/-55 I-E^k^/T102S), which diminished with increasing densities of co-presented agonist I-E^k^/MCC (ranging from 0 to 202 I-E^k^/MCC per µm^2^, **Supp. Fig. 16A**). In a similar fashion, 5c.c7 TCR-binding to I-E^k^/MCC-AF647 (presented at 38 +/-2 I-E^k^/MCC) dropped substantially with increasing densities of I-E^k^/T102S-AF488 (up to 1235 I-E^k^/T102S per µm^2^, **Supp. Fig. 16B**). We observed a similar trend for the antagonist I-E^k^/K99R, which predictably competed more strongly with agonist binding than I-E^k^/K99A yet to a lesser extent than I-E^k^/T102S (**Supp. Fig. 16C-D**). Of note, high densities of I-E^k^/K99A (1060 I-E^k^/K99A-AF488 µm^-2^) did not affect the synaptic lifetime of agonistic TCR:I-E^k^/MCC interactions as measured via smFRET imaging (**Fig. 6I**).

Taken together, our FRET-based synaptic binding studies render the notion, that TCR-engagement of low affinity pMHCII ligands promote the binding of rare agonist ligands, highly improbable. Instead, and as would be expected from applying the law of mass action, high affinity ligands outcompeted low affinity ligands for TCR binding, especially under conditions where pMHCIIs outnumbered TCRs. Our findings are best explained with a scenario in which pMHCIIs engage TCRs in an autonomous fashion.

### Highly abundant bystander-pMHCIIs do not sensitize T-cells for low levels of agonist pMHCIIs

As shown above we have detected higher order I-E^k^ structures on the surface of activated BMDC and B-cells only after their specific and experimentally defined crosslinking. In addition, we did not detect any influence of bystander pMHCs on the kinetics of synaptic TCR-engagement of rare agonist pMHCs. To be able to arrive at a definitive evaluation of endogenous or low affinity pMHCIIs with regard to their role in sensitized agonist detection, we stimulated T-cells with a defined APC-mimetic biointerface (**Fig. 7A**). The latter involved PLB-embedded and laterally mobile yet structurally rigid DNA origami platforms, which we had engineered to control for pMHCII identities and inter-pMHC distances at the nanoscale level and which allowed for the spatially defined display of I-E^k^ molecules loaded with agonist (MCC) and co-agonist (K99A) peptides (Hellmeier et al., 2021a; Hellmeier et al., 2021b). More specifically and as shown in **Fig. 7A**, we built rectangular DNA origami tiles of 100 x 70 nanometers in size, with either a single or two functionalization sites at ten and 20 nm distance on the top side and attachment sites for cholesterol-DNA at the bottom side for SLB anchorage (for details please see Methods section). To create defined pMHCII homodimers, DNA origami structures were functionalized with biotinylated oligonucleotides, divalent streptavidin and biotinylated pMHCII; for heterodimers, we preassembled DNA-conjugated monovalent streptavidin and biotinylated pMHCIIs prior to DNA origami decoration.

**Figure 7:**
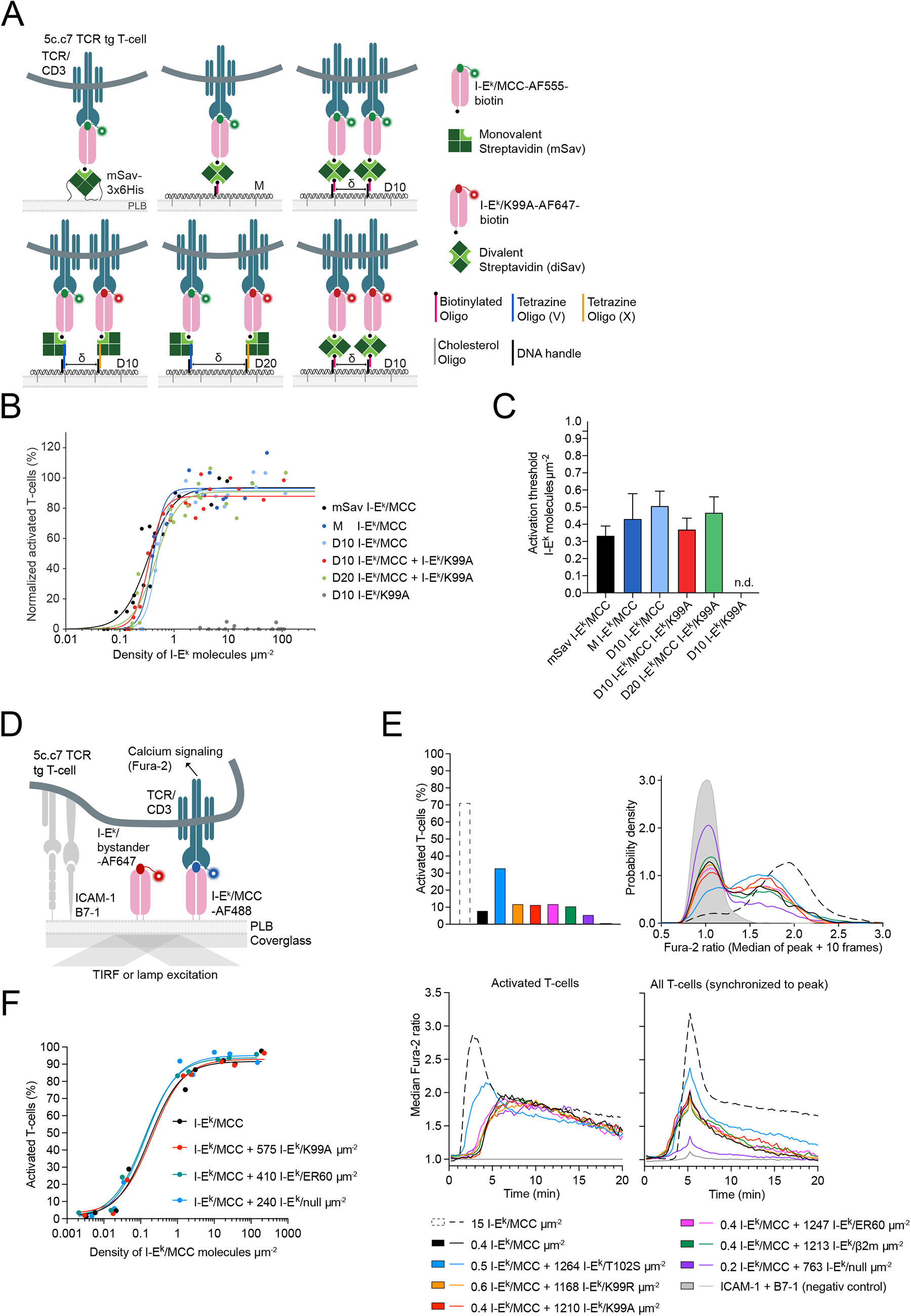
Rare antigenic pMHCIIs are recognized by T-cells independently of highly abundant bystander-pMHCIIs. **(A)** Strategies for the site-specific functionalization of DNA origami structures (70 x 100 nm) with biotinylated I-E^k^/MCC and I-E^k^/K99A molecules to investigate the effect of ligand spacing (D10 = 10 nm, D20 = 20 nm) on T-cell activation. **(B)** Dose-response curves for T-cell activation mediated by I-E^k^/MCC displayed in the absence or presence of the co-agonist I-E^k^/K99A and in the context of the mSav platform or different DNA origami constructs. Each data point corresponds to the percentage of activated T-cells determined in an individual experiment at a specific I-E^k^/MCC density. Data were normalized to activation levels recorded for a positive control (=100%) involving the use of PLBs featuring I-E^k^/MCC at a density of 150 molecules per µm^2^ together with ICAM-1 and B7-1 at 100 molecules per µm^2^. **(C)** Dose-response curves were fitted with **equation (12)** to extract the activation threshold (T_A_, i.e. pMHCII density at half-maximum response, EC50). Statistics: mean and standard deviation (from a bootstrapping analysis). For each dose−response curve, data were derived from at least two independent experiments involving T-cells isolated from at least three different mice. Results of the fit and the significance test of the T_A_ are summarized in **Supp. Table 1B and C**. **(D)** Scheme illustrating the means by which intracellular calcium levels were recorded to assess the impact of highly abundant bystander-pMHCIIs on the recognition of agonist pMHCIIs present in low abundance. **(E)** Calcium response of T-cells in contact with PLBs featuring the agonist I-E^k^/MCC at a density of 0.2-0,6 I-E^k^/MCC µm^-2^, bystander-pMHCIIs at high densities (763 to 1264 I-E^k^ µm^-2^) as well as ICAM-1 and B7-1 (both present with 100 molecules µm^-2^). Shown are percentage of activated T-cells (upper left), probability density plot of Fura-2 ratios of individual T-cells (upper right), mean Fura-2 ratios of activated T-cells over time (lower left), and mean Fura-2 ratios of the entire T-cell population synchronized to onset of signaling (indicated by the highest Fura-2 value within a given track; lower right). Data are representative for more than three independent experiments with the use of antigen-experienced T-cells isolated from more than three different 5c.c7 TCR-transgenic mice **(Supp. Fig. 17B-C)**. **(F)** Calcium response of antigen-experienced 5c.c7 TCR-transgenic T-cells exposed to increasing levels of the agonist I-E^k^/MCC presented together at indicated densities with I-E^k^/K99A, I-E^k^/ER60 or I-E^k^/null.

As shown in **Fig. 7A**, both strategies allowed for site-specific DNA origami attachment of I-E^k^/MCC-AF555 or I-E^k^/K99A-AF647. Biotinylated I-E^k^/MCC-AF555, which had been attached to poly-histidine-tagged monovalent streptavidin, served as control for monomeric pMHCII. Engineered ligand occupancies of DNA origami-based platforms were verified by comparing the brightness of single DNA origami platforms to single fluorescence emitters recorded via TIRF-based microscopy. As shown in **Supp. Table 1A**, we arrived at a functionalization efficiency of ∼ 60% for a single modification site; for two modification sites we detected a mixed population of DNA origami structures featuring one (∼ 20%), two (∼ 50%) and zero ligands (∼10%).

To gauge the impact of homo- or heterodimerization of pMHCIIs on T-cell activation, we loaded 5c.c7 TCR transgenic T-cells with the ratiometric calcium-sensitive dye Fura-2 AM and allowed the cells to engage PLBs displaying functionalized DNA origami platforms at increasing densities and ICAM-1 and B7-1 at densities of ∼100 molecules µm^-2^ (**Fig. 7B**). Ligand densities on PLBs were quantitated by dividing the average fluorescence brightness per area by the average brightness of a single fluorophore molecule. TCR-agonist pMHCII binding resulted in TCR-proximal signaling as witnessed by increasing levels of intracellular calcium acting as second messenger to promote T-cell activation.

For quantitative analysis, we calculated Fura-2 intensity ratios for each T-cell trace and classified the T-cells as activated or non-activated based on a receiver operating curve between a positive control (i.e. > 10 I-E^k^/MCC µm^-2^, ICAM-1 and B7-1 at 100 µm^-2^) and a negative control (ICAM-1 and B7-1 at 100 µm^-2^). For better display of low-level calcium signaling in individual T-cells, we identified the maximum Fura-2 ratio in each cell trace and plotted the median value of the peak Fura-2 ratio (i.e. median of the peak Fura-2 ratio plus ten imaging frames) in a probability density plot (see Material and Methods and **Supp. Fig. 17A**). Non-activated T-cells featured median Fura-2 intensity ratios of 1.0 (after normalization to the median value representing the negative control), whereas oscillatory or activated T-cells exhibited median Fura-2 ratios ranging from 1.3 to 4.0 (**Supp. Fig. 17A**). In **Fig. 7B-C**, we plotted the fraction of activated T-cells as dose response curves as a means to determine activation thresholds, i.e. ligand densities which had given rise to half-maximum responses. To this end, we fitted the data with a four-parameter dose response curve (for more information see Materials and Methods, **equation (12)**; all fit parameters are listed in **Supp. Table 1B**).

When confronted with I-E^k^/MCC-AF555 tethered to monovalent streptavidin, T-cells responded at density of 0.33 antigens per square micron (**Fig. 7C**). Attachment of either a single or two I-E^k^/MCC-AF555 molecules to a DNA origami platform did not lower but rather slightly increased the activation threshold to 0.4 and 0.48 antigens per square micron, respectively. Likewise, confronting T-cells with DNA origami platforms that featured a single I-E^k^/MCC-AF555 as well as a single I-E^k^/K99A-AF647 molecule spatially separated by either ten or 20 nanometers, did not significantly affect activation threshold with values amounting to 34 and 0.46 molecules per square micron, respectively. Of note, T-cells failed to respond to I-E^k^/K99A when displayed in the absence of the agonist ligand I-E^k^/MCC at ∼ 100 molecules µm^-2^

Taken together, pre-organization of two agonist pMHCIIs or one agonist pMHCII and one co-agonist pMHCII did not measurably alter the activation threshold of agonist pMHCIIs in 5c.c7 TCR transgenic T-cells (see **Supp. Table 1C** for significance table). We hence conclude that T-cell activation was independent of ligand spacing. Of note, T-cells failed to respond to I-E^k^/K99A when displayed in the absence of the agonist ligand I-E^k^/MCC at ∼ 100 molecules µm^-2^ (**Fig. 7B**).

We next assessed whether bystander-pMHCIIs in high abundance can trigger low-level calcium signaling in T-cells, which may ultimately affect overall TCR-proximal signaling in the presence of limiting agonist ligands. To test this, we recorded changes in intracellular calcium in 5c.c7 TCR transgenic T-cells in response to increasing densities of I-E^k^ loaded with the agonist peptide MCC, the weak agonist T102S, the co-agonist K99A, the endogenous peptides ER60 and *β*2m, the antagonist K99R as well as the null peptide (**Fig. 7D-E**).

As shown in **Suppl. Fig 17B**, the endogenous ligands I-E^k^/ER60, I-E^k^/*β*2m as well as I-E^k^/null failed to trigger T-cell calcium signaling, even when displayed on the PLB at densities higher than 1000 molecules µm^-2^, as Fura-2 ratio values were comparable to those recorded in the absence of pMHCIIs. The MCC-based altered peptide ligands (APLs) K99A and K99R induced low-level oscillatory calcium signaling patterns even when presented at supraphysiological densities above 1000 molecules µm^-2^ (**Supp. Fig. 17B**).

To assess whether TCR-proximal signaling in response to limiting numbers of agonist ligands was affected by bystander-pMHCIIs co-presented in high abundance, we exposed T-cells to PLBs functionalized with I-E^k^/MCC in low numbers (∼ 1 molecule per µm^2^) together with ICAM-1, B7-1 and high numbers of I-E^k^ loaded with T102S, K99A, K99R, ER60, *β*2m or the non-binding null peptide. As expected, I-E^k^/T102S acted as weak agonist: when present in high abundance, it increased the number of fully activated T-cells as well as the overall calcium signaling capacity (Fura-2 ratio) and accelerated furthermore the onset of signaling. Quite in contrast, endogenous I-E^k^/ER60 and I-E^k^/*β*2m, as well as I-E^k^/K99A, I-E^k^/K99R and I-E^k^/null did not affect calcium response parameters even when present in more than 1000-fold excess (**Fig. 7E** **and Supp. Fig. 17C**). Furthermore, T-cells displayed comparable, if not identical dose response curves towards the agonist I-E^k^/MCC, regardless of the absence or presence of I-E^k^/K99A, I-E^k^/ER60 or I-E^k^/null (**Fig. 7F** **and Supp. Fig. 18A-B**).

In summary, our findings challenge the very concept of pMHCII co-agonism and underscore the stimulatory potency of single, freely diffusing agonist pMHCIIs, as these activated T-cells in a peptide-specific manner and without contributions from endogenous bystander-pMHCIIs.

## DISCUSSION

Advances in our understanding of T-cell antigen recognition and the extent to which it might be sensitized by pMHCII nanoscale organization on professional APCs have been stalled for the last decade, in large part because the readout of most biochemical and imaging approaches employed did not support quantitative conclusions at the molecular level in living cells. We have therefore combined minimally invasive molecular live-cell microscopy with the use of custom-prepared imaging probes to arrive at a meaningful spatiotemporal resolution in the millisecond and nanometer regime without adversely affecting cellular integrity. Following this strategy, we demonstrated that APCs express pMHCIIs at high numbers which become quickly distributed over the entire cell surface as single stimulatory entities as soon as they arrive at the plasma membrane. Highly abundant endogenous pMHCIIs did not enhance TCR-binding and T-cell detection of rare agonist pMHCIIs at early or later stages of synapse formation. Endogenous pMHCIIs need to engage at least a fraction of TCRs in order to induce low-level signaling, which we observed only for MCC-derived APLs when these were presented to T-cells at exceptionally high densities. Together, this leaves us with the conclusion that single agonist pMHCII entities elicit the full response of the scanning T-cell in a truly autonomous fashion. Following this concept, most efficient detection of rare agonist pMHCIIs would hence be best promoted by their rapid redistribution as soon as they arrive on the APC surface.

Central to the robustness of our analysis was the use of a monovalent and site-specifically fluorophore-conjugated (and optionally biotinylated) 14.4.4 mAb-derived scF_V_, which targeted the pMHCII molecule I-E^k^ in a highly specific and stable manner. This allowed for tagging surface-resident I-E^k^ molecules with a single fluorescence emitter and supported quantitative analyses. We can exclude the possibility that we missed surface-resident I-E^k^ in significant numbers as antigen-pulsed APCs preabsorbed with 14.4.4-scF_V_ effectively blocked T-cell activation at 37 °C due to their direct and effective competition with TCRs for pMHCII binding.

Underscoring the validity of employing monovalent and site-specifically labeled scF_V_s for quantitation, we detected 40-times more I-E^k^ molecules per APC (17 times more on B-cells and 40 times more on BMDCs) and also 4-to 5-fold higher molecular densities than previously reported with the use of phycoerythrin (PE)-conjugated full mAbs for cell staining (Unternaehrer et al., 2007). Given their large size, in particular when linked to PE, and their divalent nature, mAbs may be in part sterically hindered or bind instead of a single two pMHCIIs, which is plausible especially when facing exceedingly high epitope densities. Other possibilities include non-quantitative PE-conjugation or mAb-linkage to photo-bleached PE, which together contrast the use of verifiably site-specifically dye-conjugated monomeric scF_V_s, as done in our study.

It was speculated that the high density of pMHCIIs present in MIIC-derived vesicles on their anterograde transport to the cell surface may be preserved at least transiently after vesicle fusion with the plasma membrane to support sensitized T-cell detection (Bosch et al., 2013). Diffusion measurements of pulse-chased I-E^k^ as done in our study render such a scenario unlikely, since 99% of newly arriving pMHCII molecules were mobile and dispersed over the entire APC surface within less than one minute. Single particle tracking revealed furthermore that 60 to 80% of pMHCIIs diffuse rapidly without any evidence of confinement. We examined whether the remaining fraction of more slowly diffusing pMHCIIs (20-40%) were associated with membrane entities such as sphingolipid rafts as had been previously reported (Anderson et al., 2000; Bosch et al., 2013; Hiltbold et al., 2003; Knorr et al., 2009). However, unlike Bosch et al. we failed to recover any surface I-E^k^ in DRMs to an extent that was larger than 1%, when applying a highly quantitative LGC centrifugation- and flowcytometry-based method. This implies that such association is either non-existent on the cell surface or too weak to be conserved when using the non-ionic detergent NP40 employed to discriminate between DRM-resident proteins (e.g. CD44 and CD55) and solubilized membrane proteins (e.g. CD71). Since we failed to co-isolate surface I-E^k^ with DRMs, as would be expected from raft-resident proteins, we consider transient I-E^k^-association with tetraspanins (Zuidscherwoude et al., 2014) or re-internalization (Ma et al., 2012) a likely cause for the observed reduction in lateral mobility.

Of note, living BMDCs and B-cells decorated with monovalent, site-specifically labeled 14.4.4-scF_V_ did not feature in our hands any form of I-E^k^ clustering as was previously reported (Bosch et al., 2013), both above and below the diffraction limit. Decoration of living BMDCs and B-cells with randomly labeled I-E^k^ and I-A^k^-reactive mAbs showed homogenous staining, whereas co-staining with secondary IgG mAbs produced a punctate appearance (**Supp. Fig. 5A-B**). In view of our diffusion and TOCCSL analysis, we consider it likely that the use of divalent mAbs, a temperature shift from 37 °C to 0 °C and PFA fixation had in fact caused the reported dotted appearance of I-E^k^.

Unlike what had been reported earlier, (Bosch et al., 2013; Turley et al., 2000; Zuidscherwoude et al., 2015), neither hsPALM on living cells nor hsPALM and STED microscopy on fixed cells provided in our hands any evidence for clustered pMHCIIs on activated BMDCs and B-cells. Previous studies might have suffered from protein cross-linking as caused by antibodies applied at insufficient concentrations as well as by fixation artefacts impeding an unperturbed assessment of the pMHCII surface distribution. We circumvented these issues altogether by using monomeric and monovalent scF_V_ to avoid receptor cross-linking, even when scF_V_s had been applied at non-saturating conditions, and to allow for non-invasive labeling and imaging strategies. Moreover, we implemented an unbiased cluster analysis protocol which included every single localization and improved the detection of true nanoscale enrichments from false-positive clusters emerging as a result of continuously blinking photo-switchable fluorescent proteins (Baumgart et al., 2016; Platzer et al., 2020). Contrary to our observations, a study involving STED microscopy reported exclusively pMHCII clusters on immature and activated APCs (Zuidscherwoude et al., 2015). It should be noted though that neither (i) the extent to which such reported pMHCII clusters differed in intensity from single molecule emitters nor (ii) the frequencies of their occurrence had been further compared to that of single pMHCIIs. By visually inspecting STED images of I-E^k^ present on activated BMDCs and B-cells, we did not observe a clustered appearance of pMHCIIs. In contrast, especially in areas with low surface density on the APC cell membrane, we detected most I-E^k^ molecules (i.e. 14.4.4 scF_V_-abSTAR635P) as single events which gave rise to the same mean intensity and FWHM values as single fluorescence emitters.

A dimeric arrangement of pMHCIIs was suggested to be a potent mediator of sensitized detection of antigen by T-cells (Krogsgaard et al., 2005), yet without direct confirmation of the existence or the formation of higher-order pMHCII structures within the plasma membrane of living APCs. In fact, despite using all available means we were unable to identify such entities on the surface of activated B-cells and BMDCs. This pertained to their detection via TOCCSL, for which they had to be stable for no longer than 2 to 20 seconds, and via inter-pMHC FRET-based measurements, which as a whole did not provide any evidence for their existence. The latter experiments were performed to account in particular for pMHCIIs diffusing slowly on the cell surface. FRET-based measurements relied on FRET dye combination featuring a Förster radius of 5.1 nm (AF555 and AF647) and site-specifically labeled probes present at ratios of 1:1 to 1:4 for quantitative readout. An earlier study employing FITC and TRITC-conjugated 14.4.4 monoclonal antibodies (Förster radius: 5.5 nm) to reveal homo-association of pMHCII on B lymphoma cells arrived at an average FRET efficiency of 6.5 % without further description of the donor to acceptor ratios applied and the numbers of fluorophores attached to the 14.4.4 mAb (Gombos et al., 2004). In contrast, our quantitative analysis, which included the use of site-specifically conjugated scF_V_ and FRET simulations involving monomeric and randomly distributed FRET pairs, and it failed to provide any evidence for pMHCII homodimerization. Instead, we showed with the use of PLB-anchored pMHCIIs that random encounters of freely diffusing monomeric entities can give rise to FRET values of such magnitude.

With this information we set out to recapitulate antigen recognition by T-cells in contact with PLBs featuring pMHCIIs as well as ICAM-1 and B7-1 at physiological densities. In preceding T-cell imaging studies involving fluorescence and calcium measurements it had been reported that co-agonist pMHCIIs contributed in part to the response to agonist pMHCIIs (Anikeeva et al., 2012; Krogsgaard et al., 2005; Wulfing et al., 2002). Yet other studies featuring similar techniques questioned the role of co-agonists (Ma et al., 2008) or concluded that CD8 co-receptor affinities dictated the co-agonistic effect (Gascoigne, 2008; Hoerter et al., 2013). In this study we found TCR-engagement of all endogenous pMHCIIs with previously attributed co-agonist qualities at the level of random molecular collisions. Only for the altered peptide ligand I-E^k^/K99A (and I-E^k^/K99R, a proposed antagonist), we detected specific TCR-binding, which amounted to fewer than 500 interactions per synapse at any given time when these pMHCII had been present at a density of 1390 molecules per square micron. Assuming that the T-cell synapse covers an area of 100 square microns with a TCR density of 100 TCRs per square micron, this number indicates that on average fewer than 0.4% of available I-E^k^/K99A and only 1% of all synaptic TCRs had been occupied. Not surprisingly, highly abundant I-E^k^/K99A (outnumbering TCRs in our example 14:1) did not alter TCR-binding of rare agonist pMHCIIs in early and mature synapses. In fact, the ensuing TCR-proximal signaling response was unaffected, even when I-E^k^/K99A was co-presented on DNA origami-based platforms in a spatially highly defined fashion with the agonist ligand I-E^k^/MCC. Interestingly, our observations contrast previous findings we had based on the use of DNA-origami tethered anti-TCRβ-scF_V_s as a means of T-cell stimulation (Hellmeier et al., 2021a). Here, we identified distances separating individual stimulatory scF_V_s that were smaller than 20 nm as permissive for T-cell activation, a notion that did not hold up for the recognition of nominal pMHCII antigens. A possible explanation for such divergent behavior may include ligand-dependent differences in stimulatory potency as well as the highly transient nature of TCR-pMHC engagement, which enables single antigens to trigger more than one TCR/CD3 complex in rather close proximity.

It should also be noted that instead of promoting or functionally enhancing rare agonist-TCR interactions, highly abundant bystander-pMHCIIs rather competed with agonist pMHCIIs for available TCRs with increasing TCR-affinities. We also observed that unlike endogenous I-E^k^/ER60 and I-E^k^/*β*2m, the MCC-based APLs I-E^k^/K99A and I-E^k^/K99R promoted low-level signaling when present at supraphysiological pMHCII densities (> 1000 molecules µm^-2^), yet without sensitizing T-cells for agonist pMHCIIs.

In summary, our results leave us with no plausible conclusion other than that single agonist pMHCII entities elicit the full response of the scanning T-cell in a fully autonomous fashion. In view of future improved designs of T-cell-based immunotherapies, this realization places a particular emphasis on a deep subcellular and molecular understanding of effective, faithful and fail-safe TCR-proximal signal amplification, which appears to rely solely on the distinct nature of synaptic TCR-pMHCII interactions and CD4 co-receptor engagement within the context of plasma membrane biophysics and physiology. Our findings imply furthermore that rapid redistribution of newly arriving pMHCIIs rather than clustering underlies highly efficient detection by scanning CD4+ T-cells. It will be highly informative to reveal communalities and differences concerning the organization of MHCI on the surface of target cells with possible consequences for CD8+ T-cell recognition.

## METHODS

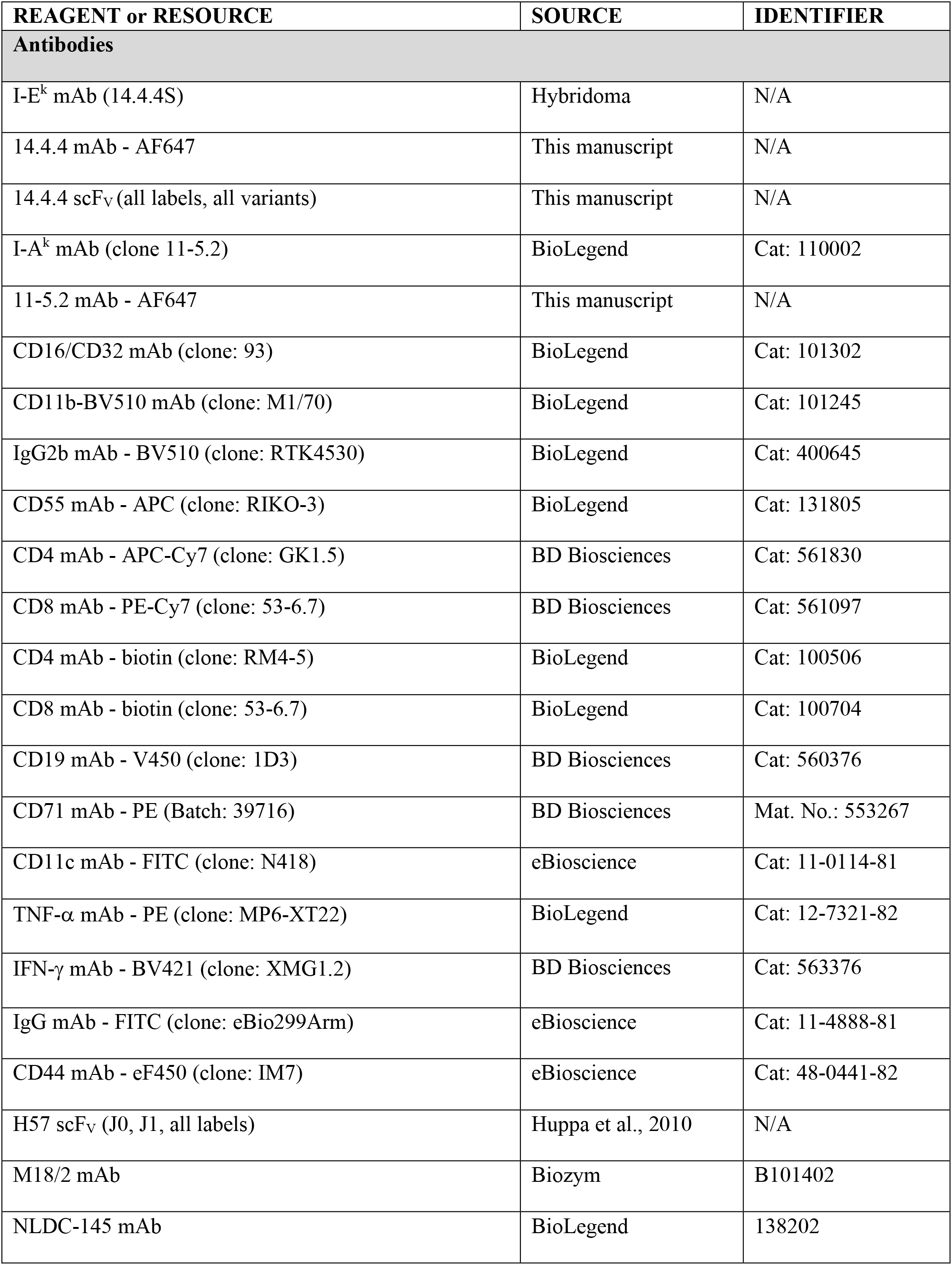
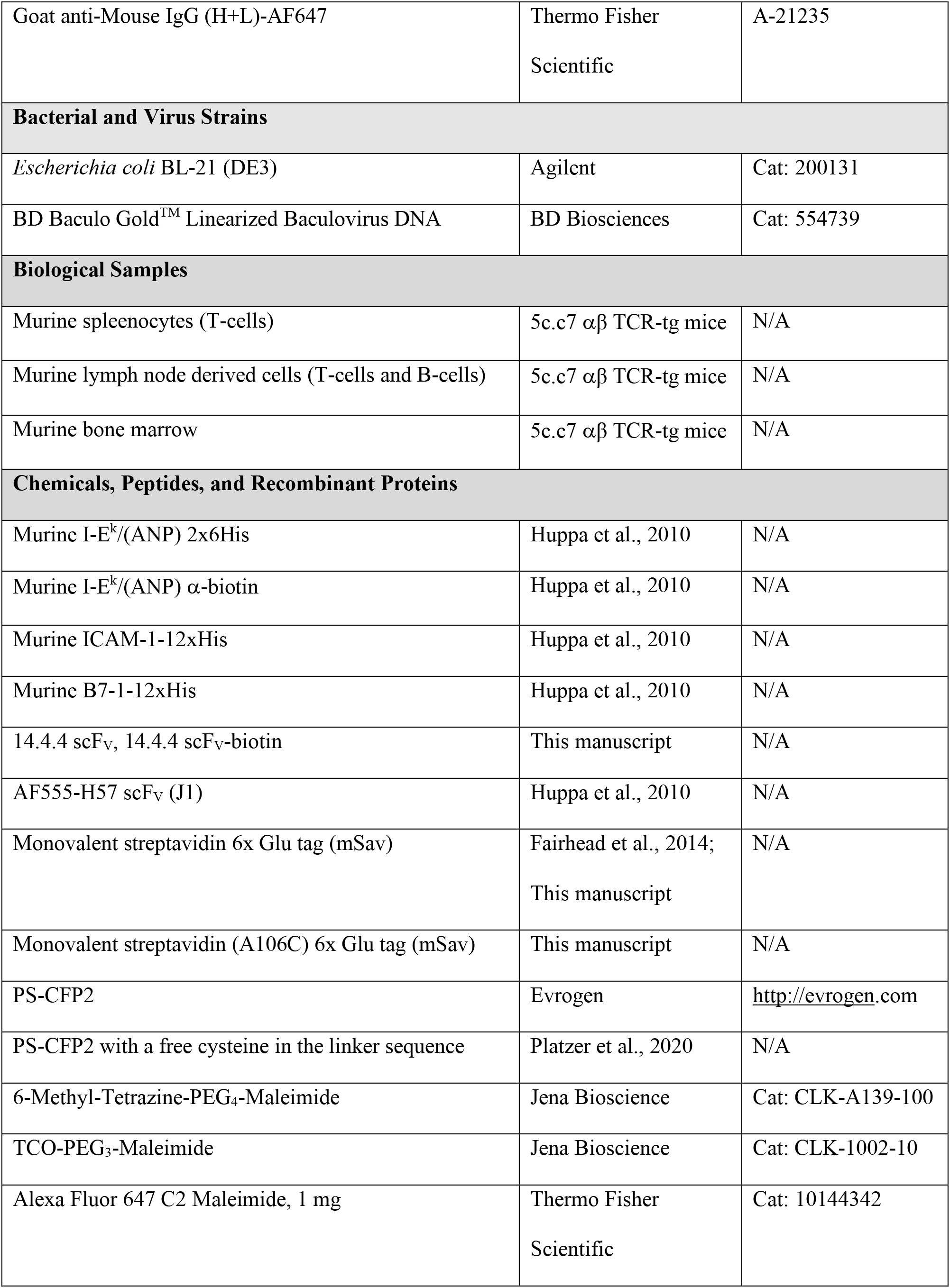
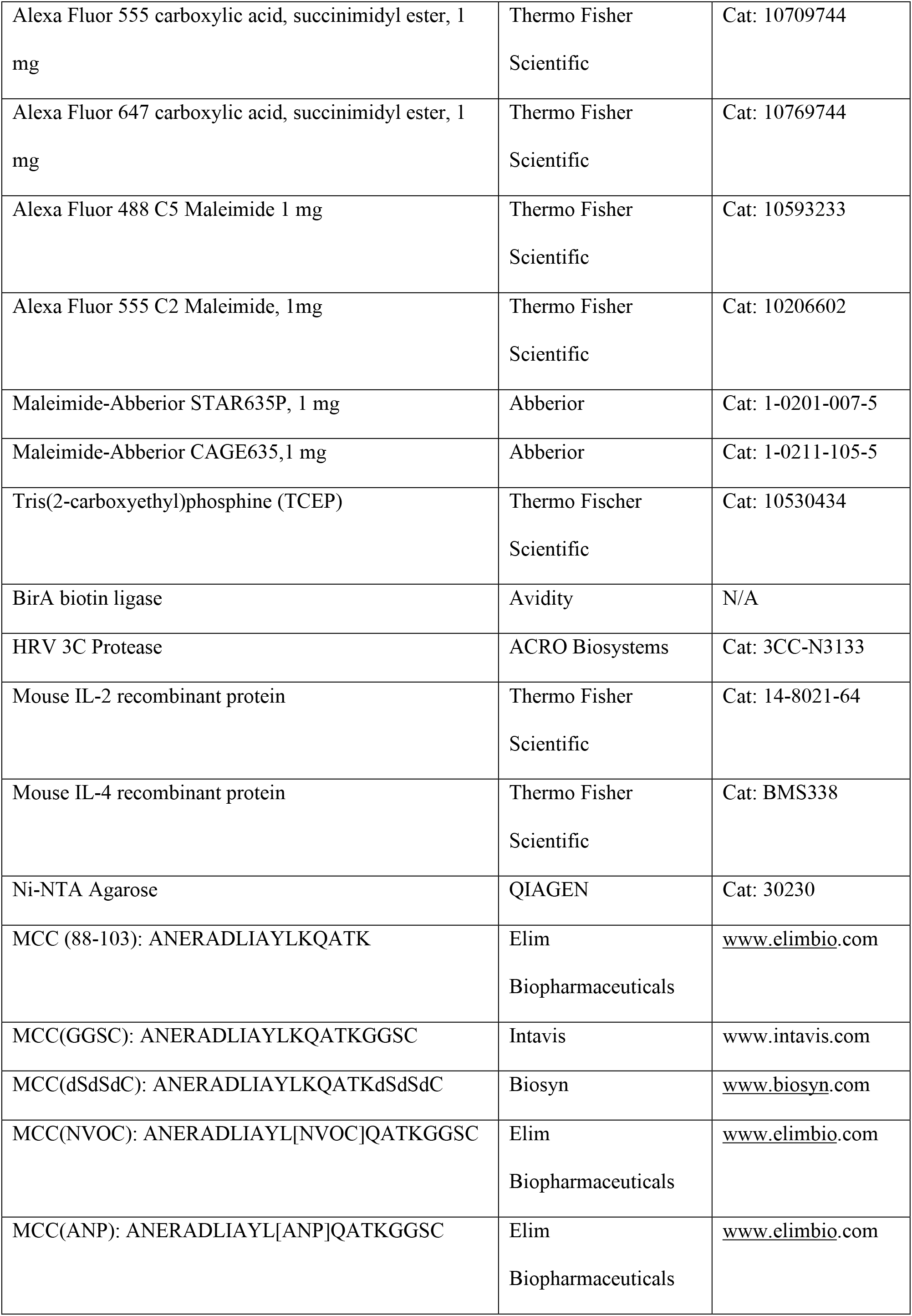
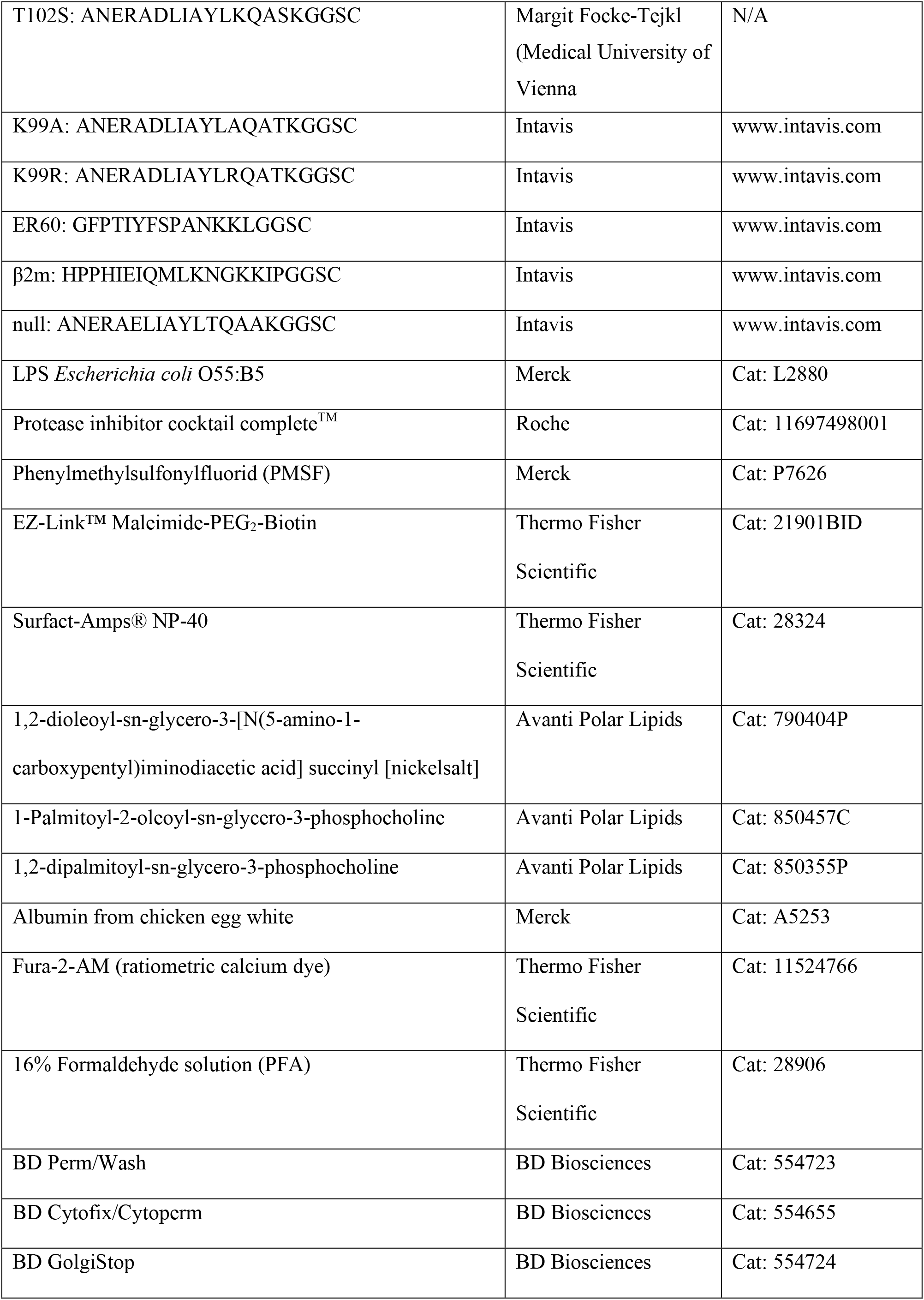
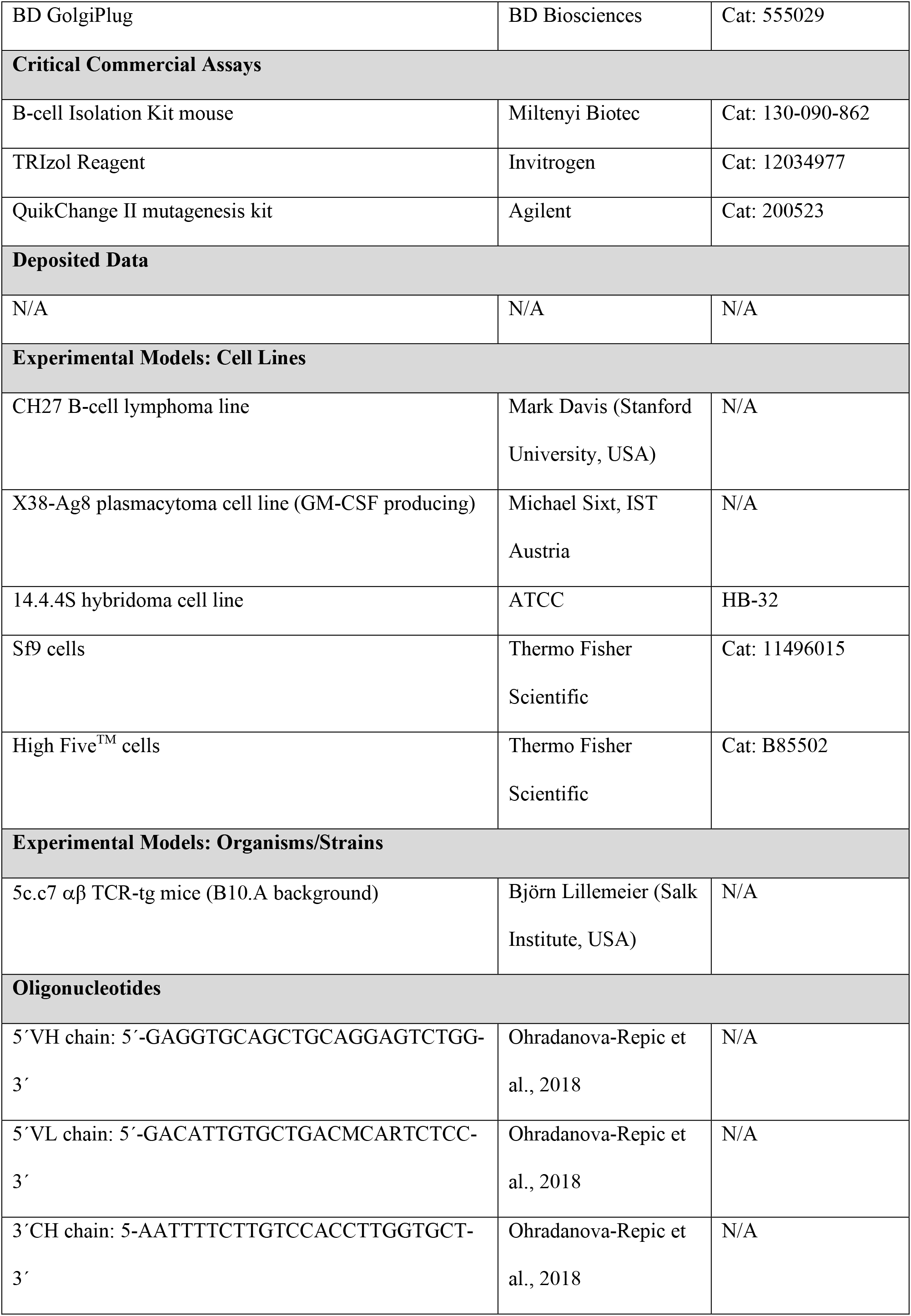
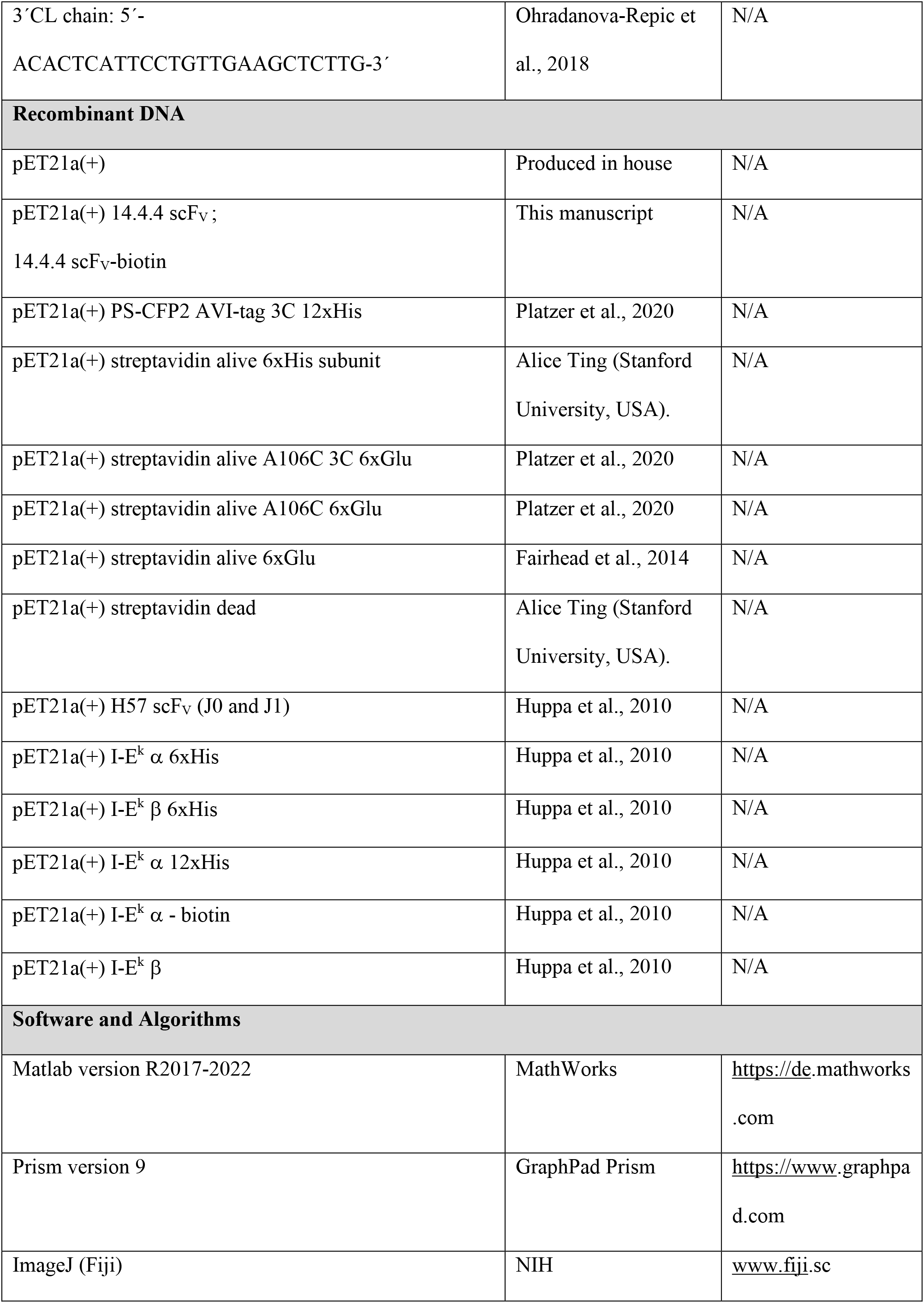
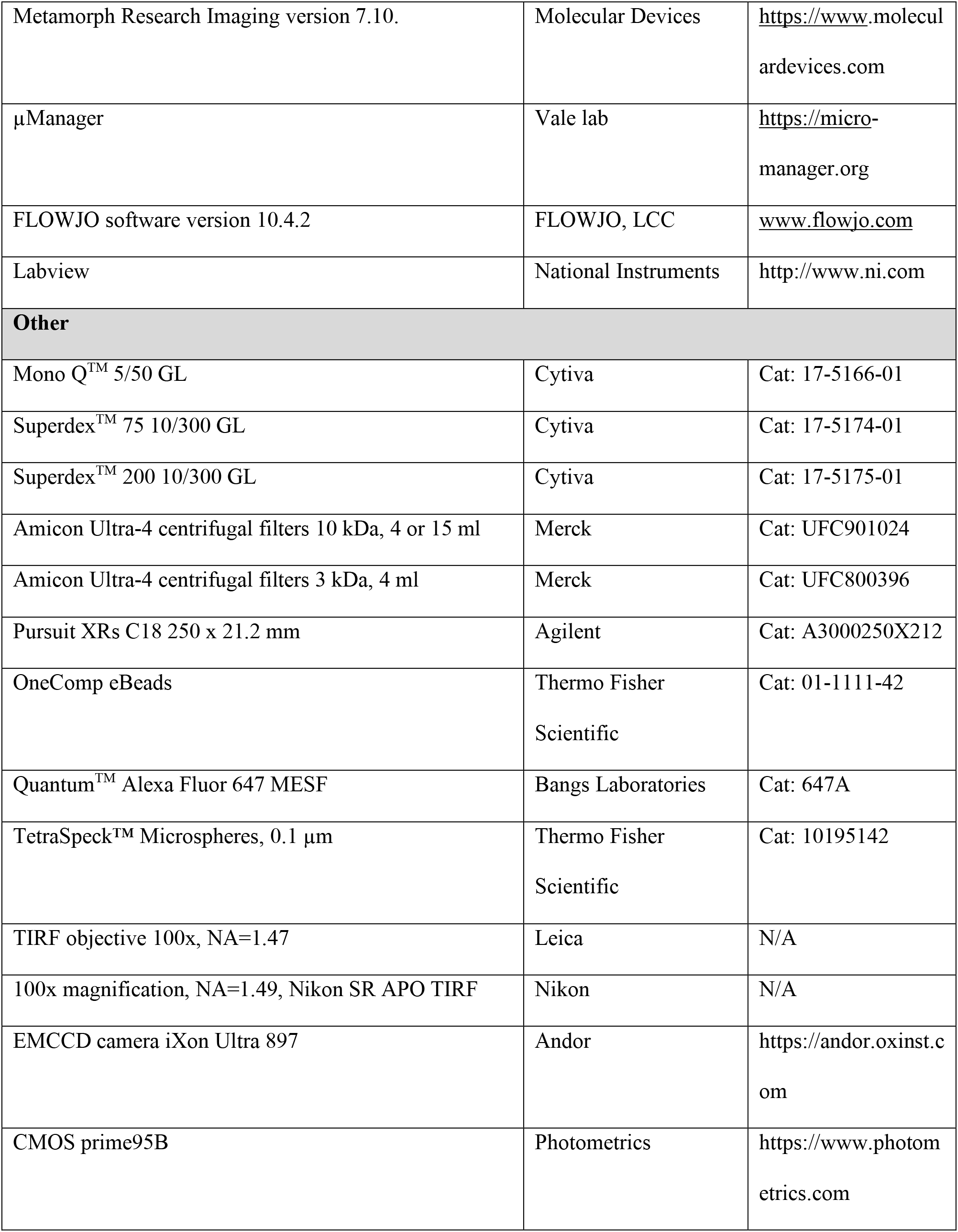

### Animal model and ethical compliance statement

5c.c7 *α*β TCR-transgenic mice bred onto the B10.A background were a kind gift from Michael Dustin (University of Oxford, UK). Mouse breeding and euthanasia were evaluated by the ethics committees of the Medical University of Vienna and approved by the Federal Ministry of Science, Research and Economy, BMWFW (BMWFW-66.009/0378-WF/V/3b/2016). All procedures to isolate lymphocytes, splenocytes and bone marrow from 8-12 weeks old gender-mixed mice were performed in accordance to Austrian law (Federal Ministry for Science and Research, Vienna, Austria), the guidelines of the Federation of Laboratory Animal Science Associations (FELASA), which match those of Animal Research Reporting In Vivo Experiments (ARRIVE), and the guidelines of the ethics committees of the Medical University of Vienna.

### Tissue culture

Splenocytes or lymphocytes of 5c.c7 *α*β TCR-transgenic mice were isolated and pulsed with 0.5 µM C18 reverse-phase HPLC–purified MCC (88-103) peptide (sequence: ANERADL<u>IAYLKQATK</u>, underlined amino acids represent the T-cell epitope; Elim Biopharmaceuticals) and 50 U ml^-1^ IL-2 (eBioscience) for 7 days to arrive at a transgenic T-cell culture (Huppa et al., 2003). T-cells were maintained at 37°C in an atmosphere of 5% CO_2_ in 1640 RPMI media (Life technologies) supplemented with 100 µg ml^-1^ penicillin (Life technologies), 100 µg ml^-1^ streptomycin (Life technologies), 2 mM L-glutamine (Life technologies), 10% FCS (Biowest), 0.1 mM non-essential amino acids (Lonza), 1 mM sodium pyruvate (Life technologies) and 50 µM β-mercaptoethanol (Life technologies). After expansion, T-cells were subjected to Histopaque-1119 (Merck) density gradient centrifugation to remove dead cells. Antigen-experienced T-cells were used for experiments from day eight to ten.

B-cells were isolated from lymph nodes of 5c.c7 *α*β TCR-transgenic mice and enriched using a murine B-cell isolation kit in order to deplete activated B-cells and non-B-cells (B-cell Isolation Kit mouse, Miltenyi Biotec). B-cells were cultured at 37 °C and 5% CO_2_ in 1640 RPMI (Life technologies) supplemented with 10% FCS (Biowest), 100 µg ml^-1^ penicillin, 100 µg ml^-1^ streptomycin (Life technologies), 2 mM L-glutamine (Life technologies) and 50 µM β-mercaptoethanol (Life technologies). B-cells were either used for experiments directly after isolation and enrichment as naïve (non-activated) B-cells or activated with 50 µg ml^-1^ LPS derived from *E. coli* serotype O55:B5 (Merck) for 24 to 48 hours to induce B-cell blast formation.

BMDCs were isolated from 5c.c7 *α*β TCR-transgenic mice and differentiated for 7 days as described (Inaba et al., 2009). BMDCs were maintained at 37 °C and 5% CO_2_ in IMDM (Life technologies) supplemented with 10% FCS (Biowest), 100 µg ml^-1^ penicillin (Life technologies), 100 µg ml^-1^ streptomycin (Life technologies), 2 mM L-glutamine (Life technologies), 50 µM 2-mercaptoethanol (Life technologies), 20 ng ml^-1^ GM-CSF (Biozym) or X38-Ag8 supernatant (1%) and from day 5 on additionally with 10 ng ml^-1^ IL-4 (Life technologies). At day 7, BMDCs were either activated with 100 ng ml^-1^ LPS from *E. coli* serotype O55:B5 (Merck) for 24 to 48 hours to arrive at a mature BMDC culture or alternatively frozen to -80°C in 90% FCS (Biowest) and 10% DMSO (Merck) using a cell freezing container (CoolCell LX, Biozym) and stored in liquid N_2_ for later experiments.

The murine CH27 B-cell lymphoma line was cultured at 37°C and 5% CO_2_ in 1640 RPMI (Life technologies) supplemented with 10% FCS (Biowest), 100 µg ml^-1^ penicillin (Life technologies), 100 µg ml^-1^ streptomycin (Life technologies), 2 mM L-glutamine (Life technologies) and 50 µM 2-mercaptoethanol (Life technologies).

The 14.4.4 mAb producing hybridoma cell line (isotype: mouse IgG2a, kappa) was maintained at 37 °C and 5% CO_2_ in RPMI (Life technologies) supplemented with 5% FCS (Biowest), 100 µg ml^-1^ penicillin (Life technologies), 100 µg ml^-1^ streptomycin (Life technologies) and 2 mM L-glutamine (Life technologies).

The plasmacytoma cell line X38-Ag8 producing GM-CSF was cultivated at 37 °C and 5 % CO_2_ in IMDM (Life technologies) supplemented with 5% FCS (Biowest), 100 µg ml^-1^ penicillin (Life technologies), 100 µg ml^-1^ streptomycin (Life technologies), 2 mM L-glutamine (Life technologies), 50 µM β-mercaptoethanol (Life technologies) and 50 µg ml^-1^ gentamicin (Merck). The supernatant containing GM-CSF was filtered through a 0.45 µm Filtropur S filter (SARSTEDT) and stored at – 80°C.

### Protein expression, refolding and preparation of imaging tools

In order to genetically assemble the 14.4.4 scF_V_, we obtained the sequence information of the 14.4.4 mAb VH and VL chain by isolating total mRNA from a 14.4.4 mAb-producing hybridoma cell line with the use of TRIzol (Invitrogen) followed by reversely transcribing the isolated mRNA into cDNA (First-strand cDNA Kit, Roche).

The 5’primers for the VH chain (5‘-GAGGTGCAGCTGCAGGAGTCTGG-3’) and kappa VL chain (5‘-GACATTGTGCTGACMCARTCTCC-3’) as well as the 3’ primers for the CH (5-AATTTTCTTGTCCACCTTGGTGCT-3’) and kappa CL chain (5‘-ACACTCATTCCTGTTGAAGCTCTTG-3’) were kindly provided by Anna Ohradanova-Repic (Ohradanova-Repic et al., 2018).

We genetically combined the C-terminus of the VH domain and the N-terminus of the VL domain with a [GGGGS]_3_ –linker to obtain a functional scF_V_ transcript, which was cloned into the pet21a(+) expression vector using the restriction enzymes NdeI and HindIII (**Supp. Fig. 19A-B**). To create a target for site-specific modifications using maleimide-based click chemistry, we substituted in the VH domain the serine at position 120 to an unpaired cysteine (14.4.4 scF_V_) via site directed mutagenesis (QuikChange II, Agilent technologies). In addition, we extended the 14.4.4 scF_V_ C-terminally with the BirA biotin ligase (Avidity) recognition site (GLNDIFEAQKIEWHE) to allow for a site-specific attachment of biotin and fluorophores (e.g. 14.4.4 scF_V_-AF647-biotin).

The TCR β-reactive H57 scF_V_ (J0, GenBank: MH045460.1) and the H57 scF_V_, having a free cysteine for site-specific labeling (J1, GenBank: MH045461.1), were produced as described (Brameshuber et al., 2018; Huppa et al., 2010).

All scF_V_ constructs were expressed as inclusion bodies in *Escherichia coli* (BL-21) for 4 hours at 37°C. Bacterial cells processed via ultrasound and insoluble inclusion bodies were extracted with 1% deoxycholic acid (Merck) and 1% Triton X-100 (Merck) in 50 mM Tris pH 8.0, 200 mM NaCl and 2 mM EDTA (Merck) and finally dissolved in 6 M guanidine hydrochloride (Merck). 14.4.4 and H57 scF_V_s were refolded from inclusion bodies *in vitro*, concentrated using an Amicon stirred cell (10 kDa cutoff, Merck) and purified by gel filtration using Superdex 200 (Superdex 200, 10/300 GL, Cytiva) as described (Tsumoto et al., 1998). scF_V_ containing a free cysteine were concentrated in the presence of 50 µM Tris (2-carboxyethyl) phosphine (TCEP, Thermo Fisher Scientific).

After S200 gel filtration, fractions containing monomeric scF_V_ were concentrated with Amicon Ultra-4 centrifugal filters (10 kDa cut off, Merck). 14.4.4 scF_V_ without an unpaired cysteine were randomly conjugated on surface-exposed lysine residues with Alexa Fluor 555 carboxylic acid, succinimidyl ester (Thermo Fisher Scientific) or Alexa Fluor 647 carboxylic acid, succinimidyl ester (Thermo Fisher Scientific). 14.4.4 scF_V_ harboring an unpaired cysteine residue were site-specifically conjugated with Alexa Fluor 647 C2 Maleimide, Alexa Fluor 568 C5 Maleimide, Alexa Fluor 555 C2 Maleimide, Alexa Fluor 488 C5 Maleimide (Thermo Fisher Scientific) or Maleimide-Abberior STAR 635P (Abberior) according to the manufactureŕs instructions and in the presence of 50 µM TCEP, (Thermo Fisher Scientific). The monomeric H57 scF_V_ (J1) was conjugated with Alexa Fluor 555 C2 Maleimide or Alexa Fluor 488 C5 Maleimide according to the same protocol.

To remove excess dye, labeled 14.4.4 scF_V_s were purified by gel filtration using Superdex 75 (Superdex 75, 10/300 GL, Cytiva). Fractions containing monomeric and labeled 14.4.4 scF_V_s were concentrated with Amicon Ultra-4 centrifugal filters (10 kDa cut off, Merck) to arrive at a protein concentration between 0.2 mg ml^-1^ and 1 mg ml^-1^. Protein-to-dye ratios were determined by spectrophotometry at 280 nm and the corresponding absorption maximum of the dyes (647, 635, 568, 555 and 488 nm). Protein-to-dye ratios of randomly conjugated 14.4.4 scF_V_s were between 1 and 2. Protein-to-dye ratios of site-specifically decorated 14.4.4 scF_V_ and 14.4.4 scF_V_-biotin were 0.98 (14.4.4 scF_V_-AF488 and 14.4.4 scF_V_-AF555) and 1.0 (14.4.4 scF_V_-AF647, 14.4.4 scF_V_-AF647-biotin, 14.4.4 scF_V_-AF488-biotin, 14.4.4 scF_V_-AF555-biotin, H57 scF_V_-AF555 and H57 scF_V_-AF488). The 14.4.4 scF_V_-abSTAR635P exhibited a protein to dye ratio of 0.75. Proteins were stored in 1×PBS supplemented with 50% glycerol at -20 °C to -80 °C.

The I-E^k^-specific 14.4.4 mAb was purified from hybridoma supernatant using a protein A/G column (Merck) and gel filtration (Superdex-200 10/300 GL, Cytiva). The 14.4.4 mAb and the I-A^k^-reactive 11-5.2 mAb were randomly labeled via available lysine residues with Alexa Fluor 647 carboxylic acid succinimidyl ester (Thermo Fisher Scientific) according to the manufacturer’s instructions and purified via gel filtration (Superdex-200 10/300 GL, Cytiva). Protein-to-dye ratio of the 14.4.4 mAb and 11-5.2 mAb were 2.1 and 2.4, respectively.

The CD205-reactive mAb NLDC-145 (BioLegend) and the CD18-reactive mAb M18/2 (BioLegend) were randomly conjugated with Alexa Fluor 647 carboxylic acid succinimidyl ester (Thermo Fisher Scientific) and EZ-Link™ NHS-LC-LC-Biotin in a 1 to 2 molar ratio (protein:dye and protein:biotin) in the presence of 0.1 M NaHCO_3_ for 1 hour followed by gel filtration (Superdex-200 10/300 GL, Cytiva). The CD18-reactive mAb M18/2 was additionally purified with monomeric avidin agarose (Pierce) to remove non-biotinylated mAb. The protein-to-dye ratio of the NLDC-145 mAb and M18/2 mAb was 2.1 and 1.3, respectively. The degree of mAb NLDC-145 or mAb M18/2 biotinylation was verified by a monovalent streptavidin-based gel shift assays and SDS-PAGE analysis (**Supp. Fig. 7D**).

mSav was prepared with some modifications as described (Fairhead et al., 2014; Howarth et al., 2006). The pET21a(+) expression vectors encoding either for the “dead” (i.e. biotin non-binding) or the “alive” (i.e. biotin binding) streptavidin subunit were a kind gift from Alice Ting (Stanford University, USA). We substituted the hexa-histidine tag on the “alive” subunit with a cleavable hexa-glutamate tag to allow for purification via cation exchange chromatography (Mono Q 5/50 GL) followed by a recognition site of the 3C protease for optimal removal of the tag (Platzer et al., 2020). To produce mSav for conjugation with maleimide-linked fluorescent dyes or maleimide-tetrazine for orthogonal click chemistry-based protein conjugation, we mutated the alanine residue at position 106 (A106C) for a cysteine residue in the “alive” subunit. The mSav platform mSav-STAR635 was prepared as described (Platzer et al., 2020).

Both, dead and alive streptavidin subunits were expressed in *Escherichia coli* (BL-21) for 4 h at 37°C and refolded from inclusion bodies as described (Fairhead et al., 2014; Howarth et al., 2006; Platzer et al., 2020). After refolding *in vitro* (Howarth and Ting, 2008), the streptavidin tetramer mixture (containing D4, A1D3, A2D2, A3D1 and A4) was concentrated in an ultrafiltration cell (10 kDa cut off, Merck); mSav (A106C) was refolded and concentrated in the presence of 100 µM TCEP. After buffer exchange to 20 mM Tris-HCl, pH 8.0, with Amicon Ultra-4 centrifugal filters (10 kDa cut off, Merck), the mixture of tetramers was purified by anion exchange chromatography (MonoQ 5/50 Cytiva) using a column gradient from 0.1 to 0.4 M NaCl over 90 ml. mSav and mSav (A106C) were eluted with 0.22 M NaCl, concentrated again (Amicon Ultra-4 centrifugal filters, 10 kDa cut off) and further purified via gel filtration (Superdex 200 10/300 Cytiva). mSav was either stored at -80°C in PBS or randomly labeled with fluorescent dyes (e.g. Alexa Fluor 555 carboxylic acid, succinimidyl ester) according to the manufacturer’s instructions. mSav featuring an unpaired cysteine was site-specifically labeled with Maleimide-Abberior STAR 635P, Maleimide-Abberior CAGE 635 or Alexa Fluor 647 C2 Maleimide according to the manufacturer’s instructions. In a last step, dye-conjugated mSav was purified by gel filtration using Superdex 75 (Superdex 75, 10/300 GL, Cytiva) and monomeric fractions were concentrated with Amicon Ultra-4 centrifugal filters (10 kDa cut off, Merck).

Removal of the hexa-glutamate tag was carried out via 3C protease-based cleavage (ACRO Biosystems) for 24 hours at 4 °C followed by subtraction of the GST-tagged 3C protease via glutathione agarose (Thermo Fisher Scientific) and gel filtration (Superdex 200, 10/300 GL, Cytiva). Protein to dye ratios of the randomly dye-conjugated mSav was 1.5 (mSav-AF555); for the site-specifically decorated mSav 1.0 (mSav-AF647). Proteins were stored at -20 °C in 1x PBS supplemented with 50% glycerol.

Divalent streptavidin (diSav) in *trans* configuration (optionally with a 3C cleavage site) was prepared as described (Fairhead et al., 2014). In brief, *trans* diSav was eluted from the MonoQ with ∼ 0.25 M NaCl, purified via gel filtration (Superdex 200 10/300 GL Cytiva), concentrated with Amicon Ultra-4 centrifugal filters (10 kDa cut off), and stored in 1x PBS supplemented with 50% glycerol at – 20 °C.

Monomeric photoswitchable cyan fluorescence protein 2 (PS-CFP2, Evrogen) featuring an unpaired cysteine residue and C-terminally extended with an AVI-tag (GLNDIFEAQKIEWHE) for site-specific biotinylation with the BirA biotin ligase (Avidity) preceded by a 3C protease recognition site (LEVLFQGP) and a 12x histidine (12x His) tag was produced as described (Platzer et al., 2020). To produce a conjugate of mSav and PS-CFP2, we site-specifically decorated PS-CFP2 with 6-Methyl-Tetrazine-PEG_4_Maleimide (Jena Bioscience) and mSav (A106C) with TCO-PEG_3_Maleimide (Jena Bioscience) according to the manufacturer’s instructions. PS-CFP2-tetrazine and mSav-TCO were purified from unreacted crosslinker via gel filtration (Superdex 75 10/300 GL; Cytiva) and concentrated with Amicon Ultra-4 centrifugal filters, (10 kDa cut off) to ∼ 1 mg ml^-1^. To click PS-CFP2-tetrazine to mSav-TCO, we incubated both proteins for 1 day at 4°C in 1x PBS and purified the resulting mSav-cc-PS-CFP2 together with an excess of PS-CFP2-tetrazine via Ni-NTA based affinity chromatography (QIAGEN) and gel filtration (Superdex 200 10/300 GL, Cytiva). Adduct fractions containing mSav-cc-PS-CFP2 were verified by SDS PAGE, concentrated with Amicon Ultra-4 centrifugal filters (10 kDa cut off, Merck) and stored at -20°C in 1x PBS supplemented with 50% glycerol.

For blinking analysis of PS-CFP2, we conjugated the mSav-STAR635 platform with PS-CFP2-biotin, as described in (Platzer et al., 2020).

The murine MHC class II molecule I-E^k^ *α* subunits (C-terminally extended with a BirA biotin ligase recognition site, 6x histidine-tag or 12x histidine-tag) and the *β* subunits (no tag or C-terminally extended with an 6x histidine-tag) were expressed in *E. coli* as inclusion bodies and refolded *in vitro* with a cleavable placeholder peptide (ANERADLIAYL[ANP]QATK) as described (Xie et al., 2012). Specifically, we refolded I-E^k^/ANP 2x6His (or *α*-12His) for attachment to 18:1 DGS-NTA(Ni) and I-E^k^/ANP *α*-biotin for attachment to monovalent or divalent streptavidin. After refolding, we purified I-E^k^/ANP 2x6His via Ni-NTA-based affinity chromatography followed by gel filtration (Superdex 200 10/300 GL; Cytiva). After refolding I-E^k^/ANP *α*-biotin, we purified the protein via affinity chromatography using a custom-made 14.4.4 mAb column (Gohring et al., 2021) followed by gel filtration (Superdex 200 10/300 GL; Cytiva). I-E^k^/ANP *α*-biotin was site-specifically biotinylated using the BirA biotin ligase (Avidity) and purified via Ni-NTA agarose (to remove BirA) and gel filtration (Superdex 200 10/300 GL; Cytiva). Monomeric protein fractions were concentrated to ∼1 mg ml^-1^ (Amicon Ultra-4 centrifugal filters, 10 kDa cutoff), snap-frozen in liquid nitrogen and stored at -80 °C for later use and exchange of the placeholder peptide.

The placeholder peptide can be substituted with any peptide that binds into the peptide-binding cleft of I-E^k^ under acidic conditions (pH adjusted to 5.1 with 0.2 M citric acid buffer) over a period of three days (Xie et al., 2012).

We employed peptides C-terminally extended with a GGSC-linker to render these peptides fit for maleimide-based conjugation to fluorophores. The following peptides were used to exchange the placeholder peptide: MCC (ANERADL<u>IAYLKQATK</u>GGSC, T-cell epitope is underlined), T102S (ANERADL<u>IAYLKQASK</u>GGSC), K99A (ANERADL<u>IAYLAQATK</u>GGSC), K99R (ANERADL<u>IAYLRQATK</u>GGSC), ER60 (GFPT<u>IYFSPANKKL</u>GGSC), β2m (HPPHIEI<u>QMLKNGKKIP</u>GGSC) and null (ANERAEL<u>IAYLTQAAK</u>GGSC). The same loading strategy was used to produce I-E^k^/MCC[NVOC], which contains a UV-cleavable 6-nitroveratryloxycarbonyl (NVOC) attached to the lysine residue in the central position of the MCC peptide (ANERADLIAYLK[NVOC]QATKGGSC) facing the TCR (DeMond et al., 2006).

All peptides were ordered from Intavis as lyophilized product, purified via reversed-phase chromatography (Pursuit XRs C18 column, Agilent) and verified via MALDI-TOF (Bruker) mass spectrometry. For site-specific labeling, peptides were dissolved at a concentration of 1 to 5 mg ml^-1^ in 1x PBS and incubated with TCEP agarose CL4-B (Merck) for 1 hour to reduce paired cysteines. The peptide sample was filtered through a 0.22 µm filter to remove TCEP agarose and incubated with a 1.2-fold excess of maleimide-conjugated dye (Alexa Fluor 647 C2 Maleimide, Alexa Fluor 488 C5 Maleimide or Alexa Fluor 555 C2 Maleimide) for 2 hours in 1 x PBS supplemented with 10 mM HEPES, pH 6.8.

In addition, we produced an MCC peptide derivative (ANERADL<u>IAYLKQATK</u>dSdSdC, d refers to D enantiomer, underlined amino acids are recognized by the T-cell) conjugated with EZ-Link™ Maleimide-PEG_2_-Biotin (Thermo Fisher Scientific) according to the manufactureŕs instructions.

Fluorophore-conjugated (or biotinylated) peptides were purified via reversed phase chromatography on a C18 column (Pursuit XRs C18 column, Agilent) to separate peptide-dye conjugates from unconjugated peptides or dyes. Efficient fluorophore (or biotin) coupling was again verified via MALDI-TOF (Bruker) and nanoHPLC-nanoESI-Linear-Trap-Quadrupole-Orbitrap mass spectrometry (UltiMate 3000 RSLCnano, LTQ Orbitrap Velos, Dionex/Thermo Fisher Scientific). Stoichiometric I-E^k^/MCC[ANP] peptide replacement was gauged via spectrophotometry (280 to 488 or 555 or 640 nm ratio), which was in all cases 90 to 100%.

ICAM-1-12xHis and B7-1-12xHis were expressed in a baculovirus system in Drosophila S2 cells and purified as described (Huppa et al., 2010).

### Antibodies

The I-E^k^-reactive 14.4.4 mAb and 14.4.4 scF_V_, and the TCR *β*-reactive H57 scF_V_ was designed and produced as described above. Antibodies binding to CD4 (APC-Cy7, clone GK1.5), CD8 (PE-Cy7, clone 53-6.7), CD19 (V450, clone: 1D3), CD71 (PE) and IFN-*γ*(BV421, clone: XMG1.2) were purchased from BD Biosciences. Antibodies reactive to CD11c (FITC, clone: N418), Armenian hamster IgG (isotype control-FITC, clone: eBio299Arm) and CD44 (eFluor450, clone: IM7) were purchased from eBioscience. Antibodies reactive to I-A^k^ (clone: 11-5.2), CD16/CD32 (clone: 93), CD11b (BV510, clone: M1/70), rat IgG2b (isotype control-BV510, clone: RTK4530), CD55 (APC, clone RIKO), CD4 (biotin, clone: RM4-5), CD8 (biotin, clone: 53-6.7), CD205 (clone: NLDC-145) and TNF-*α*(PE, clone: MP6-XT22) were purchased from BioLegend. The CD18-reactive mAb M18/2 was purchased from Biozym. The secondary antibody reactive to mouse IgG (anti-Mouse IgG (H+L)-AF647) was purchased from Thermo Fisher Scientific.

### Flow cytometry experiments

For cell surface labeling in general, 0.25 x 10^6^ to 1 x 10^6^ cells were stained with fluorescently-labeled mAbs or scF_V_s for 20 to 30 minutes on ice and washed 2 to 3 times in FACS buffer (1x PBS, 1% BSA, 0.02% NaN_3_). Prior to addition of mAbs, cells were treated with the Fc block CD16/CD32 (20 µg ml^-1^) for 15 minutes. Lymphocytes or splenocytes were stained with CD19 mAb-V450 (1:200), H57 scF_V_-AF488 (10 µg ml^-1^) and modified 14.4.4 scF_V_s (10 – 20 µg ml^-1^). BMDCs were stained with CD11c mAb-FITC (1:200), CD11b mAb-BV510 (1:200), Armenian hamster IgG isotype control mAb-FITC (1:200), rat IgG2b isotype control mAb-BV510 (1:200) and fluorophore-conjugated 14.4.4 scF_V_s (10 µg ml^-1^). CH27 B-cells were stained with fluorophore-conjugated 14.4.4 scF_V_s (titration up to 20 µg ml^-1^). To determine total I-E^k^ expression levels (14.4.4 scF_V_-AF647 staining under saturating conditions) fluorescence associated with AF647-antibody-labeled cells was compared to that of AF647 quantification beads (Quantum^TM^ Alexa Fluor 647 MESF, Bangs Laboratories) serving as reference and measured as a separate sample during the same experiment with the same flow cytometer settings.

For I-E^k^ recovery experiments, B-cells and BMDCs were pulsed with 25 µM and 50 µM, respectively, MCC-PEG_2_-biotin for up to 36 hours, washed and stained with saturating amounts of mSav-AF647 (10 µg ml^-1^) or 14.4.4 scF_V_-AF647 (10 µg ml^-1^).

For intracellular cell-staining, we first labeled cell surface proteins in FACS buffer for 30 minutes before washing with 1x PBS and fixation with BD Cytofix/Cytoperm (BD Biosciences) for 15 minutes. Samples were then washed again with BD Perm/Wash (BD Biosciences) and subsequently stained for intracellular proteins (e.g. mAbs binding to TNF-*α* or IFN-*γ*, both 1:200) in the presence of BD Perm/Wash for 30 min. After another washing step with BD Perm/Wash and FACS buffer, samples were directly subjected to flow cytometric analysis.

Samples were analyzed on the LSR II flow cytometer (BD Biosciences) or the Cytek Aurora (Cytek Biosciences). Data derived from flow cytometry measurements were analyzed with the FlowJo v10 software (BD Biosciences).

### Lysis gradient centrifugation

Association of surface-labeled molecules with DRMs has been determined via lysis gradient centrifugation followed by flow cytometry as described (Schatzlmaier et al., 2015). In detail, 0.2 to 0.5 x 10^6^ primary B-cells and BMDCs were treated with the Fc block CD16/CD32 (20 µg ml^-1^) in RPMI supplemented with 10% FCS (Biowest) on ice for 15 minutes prior to staining with labeled mAbs in a total volume of 50-100 µl. Enriched non-activated and activated B-cells were stained with saturating amounts of CD71 mAb-PE (1:200), CD44 mAb-eF450 (1:200), CD55 mAb – APC (1:200), 14.4.4 mAb-AF647, 11-5.2 mAb – AF647 and 14.4.4 scF_V_ – AF647 (10 µg ml^-1^). BMDCs were stained with CD11c mAb-FITC (1:100) and sorted for CD11c+ cells using a SH800S cell sorter (Sony). CD11c+ (FITC+) cells were further stained with CD71 mAb-PE (1:200), CD44 mAb-F450 (1:200), 14.4.4 mAb-AF647, 11-5.2 mAb-AF647 and 14.4.4 scF_V_-AF647 (10 µg ml^-1^). After staining, cells were washed 2-times in FACS buffer (1x PBS, 1% BSA, 0.02% NaN_3_), and one quarter of the labeled cell samples were kept on ice for intact cell recordings, while three quarters of cells (at least 1 x 10^5^ cells) were subjected to 0.5% NP-40 lysis gradient centrifugation (LGC) to isolate DRM-associated nuclei.

Fluorescence intensities of intact cells and DRM-coated LGC-derived nuclei were analyzed with an LSR II flow cytometer (BD Biosciences). Appropriate single-color staining of cells and OneComp eBeads (Thermo Fisher Scientific) were performed to compensate for spectral overlap. Unstained cells and unstained isolated nuclei were used to evaluate background fluorescence. The FLOWJO software (BD) was employed to determine mean fluorescence intensities (MFIs) from cellular and nuclear single event populations per sample. DRM-association for mAb-stained cell-surface molecules was calculated in percent as [(MFI_nuclei_ – MFI_nuclear background_) / (MFI_intact cells_ – MFI_cellular background_)] x 100.

### BMDC and T-cell co-culture assays

Activated BMDCs were pulsed with 0.008 µM to 1.0 µM MCC (88-103) peptide in the presence of 100 ng ml^-1^ LPS for 1 hour followed by an epitope blocking step with 20 µg ml^-1^ 14.4.4 scF_V_ on ice for 30 minutes. BMDCs were then washed once in T-cell medium (see above) and transferred to a 96-well plate together with 5c.c7 TCR transgenic T-cells at a 1:1 ratio (60 000 cells in total) in the presence or absence of 20 µg ml^-1^ 14.4.4 scF_V_ (total volume 100 µl). The co-culture was maintained in T-cell medium supplemented with BD GolgiStop (1 µl ml^-1^) and BD GolgiPlug (0.7 µl ml^-1^) at 37°C in an atmosphere of 5% CO_2_ for 4 hours prior to intracellular staining and flow cytometric analysis.

### SDS PAGE and silver staining

Protein samples were mixed 3:1 with a 4x loading buffer (250 mM Tris–HCl, 8% SDS, 40% glycerol and 0.04% bromophenol blue, pH 6.8) optionally supplemented with 20 mM dithiothreitol (Merck) to arrive at non-reducing and reducing conditions, respectively, and then subjected to 10-12% SDS-PAGE in running buffer (25 mM Tris–HCl, 192 mM glycine and 0.1% SDS, pH 8.2). Silver staining and colloidal Coomassie G-250 staining was carried out for visualization of the protein bands as described (Chevallet et al., 2006; Dyballa and Metzger, 2009).

### Preparation of functionalized glass-supported lipid bilayers (PLBs)

1,2-dioleoyl-sn-glycero-3-[N(5-amino-1-carboxypentyl)iminodiacetic acid] succinyl [nickelsalt] (Ni-DOGS-NTA, Avanti Polar Lipids) and 1-palmitoyl-2-oleoyl-sn-glycero-3-phosphocholine (POPC, Avanti Polar Lipids) were premixed at a 1:9 molar ratio or a 2:98 molar ratio in chloroform (Merck) to arrive at PLBs harboring Ni-DOGS-NTA at 10% or 2%, vacuum-dried overnight in a desiccator, re-suspended in 10 ml of 1x PBS and sonicated under nitrogen in a water bath sonicator (Q700, QSonica). To produce immobile PLBs we used 1% Ni-DOGS-NTA and 99% 1,2-dipalmitoyl-sn-glycero-3-phosphocholine (DPPC, Avanti Polar Lipids) instead of POPC. Vesicles were subjected to high-speed centrifugation for 4 h at 55,000 g (25°C) and subsequently at 8 h at 75,000 g (4°C) with an ultracentrifuge (Sorvall RC M150GX, Thermo Fisher Scientific) using a fixed angle rotor (S150AT-0128, Thermo Fisher Scientific) to pellet large and non-unilamellar lipid vesicles. Glass slides (24 mm x 50 mm #1 borosilicate, VWR) were either immersed in a 1:1 mixture of 30% hydrogen peroxide (Merck) and concentrated sulfuric acid (Merck) for at least 30 minutes, rinsed with deionized water and air-dried or the surface was treated with a plasma cleaner (Diener ZEPTO) using plasma created from air for ten minutes, and finally glued with picodent twinsil extrahart (Picodent) to the bottom of 8- or 16-well LabTek chambers (Nunc), from which the original glass bottom had been removed. Slides were exposed to a tenfold diluted lipid vesicle suspension (in 1x PBS) for 15 minutes at room temperature (for mobile PLBs) or 60°C (for immobile PLBs) and rinsed with 15-30 ml 1x PBS (room temperature). Subsequently, His-tagged proteins were incubated for 60 to 90 minutes at room temperature and in the dark. After incubation, functionalized PLBs were rinsed twice with 15-30 ml 1x PBS to remove unbound proteins. Shortly before adding the cells to the imaging chamber, 1x PBS was exchanged for imaging buffer containing 1x HBSS (Life technologies) supplemented with either 0.4 mg ml^-1^ Ovalbumin (Merck) or 1% FCS (Biowest) and 2 mM CaCl_2_ and 2 mM MgCl_2_ (Merck).

To prepare ICAM-1 coated glass slides, 10 µg ml^-1^ murine ICAM-1-His_12_ was diluted in 1x PBS and applied to clean chambers for 2 hours at 37°C. ICAM-1 coated glass slides were washed once with 10 ml 1x PBS, which was exchanged for imaging buffer consisting of 1x HBSS (Life technologies) supplemented with 1% FCS (Biowest) and 2 mM CaCl_2_ and 2 mM MgCl_2_ (Merck) prior to adding the cells.

### Preparation of DNA origami

DNA origami structures were assembled in a single folding reaction followed by purification and step-wise functionalization as previously described (Hellmeier et al., 2021a). At sites chosen for ligand attachment, staple strands were elongated at their 3’-end with 21 bases. At sites chosen for cholesterol anchor attachment, staple strands were elongated at either their 3’-end 25 bases, respectively. DNA origami were stored up to 2 weeks at -20°C. DNA origami decorated with a single pMHCII species were functionalized as described in (Hellmeier et al., 2021a). Hetero-dimeric configurations on DNA origami were created by pre-incubating biotinylated pMHCII with a 3x molar excess of DNA-conjugated mSAv (X-strand (K199) – CTTCTGTCTATCTTGGC , V-strand (MCC) – ACATGACACTACTCCAC) for 30 min at 24°C. Unbound proteins were removed by spin column purification (100kDa cut-off, Amicon*®* Ultra centrifugal filters (UFC210024, Merck)). As a last step, DNA-conjugated mSAv bound biotinylated pMHC (K199-MHC, MCC-MHC) were hybridized to elongated staple strands on DNA origami at a 5x molar excess for 60 min at 24°C followed by spin column purification (100kDa cut-off). Functionalized DNA origami platforms were seeded onto SLBs decorated with B7-1 and ICAM-1 at 100µm-2 and were used for experiments at the same day. Functionalization efficiency of DNA origami were determined via two-color colocalization microscopy as previously described (Hellmeier et al., 2021b).

### Microscopy setup

For molecular live cell imaging, we operated setup #1 and setup #2 at the Medical University of Vienna and setup #3 at the Technical University of Vienna. Setup #1 was a Leica DMI 4000B inverted microscope, which was used to image samples in epifluorescence or total internal reflection fluorescence mode (TIRF) with a 100x objective (HCX PL APO 100x, NA=1.47, Leica) and a 20x objective (HCX PL Fluotar 20x, NA=0.5, Leica). As setup #2 we operated an Eclipse Ti-E (Nikon) inverted microscope system that was equipped with a 100x objective (Nikon SR APO TIRF 100x, NA=1.49) and a 20x objective (Nikon S Fluor, NA=0.75).

If not otherwise mentioned, we used diode lasers with the wavelengths 405 nm (LBX-405-180), 488 nm (LBX-488-200), 515 nm (LBX-515-150), 532 nm (LCX-532-500), 561 nm (LCX-561-250) and 640 nm (LCX-640-500) coupled into a polarization-maintaining fiber with the L6Cc laser combiner box (Oxxius) on setup #1 and #2. For some experiments, we also employed diode lasers with the wavelengths 405 nm (OBIS-405-100), 488 nm (OBIS-488-150-LS), 515 nm (iBEAM-SMART-515-S, Toptica), 532 nm (OBIS-532-100-LS), 561 nm (OBIS-561-100-LS) and 637 nm (OBIS-637-140) in a “free-beam” configuration. Laser excitation light was cleaned up with bandpass filters that matched the corresponding wavelengths (Chroma) and focused with a two-lens telescope onto the back focal plane of the objectives. In addition, setup #1 was coupled to a mercury lamp (Leica EL6000) and setup #2 was coupled to a xenon lamp (Lambda LS lamp, Sutter Instrument Company). Fluorescence lamp illumination was realized by using various sets of filters in the excitation path. The fluorescence lamp of setup #1 was equipped with two fast filter wheels with the following filters installed: ET340/26, ET387/11, ET370/36, ET474/27, ET554/23, ET635/18 (Leica). The fluorescence lamp of setup #2 was equipped with one filter wheel (Sutter Instrument Company) with the following filters installed: ET340/26, ET387/11, ET370/36, ET474/27, ET554/23, ET635/18 (Chroma). Setup #1 was further equipped with the beam splitters calcium cube FU2 (Leica), ZT405/488/561/647rpc and ZT405/488/532/647rpc (Chroma) and a custom Notch filter (405, 488, 532 and 642 nm, Chroma) to block the corresponding laser lights in the excitation pathway. Setup #2 was equipped with the beam splitters ZT405/488/561/647rpc, ZT405/488/532/640rpc, ZT405/514/635rpc (Chroma), ET Dualb. “Sedat” CFP/YFP sbxm + HC510/20, and HC Quadband Fura, GFP, Cy3, Cy5 (Semrock). In the emission path, setup #1 was operating a fast emission filter wheel (Leica) equipped with the custom bandpass filters ET455/50, ET525/36, ET605/52 and ET705/72 (Leica). Setup #2 was equipped with a filter wheel (Sutter Instrument Company) featuring the custom bandpass filters ET450/50, ET510/20, ET525/50, ET605/52 and ET700/75 (Chroma).

Emission light was recorded with the EMCCD camera (Andor iXon Ultra 897) or the CMOS prime95B (Photometrics). To program and apply timing protocols and regulate hardware components, both setups were either operated with an 8 channel DAQ-card (National Instruments) or the USB-3114 analog and digital output system (Measurement Computing) controlled by the microscopy automation and image analysis software Metamorph (Molecular Devices). Setup #1 was additionally operated with the Leica software (Leica Suite) and µManager (Open Imaging). All setups were enclosed with a heating chamber (Leica (setup #1), Pecon (setup #2)) and equipped with a heating unit to adjust the temperature within the box.

Setup #3 was equipped with a monochromatic light source (Polychrome V, TILL Photonics) coupled to a Zeiss Axiovert 200M which was equipped with a 10x objective (UPlanFL N 10x, NA = 0.3, Olympus), 1.6x tube lens, and the EMCCD camera Andor iXon Ultra 897. A longpass filter (T400lp, Chroma) and an emission filter were used (510/80ET, Chroma). Calcium imaging was performed with excitation at 340 nm and 380 nm, with illumination times of 50 ms and 10 ms, respectively. The total recording time was 10 min at 1 Hz. Precise temperature control was enabled by an in-house-built incubator equipped with a heating unit.

### Conventional fluorescence microscopy

For conventional fluorescence microscopy analysis, BMDCs and B-cells were stained with saturating amounts of fluorophore-conjugated scF_V_ (e.g. 10 µg ml^-1^ 14.4.4 scF_V_-AF647) or mAbs (14.4.4 mAb-AF647, 11-5.2 mAb-AF647, IgG (H+L)-AF647) for 30 minutes on ice, washed twice with 10 ml 1x HBSS (Life technologies) supplemented with 1% FCS (Biowest), 2 mM CaCl_2_ and 2 mM MgCl_2_ (Merck) prior to seeding the cells onto ICAM-1-coated glass-slides. Cells were treated with the Fc block CD16/CD32 (BioLegend) at 20 µg ml^-1^ for 15 minutes prior to the addition of mAbs. For experiments involving chemical fixation, cell samples were treated with 4% PFA (Thermo Fisher Scientific) in 1x PBS for 15 minutes at room temperature and subsequently washed in 1x HBSS (Life technologies) supplemented with 1% FCS (Biowest), 2 mM CaCl_2_ and 2 mM MgCl_2_ (Merck) to stop the fixation reaction.

Midsections of BMDCs or B-cells were recorded as z-stacks (50 images with 0.3 µm spacing, total depth of z-stack: 15 µm) and deconvolved with the ImageJ plugin iterative deconvolve 3D, which is based on the DAMAS (Deconvolution Approach for the Mapping of Acoustic Sources) algorithm (Dougherty, 2005). The point spread function in x, y and z used to deconvolve the images was measured with single fluorescent beads.

### Quantification of protein densities within cell membranes and PLBs

Rectangular regions of a cell or planar supported lipid bilayer were specified for further calculations. Next, the camera background was subtracted from the intensity of the selected images and divided by the single molecule intensity of the corresponding fluorophore. The intensity of single fluorescence emitters was obtained from cells or planar supported lipid bilayers decorated with single molecules (e.g. 14.4.4 scF_V_-AF647 or I-E^k^/MCC-AF647) and imaged at a power density of 0.1 – 0.4 kW cm^-2^ and 10-20 ms illumination times. To avoid saturation of the camera when imaging highly fluorescent PLBs, we reduced either the power density by a factor of 10 using neutral density filters in the beam path (OD 1.0, Thorlabs) or reduced the linear EM gain of the EMCCD camera by a pre-determined factor resulting in a ten-fold reduction in pixel counts ranging between 2-3. Protein densities were specified as number of molecules per µm^2^.

### High speed photoactivation localization microscopy (hsPALM)

0.1 to 0.5 x10^6^ BMDCs or B-cells were labeled with 20 µg ml^-1^ 14.4.4 scF_V_-AF647-biotin for 30 minutes on ice, washed twice with 1x HBSS (Life technologies) supplemented with 1% FCS (Biowest), 2 mM CaCl_2_ and 2 mM MgCl_2_ (Merck) and afterwards incubated with excess amounts of mSav-cc-PS-CFP2 for 30 minutes on ice. Next, cells were washed twice and either kept on ice or allowed to spread on ICAM-1-coated glass slides for 1 to 5 minutes and were then directly subjected to live PALM imaging. Optionally, cell samples were fixed with 4% PFA (Thermo Fisher Scientific) in 1x PBS for 15 minutes at room temperature, washed, stored at 4°C (to complete fixation) and imaged either immediately after or the next day. PALM images were recorded in TIRF mode at 25°C. We recorded hsPALM image stacks only from adherent cells, a condition that was determined by one red image (637 nm laser excitation (OBIS-637-140), 705/72 emission filter) of the AF647-conjugated 14.4.4 scF_V_-biotin seconds before streaming the hsPALM image stack. With the intention to excite and bleach PS-CFP2 in one single frame (with 2 milliseconds illumination time and 0.3 milliseconds readout time) we used the 488 nm laser line (OBIS-488-150-LS) with a power density of 3.5 kW cm^-2^. After a short pre-bleach stream with 488 nm laser excitation (50 frames), we used continuous 405 nm laser excitation (OBIS-405-100) operated at a very low power density (1-10 W cm^-2^) to photo-convert PS-CFP2 from cyan to green emission and filtered the emission channel with the bandpass filters ET525/36 (Leica) or ET525/50 (Chroma). With the use of the Andor iXon Ultra 897 EMCCD camera, we collected at a frame rate of 500 Hz 3050 image frames (50 x 200 pixel) sufficient for one reconstructed superresolution image in 7 seconds.

We obtained the blinking statistics of PS-CFP2 under the same experimental conditions (PFA fixed or unfixed) and with the same illumination settings and 488 nm power densities following a procedure we had previously published (Platzer et al., 2020). More specifically, we decorated immobile PLBs with spatially well-separated mSav-abSTAR635P in complex with PS-CFP2 molecules which had been site-specifically biotinylated via an AVI-tag. Imaging of hundreds of different PLB positions was conducted at room temperature and in TIRF mode as described (Platzer et al., 2020). To visualize the position of single mSav-STAR635-PS-CFP2 molecules, we used the 637 nm laser line (OBIS-637-140) with a power density of 0.5-1.0 kW cm^-2^ and the emission bandpass filter ET700/75 (Chroma). We recorded 20 images to localize (and photo bleach) the mSav-STAR635 platform prior to streaming 3050 frames to create the hsPALM stack for the PS-CFP2 blinking analysis.

Single molecule blinking analysis of PS-CFP2 conjugated to the mSav-STAR635 platform was carried out as described (Platzer et al., 2020) with the following modifications: The position of mSav-STAR635 molecules was averaged by using the localization merging algorithm implemented in ThunderSTORM (Ovesny et al., 2014) with the following parameters: maximum t_off_ = 20 frames, maximum displacement = 100 nm. A localization cluster threshold was set to 30 to filter out extreme outliers. Data was pooled from two independent experiments.

PALM image analysis via Ripley’s K was performed as described in (Platzer et al., 2020). We first selected for each cell a rectangular region of interest (ROI) with a maximum size specified by the boundaries of the cell. Single molecule PS-CFP2 signals within the ROI were fitted with a Gaussian intensity distribution by maximum likelihood estimation using the ImageJ plug-in ThunderSTORM and filtered for sigma (< 320 nm) and positional accuracy (< 150 nm). We obtained a positional accuracy of median 51 nm for single PS-CFP2 signals with the illumination settings described above (**Supp. Fig. 6F**). We then performed a qualitative Ripley’s K analysis of a region of interest and compared the calculated maxima of the Ripley’s K function of the cell-derived localization map with a simulated localization map, which represented a random distribution of molecules present at the same molecular density and exhibiting the blinking statistics of PS-CFP2 and in selected cases the diffusion parameters of the respective molecule (Platzer et al., 2020).

PALM image analysis via label density variation was carried out as described in (Baumgart et al., 2016). After obtaining the X and Y coordinates for every localization as described above, we assigned a Gauss function with a sigma (σ) of 35 nm (that was cut at 2σ) to every localization within the sample regions. Briefly, overlapping Gauss functions were summed subpixel-wise after applying a threshold (from 0.5 to 4.5; here 1.0) to generate a three-dimensional cluster map (**Fig. 2E**). Next, we calculated the density within the selected cluster regions (*ρ*; localizations per µm^2^) and the relative area of selected cluster regions (*η*; cluster area divided by total area). The pair-wise numbers obtained for each sample were plotted in a ρ/η diagram and compared to a reference curve for randomly distributed localizations (**equation (1)**). The reference curve was determined by simulating randomly distributed molecules under different conditions (variations in blinking statistics and thresholds) and applying a polynomial function that fits every scenario, as described in (Baumgart et al., 2016).

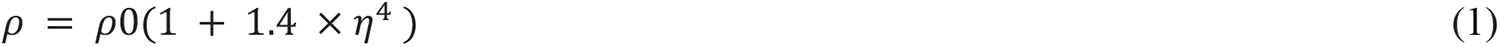

### Single molecule tracking

BMDCs and B-cells were stained with a single molecule dilution of the 14.4.4 scF_V_-AF555 on ice for 30 minutes and washed twice with imaging buffer containing 1x HBSS (Life technologies) supplemented with 1% FCS (Biowest), 2 mM CaCl_2_ and 2 mM MgCl_2_ (Merck). Cells were kept on ice or seeded onto ICAM-1-coated glass slides for imaging at 37°C and in TIRF mode. AF555 was excited with the use of a 532 nm laser (OBIS-532-100-LS), and the emission channel was cleaned up with the bandpass filter ET605/52. We recorded movies of single I-E^k^ (AF555-14.4.4 scF_V_) localizations over 500 frames with a power density of 0.25 – 0.5 kW cm^-2^, an illumination time of ten milliseconds and a total time lag of 10.26 – 10.5 milliseconds between two adjacent images (frame transfer mode).

For I-E^k^ recovery experiments, BMDCs and B-cells were pulsed for twelve and six hours, respectively, with 50 µM MCC-PEG_2_-biotin. We divided each cell sample into two aliquots. The first aliquot represented the total pool of I-E^k^/MCC-PEG_2_-biotin and was kept on ice. The second one was blocked with unlabeled monovalent streptavidin for 30 minutes on ice, washed in imaging buffer and again divided into two aliquots: One was incubated on ice for 10 minutes to serve as blocking control. The second one was incubated for 10 minutes at 37°C to allow for newly synthesized I-E^k^ molecules to arrive at the cell membrane via vesicular transport (recovered I-E^k^ fraction). Finally, all three sample groups representing (1) the total pool of I-E^k^ molecules (2) the blocking control and (3) the fraction of newly arrived I-E^k^ molecules were stained with abCAGE635-conjugated mSav (mSav-abCAGE635) for 5 minutes at room temperature and afterwards washed once with warm 10 ml imaging buffer. Stained BMDCs and B-cells were seeded onto ICAM-1 coated glass slides for immediate imaging at 37°C in TIRF mode. The photoactivatable dye abCAGE635 was activated with low powered 405 nm laser illumination (0.01 – 0.1 W cm^-2^) and excited with 637 nm laser light (OBIS-637-140) with a power density of 0.1 – 0.2 kW cm^-2^. The emission channel was cleaned up with the bandpass filter ET705/72. We recorded movies of single I-E^k^/MCC-PEG_2_-biotin (mSav-abCAGE635) trajectories for up to 1000 frames with an illumination time of 10 milliseconds and a total time lag of 10.26 milliseconds (100 x 100-pixel ROI). B-cells and BMDCs stained with mSav-AF555 were excited with the use of a 532 nm laser (0.25 – 0.5 kW cm^-2^), and the emission channel was cleaned up with the ET605/52 bandpass filter.

Single fluorescent emitters (e.g. I-E^k^ labeled with 14.4.4 scF_V_-AF555) were tracked using a custom written algorithm in Matlab. We calculated the XY localization and intensity of each fluorophore detected within the image stack using the ImageJ plugin ThunderSTORM and filtered for sigma (< 320 nm) and positional accuracy (< 150 nm). Single localizations within the imaging stack were combined to trajectories based on a published algorithm (Schutz et al., 1997). We calculated the mean square displacement (MSD) describing the average of the square displacements between two points of the trajectory with the single molecule position r(t) and the accuracy σ according to **equation (2)**.

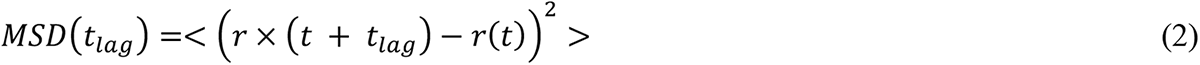

The first two MSD values as a function of t_lag_ were used to calculate the diffusion coefficient (D) for each trajectory according to *MSD* = 4 × *D* × *t_tag_* + 4σ^2^ (Wieser et al., 2007). Multiple mobile fractions were discriminated by analyzing the step-size distributions of square displacements for several time lags (Schutz et al., 1997). By assuming free Brownian motion of one mobile fraction, the cumulative probability for finding a square displacement smaller than r² is given by **equation (3)**.

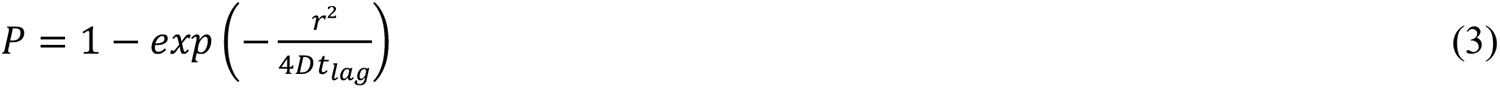

Two different fractions *α and (1-α)* with diffusion coefficients D1 and D2 can be distinguished by fitting the bi-exponential function **(equation (4)**).

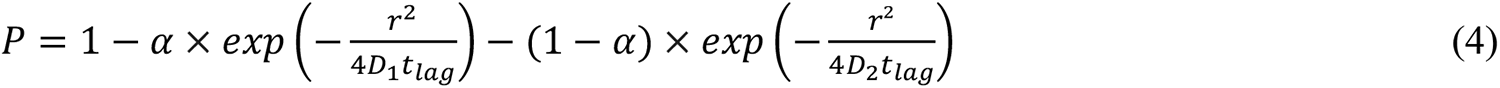

### Thinning out Clusters while Conserving the Stoichiometry of Labeling (TOCCSL)

BMDCs and B-cells (0.25 x 10^6^ to 0.5 x 10^6^) were labeled for 30 minutes on ice with 20 µg ml^-1^ 14.4.4 scF_V_-AF647-biotin, then washed with 1x HBSS (Life technologies) supplemented with 1% FCS (Biowest), 2 mM CaCl_2_ and 2 mM MgCl_2_ (Merck) and subsequently allowed to adhere on ICAM-1 coated glass slides before TOCCSL imaging. To reduce the likelihood of oligomer detection and create a monomer control, we pre-mixed the 14.4.4 scF_V_-AF647-biotin with unlabeled 14.4.4 scF_V_-biotin in a 1:10 to 1:15 ratio before labeling the cells. To serve as fluorescence dimer control, we pre-mixed the 14.4.4 scF_V_-AF647-biotin with mSav-AF647 at a 1:5 ratio for 30 minutes on ice prior to staining the cells with the conjugate. To induce dimerization directly on cells, we first stained BMDCs with 20 µg ml^-1^ 14.4.4 scF_V_-AF647-biotin, washed, and treated the samples with 125 nM divalent streptavidin (*trans* configuration). TOCCSL experiments were performed on setup #2 in TIRF mode and at room temperature as described (Brameshuber et al., 2018; Brameshuber et al., 2010; Moertelmaier et al., 2005; Ruprecht et al., 2010). I-E^k^-bound 14.4.4 scF_V_-AF647 was excited with the 640 nm laser (LCX-640-500) or the 637 nm laser (OBIS-637-140) at a power density of 0.1-0.2 kW cm^-2^ and recorded with an illumination time of 20 ms. The emission channel was filtered with the ET700/75 bandpass filter (Chroma). The unmasked cell region of interest was adjusted with a slit aperture (width 5-10 µm) and photobleached for a total time of 750-1000 ms with a power density of 2-3 kW cm^2^ in epifluorescence and TIRF mode. To confirm that photobleaching was complete, we included in some experiments a control image directly following the bleach pulse (100-750 ms delay). In order to observe well-separated fluorescence emitters diffusing back into the photobleached region, the recovery time was adjusted to 2-10 s for cells that had been stained under saturating conditions and 2-20 s for monomer controls. The first image after passing the recovery time was used to measure brightness values of the recovered as of then unbleached molecules.

Data recorded from TOCCSL experiments were analyzed as described in (Brameshuber et al., 2010). We applied a custom-made Matlab-based maximum likelihood estimator to extract XY coordinates, integrated brightness *B* values, the full width at half-maximum (FWHM) and the local background of individual fluorescence signals in the TOCCSL images (Moertelmaier et al., 2005). The brightness values *B* of single AF647-scFv molecules pooled from the TOCCSL images of the monomer control (1:15 premix of labeled to unlabeled 14.4.4 scF_V_) were used to calculate the probability density function (pdf) of monomers, *ρ*_1_(*B*). Because of the independent photon emission process of single fluorescent emitters, the corresponding pdfs of *N* colocalized emitters can be calculated by a series of convolution integrals according to **equation (5)**.

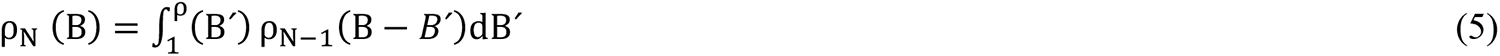

A weighted linear combination (α_N_) of these pdfs was used to calculate the brightness distribution of a mixed population of monomers and higher-order multimers.

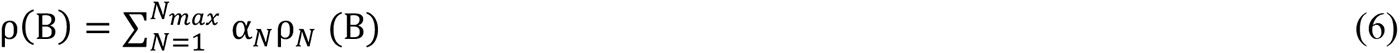

A minimum of 180 brightness values was applied to calculate *ρ*_1_(B) and *ρ*_N_ (B) from all TOCCSL images of multiple cells of one experimental condition. A least-square fit with **equation (6)** was employed to determine the weights of the individual pdfs, *α_N_*, with 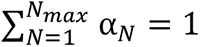

### Ensemble FRET imaging

T-cells (0.25 x 10^6^) were stained with saturating amounts (12 µg ml^-1^) of H57 scF_V_-AF555 (FRET donor) for 30 minutes on ice and washed into imaging buffer (10 ml of 1x HBSS (Life technologies) supplemented with 0.4 mg ml^-1^ ovalbumin (Merck), 2 mM CaCl_2_ and 2 mM MgCl_2_ (Merck)) and kept on ice until imaging proceeded. Stained T-cells were allowed to settle for 2 minutes on functionalized PLBs, which were equipped with unlabeled ICAM-1, B7-1 (100 molecules per µm^2^) and I-E^k^/peptide-AF647 (FRET acceptor) at different densities. Synaptic FRET was recorded as a consequence of TCR-pMHCII binding for up to 10 minutes (i.e. 2-12 minutes after cells had made first bilayer contact).

To examine the oligomeric state of pMHCIIs, 0.25 x 10^6^ to 0.5 x 10^6^ BMDCs and B-cells were labeled on ice for 30 minutes with the FRET donor 14.4.4 scF_V_-AF647 and the FRET acceptor 14.4.4 scF_V_-AF555 (10 µg ml^-1^ each) using pre-mixed (1:1, 1:2, 1:3 or 1:4) AF647 and AF555 probe cocktails. Cells were then washed with imaging buffer (1x HBSS (Life technologies) supplemented with 1% FCS (Biowest), 2 mM CaCl_2_ and 2 mM MgCl_2_ (Merck) and subsequently allowed to adhere on ICAM-1 coated glass slides prior to imaging in TIRF mode at 22.5°C. To dimerize I-E^k^ within the cell membrane, 0.5 x 10^6^ BMDCs were stained for 30 minutes on ice with 14.4.4 scF_V_-AF647-biotin and 14.4.4 scF_V_-AF555-biotin premixed at a 1:1 ratio and subsequently washed with imaging buffer (1x HBSS (Life technologies) supplemented with 1% FCS (Biowest), 2 mM CaCl_2_ and 2 mM MgCl_2_ (Merck)). Biotinylated scF_V_ were then crosslinked with varying concentrations of divalent streptavidin (ranging from 1 to 1000 nM) for 30 minutes on ice to induce I-E^k^-dimer formation, before allowing the cells to adhere to ICAM-1-coated glass slides.

For FRET measurements of bilayer-resident pMHCIIs, we functionalized PLBs with unlabeled I-E^k^/MCC which were then labeled using increasing amounts of 14.4.4 scF_V_-AF555 and 14.4.4 scF_V_-AF647 which had been premixed at a 1:1, 1:2 or 1:3 ratio. PLBs were washed with imaging buffer (1x HBSS (Life technologies) supplemented with 1% FCS (Biowest), 2 mM CaCl_2_ and 2 mM MgCl_2_ (Merck)) prior to imaging.

For all FRET experiments, the emission beam path was separated into two spectral channels employing an Optosplit 2 device (Cairn Research) which was equipped with a beam splitter (ZT640rdc, Chroma) and adequate emission bandpass filters (ET575/50 or ET640SP, ET655LP, Chroma). The following alternating excitation protocol was used to record FRET donor and FRET acceptor channels: a low-intensity AF647 acquisition (to monitor the AF647 signal, OBIS-637-140, power < 0.01 kW cm^-2^) was succeeded by a low-intensity AF555/FRET acquisition (to record the AF555 and FRET signal before FRET acceptor photo bleaching, OBIS-532-100-LS, power ∼ 0.01 kW cm^-2^), followed by a high-intensity 640-nm bleach pulse (to ablate AF647, power >2 kW cm^-2^), a second low-intensity AF647 acquisition (to verify AF647 ablation, power ∼ 0.01 kW cm^-2^) and finally a second AF555/FRET acquisition (to record the AF555 signal after FRET acceptor bleaching and verify FRET ablation, power ∼ 0.01 kW cm^-2^). The time lapsing between the two AF555/FRET acquisitions was less than 500 milliseconds. Quantitative imaging was ensured by minimizing photobleaching at a given illumination time of 30 ms. To this end, the power levels of the 532 nm FRET donor excitation laser were adjusted to keep photobleaching between consecutive images below 1%. I-E^k^/peptide-AF488 was excited with the 488 nm laser (OBIS-488-150-LS) at a power density of 0.1-0.2 kW cm^-2^ and the emission was cleaned up with the ET525/50 (Chroma) filter.

For FRET sensitized emission analysis, donor and acceptor channels were superimposed using an affine transformation matrix (including shift, stretch, rotation, and tilt), and intensity registration, both generated from a fluorescent bead reference (TetraSpeck Fluorescent Microspheres Size Kit, Thermo Fisher Scientific). Net fluorescence intensities were obtained by pixel-wise subtraction of the camera background from the raw intensity images. To compensate for laser illumination profiles, we performed a pixel-wise flatfield correction by means of median averaging ten separately recorded PLB samples representing a similar fluorophore density as the sample employed for FRET measurements.

FRET donor recovery after acceptor photobleaching analysis was performed as described (Huppa et al., 2010). After background subtraction, the AF555 intensity (FRET donor) before FRET acceptor photobleaching (*DDpre*) was subtracted from the AF555 intensity after FRET acceptor photobleaching (*DDpost*). The difference was then divided by *DDpost* to obtain the FRET yield according to **equation (7).**

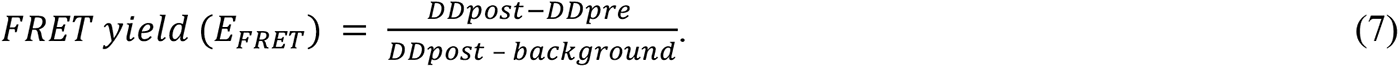

Apparent FRET efficiencies were calculated based on **equation (8)** with *α* = cross-excitation, *β* = bleedthrough, *DA* = donor excitation, acceptor channel (FRET channel), *AA* = acceptor excitation, acceptor channel (AF647 signal).

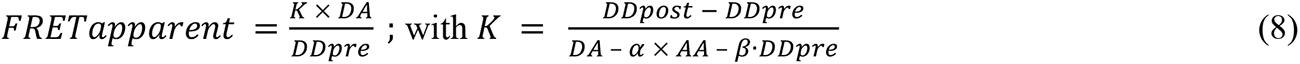

The TCR occupancy, i.e. the ratio of ligand-engaged and total TCRs was calculated based as shown in **equation (9)**:

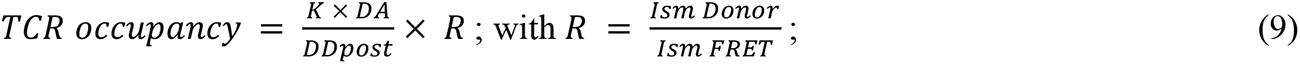

with *I_sm_ Donor* = single molecule FRET donor intensity

and *I_sm_ FRET* = single molecule FRET intensity,

as determined experimentally by the average intensity of single-molecule FRET events and the average intensity of single-molecule FRET donor signals (see single molecule FRET imaging and **Supp. Fig. 14E**). Additionally, we estimated *R* based on the transmitted spectrum of light of the filters sets ET575/50 (62.1%) and ET655LP (74.3%) as follows:

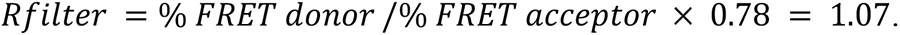

Since the TCR occupancy is directly proportional to the FRET yield (Axmann et al., 2015), we established a relationship between these two parameters by plotting the synaptic FRET yields obtained from FRET donor recovery after acceptor photobleaching measurements against TCR occupancies obtained from the same T-cell synapses. By fitting a linear regression, we experimentally determined the conversion factor *C* = 1.21 (for *R_exp_* = 1.0) to calculate TCR occupancies directly from FRET yields according to **equation (10)**.

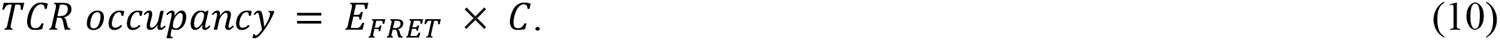

The experimentally determined conversion factor *C* = 1.21 was comparable with the estimated values of *C* = 1.29 (for *R_filter_* = 1.07) and *C* = 1.28 (obtained from the maximal FRET efficiency of the FRET system *EmaxFRET* = 1/0.78 = 1.28).

### Single molecule FRET imaging

Single molecule FRET recordings were conducted with the same experimental setup, hardware settings and filters as used for recording ensemble FRET. To attenuate bleed through from the FRET donor fluorophore into the FRET acceptor channel, we opted to stain approximately 10% of the TCRs on T-cells with the H57 scF_V_-AF555 by premixing 1 part of H57 scF_V_-AF555 with 9 parts of unlabeled H57 scF_V_ prior to cell staining (total 10 µg ml^-1^). We also adjusted I-E^k^/MCC-AF647 densities on PLBs to 5-30 I-E^k^ µm^-2^, to reduce cross-excitation of the FRET acceptor fluorophore.

To visualize earliest single TCR-pMHCII binding events, we employed in addition the caged MCC peptide derivative ANERADLIAYLK[NVOC]QATKGGSC-AF647 loaded onto I-E^k^ (I-E^k^/MCC[NVOC]-AF647) which represents a null ligand unless it becomes uncaged by means of a short 405 nm laser pulse (0.17 kW cm^-2^, 100 ms). One to two seconds after the 405 nm pulse, individual TCR-pMHCII binding events appeared as single molecule FRET events, disappeared in single steps and aligned with single FRET acceptor fluorophores (I-E^k^/MCC-AF647). We recorded smFRET events with an illumination time of 20 ms and applied a 532 nm power density of 0.1 kW cm^-2^ (OBIS-532-100-LS or LCX-532-500) to excite the donor fluorophore AF555 and a 640 nm power density of 0.05 kW cm^-2^ (OBIS-637-140 or LCX-640-500) to excite the acceptor fluorophore AF647. I-E^k^/K99A-AF488 was excited with the 488 nm laser (LBX-488-200) at a power density of 0.1-0.2 kW cm^-2^ and the emission was filtered with an ET525/50 (Chroma) filter.

Intensity and positional accuracy of single molecule donor and single molecule FRET events as a consequence of TCR-pMHCII binding were calculated with the ImageJ plugin ThunderSTORM (Ovesny et al., 2014) and filtered for sigma (< 320 nm) and positional accuracy (< 150 nm).

To measure the synaptic off-rate of 5c.c7 TCR:I-E^k^/MCC interactions at 26 °C in the presence or absence of I-E^k^/K99A, we adjusted the I-E^k^/MCC-AF647 density on PLBs to 11 I-E^k^/MCC-AF647 µm^-2^ and ICAM-1 and B7-1 to 100 µm^-2^. I-E^k^/K99A-AF488 was either completely absent or present at a density of 1060 I-E^k^/K99A µm^-2^. For off-rate determination, we recorded the appearance of smFRET traces at different time lags (56, 84, 252, 476, 924, 1820 ms) to correct for photobleaching. We then listed the smFRET traces according to their trajectory length and created a normalized inverse cumulative decay function for each time lag as described in (Axmann et al., 2015). The resulting decay function was fitted with a single exponential decay function (start 1.0, plateau 0) to calculate (app t_off_ = 1/app k_off_), the apparent off-rate (**Supp. Fig. 16E**). Lastly, we plotted the calculated apparent off-rates against the respective time lags and fitted the off-rate (t_off_) and the bleach constant c_bleach_ based on **equation (11)**.

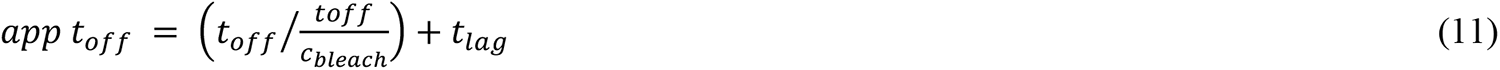

### Calcium imaging

Intracellular calcium levels were measured with the ratio-metric dye Fura-2-AM (Thermo Fischer Scientific) as published (Roe et al., 1990). 1-2 x 10^6^ antigen-experienced 5c.c7 TCR-transgenic T-cells were incubated with 2 µM Fura-2-AM (Life technologies) in T-cell growth medium for 15-20 minutes at room temperature and subsequently washed with warm (room temperature) imaging buffer (14 ml of 1x HBSS (Life technologies) supplemented with 1% FCS (Biowest), 2 mM CaCl_2_ and 2 mM MgCl_2_ (Merck)). Immediately afterwards, T-cells were seeded onto functionalized PLBs featuring increasing densities of I-E^k^ in complex with agonist or non-agonist (bystander) peptides and unlabeled B7-1 and ICAM-1 (100 µm^-2^). The usage of 16-well Lab-Tek Chambers (Nunc) enabled us to record up to 12 different conditions (i.e. various or different concentrations of I-E^k^) in a single experimental run. Calcium experiments were carried out on setup #1 or setup #3 at 37 °C. On setup #1, Fura-2-AM was excited using a monochromatic light source (Leica) coupled into an inverted Leica DMI4000B microscope that was equipped with a 20x objective (HCX PL Fluotar 20x, NA = 0.5, Leica), the dichroic beamsplitter FU2 (Leica) and the bandpass filter ET525/36 (Leica). Calcium signalling was measured ratiometrically by continuous switching between 340 nm and 387 nm excitation, which was achieved with the use of a fast excitation filter wheel (Leica) featuring the excitation bandpass filters 340/26 and 387/11 (Leica). The total recording time was 10-20 minutes with an image recorded every 15-20 seconds. An automated XY stage (Leica) allowed fast changes between different positions (i.e. Lab-Tek Chamber wells). Precise temperature control was carried out with a heating unit and a box enclosing the microscope (Leica).

For calcium imaging of T-cells approaching DNA origami structures, we used setup #3 at the Technical University of Vienna. Fura-2 was excited with a monochromatic light source (Polychrome V, TILL Photonics) coupled to a Zeiss Axiovert 200M which was equipped with a 10x objective (UPlanFL N 10x, NA = 0.3, Olympus), a 1.6x tube lens, a long-pass filter (T400lp, Chroma), the emission filter ET510/80 (Chroma) and the EMCCD camera Andor iXon Ultra 897. Imaging was performed with excitation at 340 and 380 nm with illumination times of 50 and 10 ms, respectively. The total recording time was 10 min with one image per second. Experiments were carried out at 37 °C using an in-house–built incubator equipped with a heating unit for precise temperature control.

Calcium image analysis was carried out either with a custom-code written in Matlab and as described in (Hellmeier et al., 2021b). In detail, we first generated the ratio image of the 340 nm / 380 nm image for every frame. Individual T-cells were then tracked using an in-house Matlab algorithm based on (Gao and Kilfoil, 2009) and the positional information of each cell was used to extract the intensity value from the ratio image stack. Fura-2 intensity traces were either normalized to the starting value at timepoint zero or to the median of all Fura-2 intensity traces of a negative control (ICAM-1 100 µm^−2^, B7-1 100 µm^−2^). Traces were categorized into “activating”, oscillatory and “non-activating” based on an activation threshold defined by a receiver operating curve between a negative control and a positive control (I-E^k^/MCC > 10 µm^−2^, ICAM-1 100 µm^−2^, B7-1 100 µm^−2^). T-cells were categorized as “activating” when the Fura-2 intensity value remained 80% of the entire trace above the threshold, “non-activating when it remained 80% of the trace below the threshold, otherwise the cell was classified as oscillatory. The generation of dose response curves and the bootstrap analysis of EC50 data is best described in (Hellmeier et al., 2021b). The calcium imaging data was fitted to a three (hillslope n = 1.0) or four-parameter dose response curve with **equation (12)** to extract the activation threshold (TA) at half-maximum response.

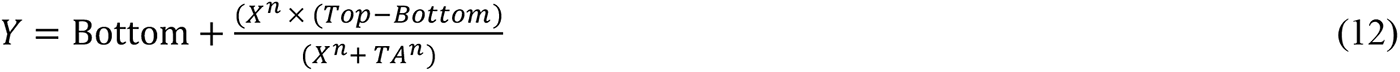

### STED microscopy

BMDCs and B-cells (0.25 x 10^6^ to 0.5 x 10^6^) were stained with 20 µg ml^-1^ 14.4.4 scF_V_-abSTAR635P for 30 minutes on ice and washed with imaging buffer (1x HBSS (Life technologies) supplemented with 1% FCS (Biowest), 2 mM CaCl_2_ and 2 mM MgCl_2_ (Merck). Cells were allowed to adhere on ICAM-1 coated glass slides for 10 minutes at 25°C before fixation. I-E^k^/MCC molecules were absorbed to a glass-slide and stained with limited amounts of 14.4.4 scF_V_ abSTAR635P to visualize single fluorescent emitters. Samples were fixed for 15 minutes at room temperature applying 4% PFA (Thermo Fisher Scientific), washed with 1x PBS, mounted in Abberior Mount Solid (Abberior) and stored at 4°C until imaging. STED imaging was performed on a Zeiss Axio Imager A2 upright microscope equipped with an 100x objective (alpha Plan-Apochromat 100x, NA = 1.46, Oil DIC M27, Zeiss) and the Stedycon super-resolution microscopy unit (Abberior Instruments). Scanning was performed with the built-in QUAD beam scanner with a pixel dwell time of 250 microseconds at a pixel size of 25 nm. A pulsed 640 nm diode laser with an output of 1.2 mW at 40 MHz (max. 5 mW), and > 20 pJ/pulse (< 150 ps) was used to excite abSTAR635P molecules and a pulsed 775 nm STED depletion laser with an output of 1.25 W, and > 30 nJ/pulse (jitter < 100 ps) was used for stimulated emission depletion. Time gating delay was set to 1 nanosecond and gate width to 6 nanoseconds. The Stedycon was controlled via a Chrome browser-based software provided by the manufacturer (Abberior Instruments).

### Simulations and software used for analysis

Microscopy images were processed and analyzed with the open-source image processing package Fiji (Schindelin et al., 2012). More extensive image analysis was carried out with custom-written algorithms written in Matlab (MathWorks). XY localization, intensity and positional accuracy of single molecules were calculated with the Fiji plugin ThunderSTORM (Ovesny et al., 2014). Prism 9 was used for statistical data analysis and evaluation (GraphPad).

### STED autocorrelation analysis

To judge the randomness of pMHCII in STED microscopy data, we calculated ACFs for four different 2.5 x 2.5 µm regions of interest per cell and compared these experimental samples with ACFs of simulated STED images of randomly distributed molecules obtained with experimentally determined parameters (Sengupta et al 2011). To characterize the point-spread function of the imaging system, we sparsely labeled activated BMDCs and B-cells with the 14.4.4 scF_V_-abSTAR65P and fitted the recorded single molecules signals using the ImageJ plugin ThunderSTORM. The background was determined from the mean fluorescence signal of non-fluorescent areas within each region of interest. I-E^k^ densities were calculated by dividing the background subtracted mean intensity in fully labeled cells by the average single-molecule intensity, which yielded 15-108 I-E^k^ molecules per µm^2^ on activated BMDCs and 24-44 I-E^k^ molecules per µm^2^ on activated B-cells. Simulated images of random signal distributions were then generated using the parameters intensity (I), and sigma (σ) derived from single-molecule signals and background and molecular density from the cell regions of interest using an in-house written Matlab code (for details see (Rossboth et al., 2018)).

### Simulation of a random distributions for blinking analysis

A 7 x 7 µm^2^ region of interest featuring randomly distributed molecules at a specified density was simulated as described (Platzer et al., 2020; Rossboth et al., 2018). To arrive at simulations to be used for comparison with cell-associated PALM data, the approximate expression levels of I-E^k^ (14.4.4 scF_V_-AF647-biotin + mSAv-cc-PSCFP2), DEC205 (NLDC145 mAb-AF647-biotin + mSAv-cc-PSCFP2) and CD18 (M18/2 mAb-AF647-biotin + mSav-cc-PS-CFP2) within a region of interest were determined by dividing the number of PS-CFP2 localizations with the mean number of detections N per fluorescence molecule as obtained from the blinking analysis (**Supp. Fig. 7A**). The experimentally-derived probability distribution of the number of detections *N* was randomly assigned to every simulated localization to include the blinking of PS-CFP2. Finally, all positions were shifted into a random direction by a distance drawn from a normal distribution with an average value of zero and localization precision (51 ± 12 nm) as standard deviation to simulate a PFA-fixed scenario. In order to simulate live cell scenarios, we additionally shifted the localizations based on the diffusion parameters F (fast diffusing fraction), D1 (diffusion constant of fast-moving fraction), D2 (diffusion constant of slow-moving fraction) of I-E^k^ on BMDCs (F = 0.83, D1 = 0.284 µm^2^ s^-1^, D2 = 0.046 µm^2^ s^-1^, **Supp. Fig. 7B**), I-E^k^ on B-cells (F = 0.77, D1 = 0.150 µm^2^ s^-1^, D2 = 0.024 µm^2^ s^-1^, **Supp. Fig. 7B**), DEC205 (F = 0.9, D1 = 0.065 µm^2^ s^-1^, D2= 0.0006 µm^2^ s^-1^, (Manzo et al., 2012)), and CD18 (F = 0.3, D1 = 0.03 µm^2^ s^-1^, D2= 0.0 µm^2^ s^-1^ (Mukherjee et al., 1998) and a time-lag of 2.3 ms.

### Monte Carlo simulation of cluster detection using PALM

Ripley’s K functions were determined from Monte Carlo simulations of nanoclusters with a cluster radius of 20, 40, 60, or 100 nm, a fraction of molecules residing inside clusters between 20 and 100%, 3, 5, 10, 15, and 20 clusters per µm², an average molecular density of 100 molecules per µm² and the experimentally determined blinking statistics of PS-CFP2 determined under 4% PFA-fixed and non-fixed imaging conditions (**Supp. Fig. 7A**). Functions were compared to those calculated from random localization distributions and categorized in “Resolvable”, “Borderline” or “Not resolvable” as described in (Platzer et al., 2020).

### Monte Carlo simulation of TOCCSL

TOCCSL results were simulated based on experimentally derived parameters using a Monte Carlo based approach as outlined in detail in Bodner et al., submitted (2023, https://owncloud.tuwien.ac.at/index.php/s/BqEeg4kAJU3X8ZY). In Brief, a cell size of 10.5 x 10.5 µm² containing 500 molecules per µm² diffusing with an average diffusion constant of 0.244 µm² s^-1^ (D = 0.83*0.284 + 0.17*0.046 µm² s^-1^, see **Supp.** **Fig. 7B**) was simulated. An area of 7x7 µm² was photobleached for 750 ms with bleaching parameters derived from photobleaching curves of AF647 illuminated with a laser power density of 3 kW cm^-^² and a diffraction-affected laser intensity profile as determined experimentally. For TOCCSL simulations, the fraction of molecular dimers was varied between 3 and 100% while keeping the overall density of molecules constant. For each dimer fraction the fraction of detected dimers after a recovery time of 2 s was determined from the simulations. Simulations were repeated 1000 for each dimer fraction.

### Monte Carlo simulation of FRET efficiency

FRET efficiencies of randomly distributed donor and acceptor molecules at increasing densities were simulated as described (Brameshuber et al., 2018). We used algorithms custom-written in Matlab to simulate random X and Y positions of pMHCII molecules for increasing surface densities between 20 and 1000 molecules per µm^2^ (increment 20). Due to higher simulated surface densities compared to our previous study (Brameshuber et al., 2018), we considered not only the nearest neighbor of every donor molecule for calculation the FRET efficiency, but also the second nearest neighbor. To this end, the distances *d_1,2_* between every donor molecule and the two nearest acceptor molecules were calculated and used to determine the FRET efficiencies *E_1,2_*, according to **equation (13)**, with *R_0_* = 5.1 nm, the Förster radius for the AF555/AF647 dye pair.

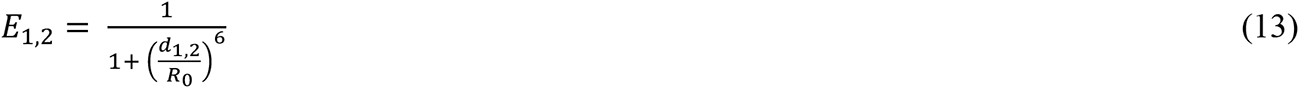

The overall FRET efficiency for a donor with two acceptor molecules (Koushik et al., 2009) was calculated according to **equation (14)**.

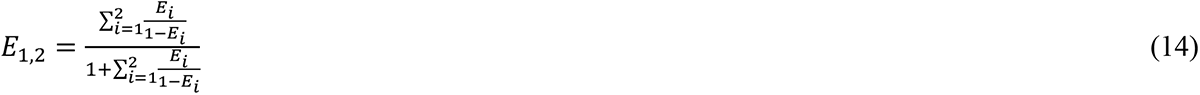

For every surface density, the simulation was repeated 20-times to arrive at a mean and range of FRET efficiencies. FRET efficiencies were plotted against the total density (sum of donor and acceptor molecules).

## Supporting information

https://drive.google.com/drive/folders/1ht8mdS0XgqRXQLcAqc8YTwD_q-iFcNqd

## Data availability

Source data are provided with this manuscript. The datasets generated and/or analyzed are available from the corresponding authors upon reasonable request.

## Code availability

Links to the software packages for analysis can be obtained from the cited papers.

## AUTHOR CONTRIBUTIONS

RP, MB, and JBH conceived the project. RP, MB, and JBH wrote the manuscript. RP designed and produced molecular probes, performed most experiments and analyzed data. IDP produced molecular probes and developed software tools. JH and ES designed and performed origami-based calcium experiments. PS performed lysis gradient centrifugation analysis. CB, JG and MB developed software tools and performed data simulations. MFT produced and purified peptides. ES, GJS and HS contributed important ideas.

## ACKNOWLEDGEMENTS

This work was supported by the Austrian Science Fund (FWF) through the PhD program Cell Communication in Health and Disease W1205 (JBH and HS), the projects I4662-B (MB), I4740-B (CB), V538-B26 (ES, JH), and P25775-B2 (JBH) and the Vienna Science and Technology Fund (WWTF) projects LS13-030 (GJS, JBH) and LS14-031 (JBH). Funding was further provided by a predoctoral fellowship from the Boehringer Ingelheim Fonds (RP).

