## Supplementary material for "Monomeric agonist peptide/MHCII complexes activate T-cells in an autonomous fashion": https://drive.google.com/drive/folders/1ht8mdS0XgqRXQLcAqc8YTwD_q-iFcNqd

### Supplementary Table 1

## A

| **Layout** | **Ligand 1** (mSav-V) | **Ligand 2** (mSav-X) | **No ligand** (%) | **Single occupancy - ligand 1 (%)** | **Single occupancy - ligand 2 (%)** | **Double occupancy (%)** |
|---|---|---|---|---|---|---|
| **M** | I-E$^k$/MCC | --- | 39.51 ± 1.38 | 60.49 ± 1.38 | --- | --- |
| **D10** | I-E$^k$/MCC | --- | 10.12 ± 1.23 | 43.39 ± 5.28 | --- | 46.50 ± 2.65 |
| **D10** | I-E$^k$/K99A | --- | 10.12 ± 1.23 | 43.39 ± 5.28 | --- | 46.50 ± 2.65 |
| **D10** | I-E$^k$/MCC | I-E$^k$/K99A | 9.87 ± 1.24 | 20.81 ± 1.85 | 22.31 ± 2.12 | 47.01 ± 2.73 |
| **D20** | I-E$^k$/MCC | I-E$^k$/K99A | 9.32 ± 0.73 | 24.35 ± 1.68 | 18.35 ± 0.87 | 47.98 ± 1.82 |

## B

| **Layout** | **Ligand 1** | **Ligand 2** | **n** | **$n_{low}$** | **$n_{high}$** | **$A_{max}$ (%)** | **$A_{max,low}$ (%)** | **$A_{max,high}$ (%)** | **$T_A$ ($\mu m^{-2}$)** | **$T_{A,low}$ ($\mu m^{-2}$)** | **$T_{A,high}$ ($\mu m^{-2}$)** | **# cells** | **# mice** |
|---|---|---|---|---|---|---|---|---|---|---|---|---|---|
| **mSAv** | I-E$^k$/MCC | --- | 1.85 | 0.98 | 2.72 | 93.58 | 79.40 | 107.75 | 0.33 | 0.25 | 0.45 | 166 ± 45 | 4 |
| **M** | I-E$^k$/MCC | --- | 4.00 | 0.30 | 7.70 | 93.15 | 85.05 | 101.25 | 0.40 | 0.33 | 0.49 | 186 ± 56 | 3 |
| **D10** | I-E$^k$/MCC | --- | 4.00 | 1.95 | 6.05 | 91.61 | 86.90 | 96.32 | 0.48 | 0.43 | 0.55 | 150 ± 35 | 3 |
| **D10** | I-E$^k$/MCC | I-E$^k$/K99A | 3.27 | 1.83 | 4.71 | 87.91 | 82.47 | 93.35 | 0.34 | 0.28 | 0.42 | 123 ± 41 | 4 |
| **D20** | I-E$^k$/K99A | I-E$^k$/K99A | 2.14 | 1.26 | 3.02 | 90.91 | 82.40 | 99.43 | 0.46 | 0.32 | 0.64 | 134 ± 23 | 4 |
| **D10** | I-E$^k$/K99A | --- | n.a. | n.a. | n.a. | n.a. | n.a. | n.a. | n.a. | n.a. | n.a. | 80 ± 25 | 3 |

## C

| **Layout** | **mSAv-** I-E$^k$/MCC | **M-** I-E$^k$/MCC | **D10-** I-E$^k$/MCC | **D10-** I-E$^k$/MCC + K99A | **D20-** I-E$^k$/MCC + K99A | **D10-** I-E$^k$/K99A |
|---|---|---|---|---|---|---|
| **mSAv-**I-E$^k$/MCC | - | n.s. | < 0.05 | n.s. | n.s. | < 0.001 |
| **M-**I-E$^k$/MCC | n.s. | - | n.s. | n.s. | n.s. | < 0.001 |
| **D10-**I-E$^k$/MCC | < 0.05 | n.s. | - | n.s. | n.s. | < 0.001 |
| **D10-**I-E$^k$/MCC + K99A | n.s. | n.s. | n.s. | - | n.s. | < 0.001 |
| **D20-**I-E$^k$/MCC + K99A | n.s. | n.s. | n.s. | n.s. | - | < 0.001 |
| **D10-**I-E$^k$/K99A | < 0.001 | < 0.001 | < 0.001 | < 0.001 | < 0.001 | - |

### Supplementary Figure 1

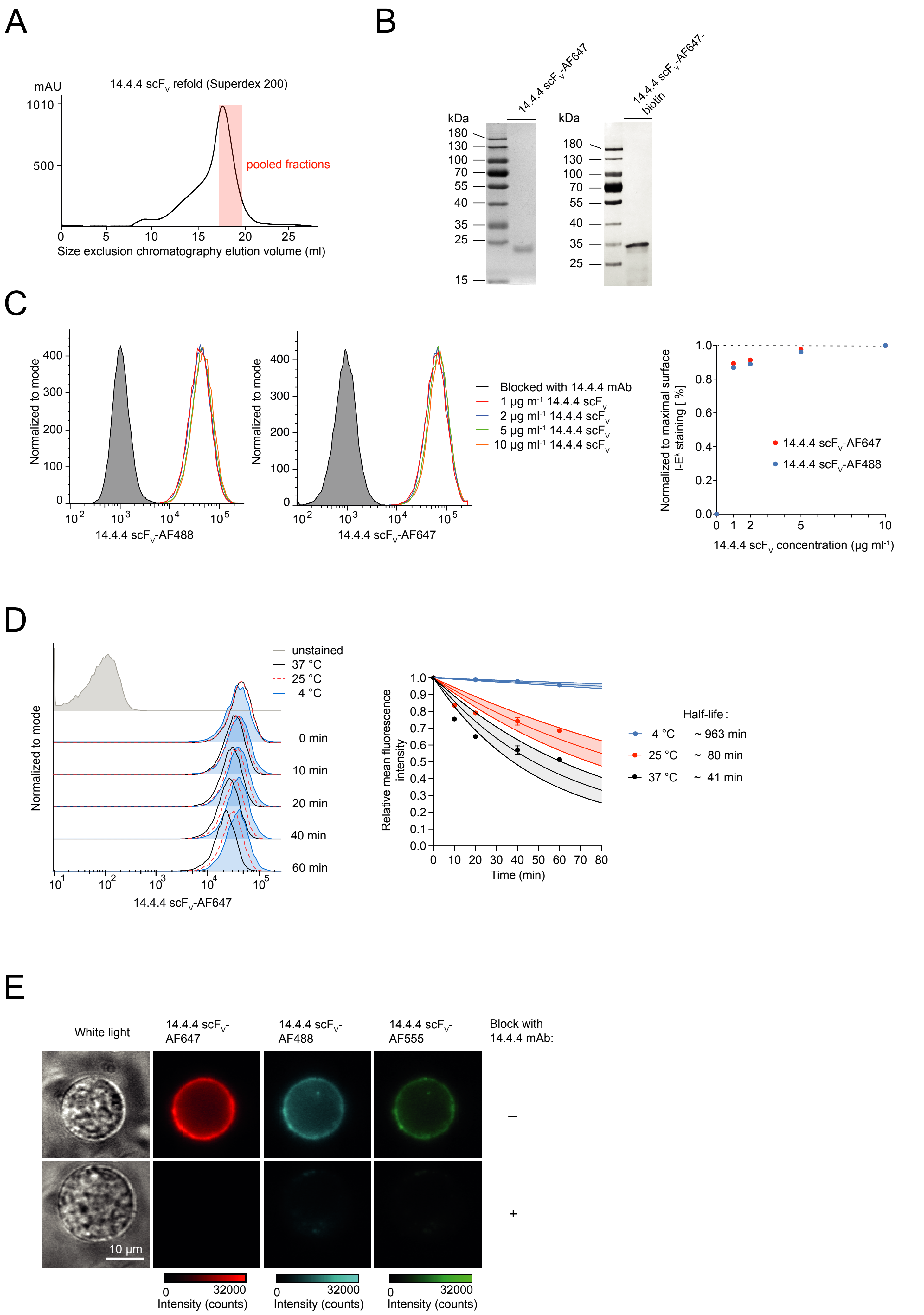

### Supplementary Figure 2

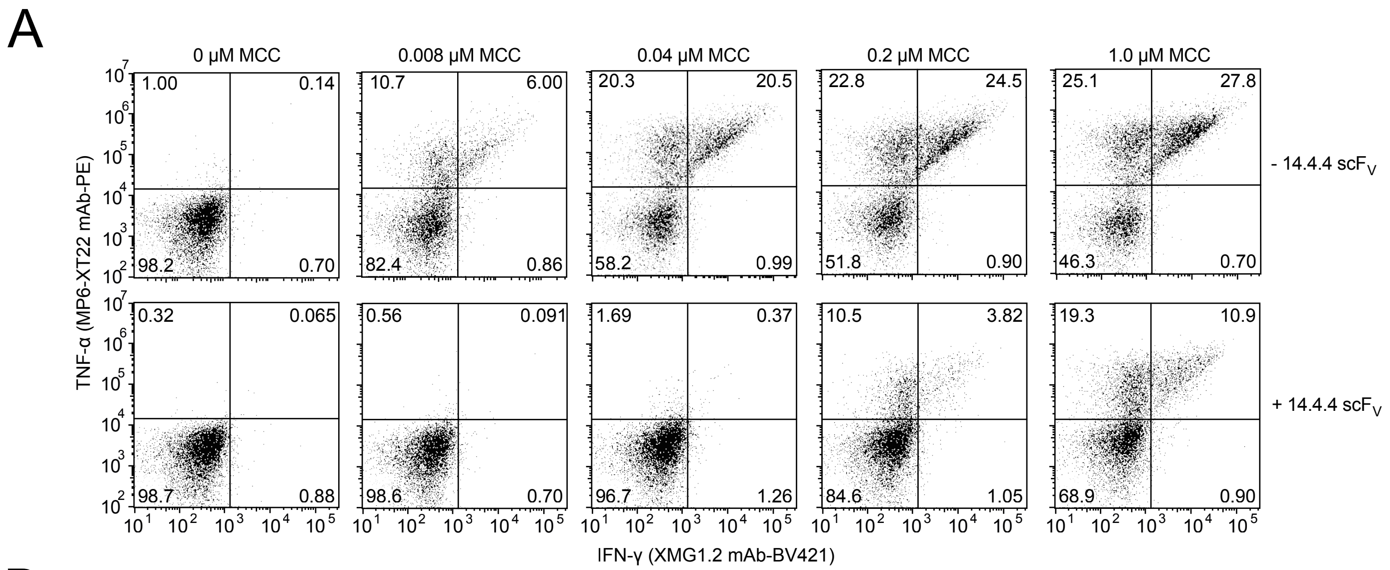


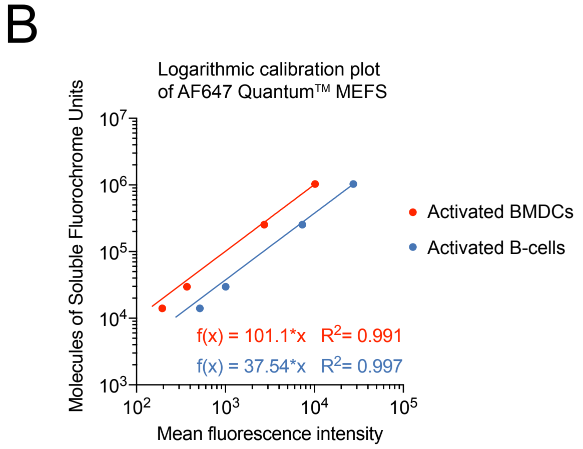

### Supplementary Figure 3

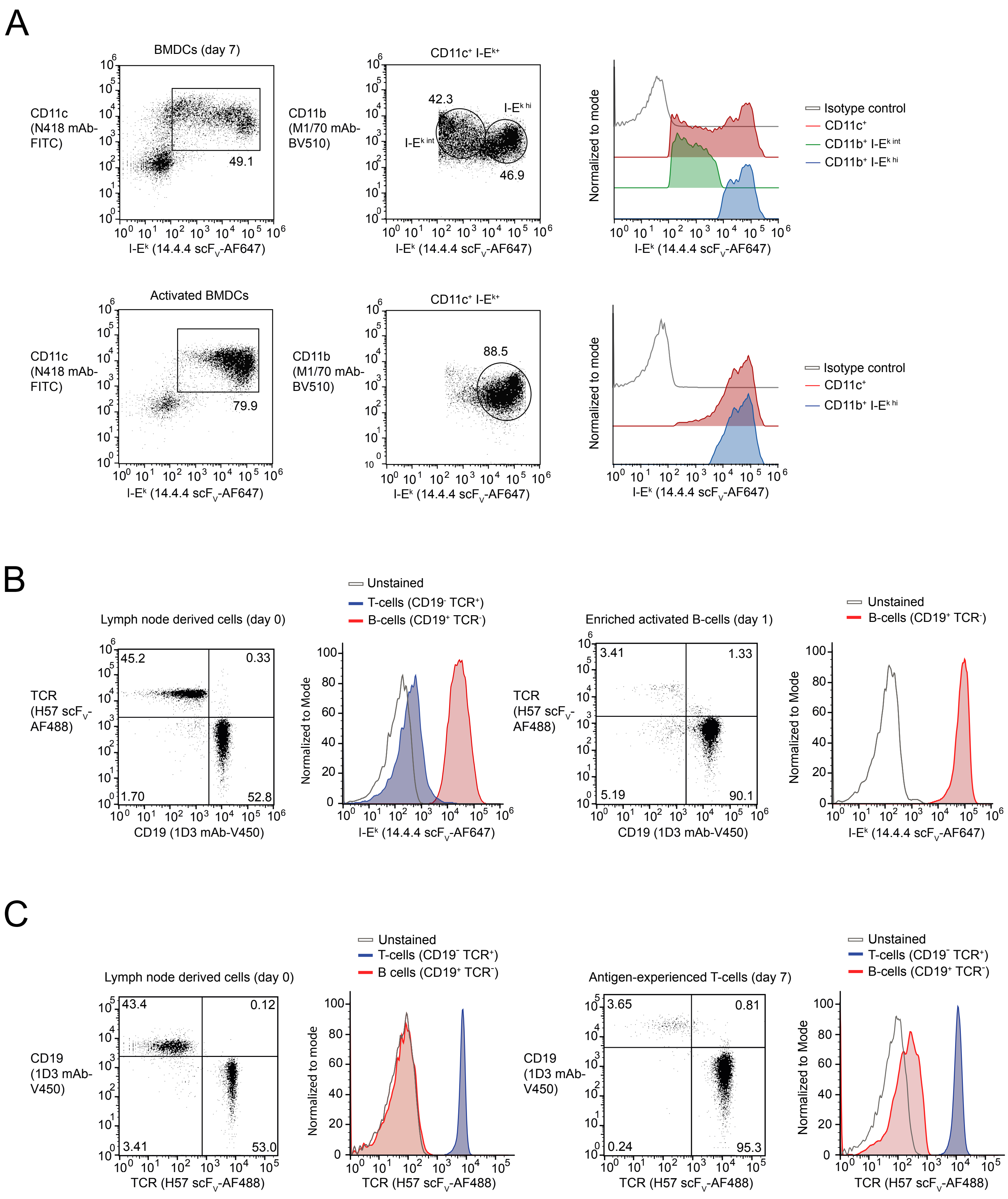

### Supplementary Figure 4

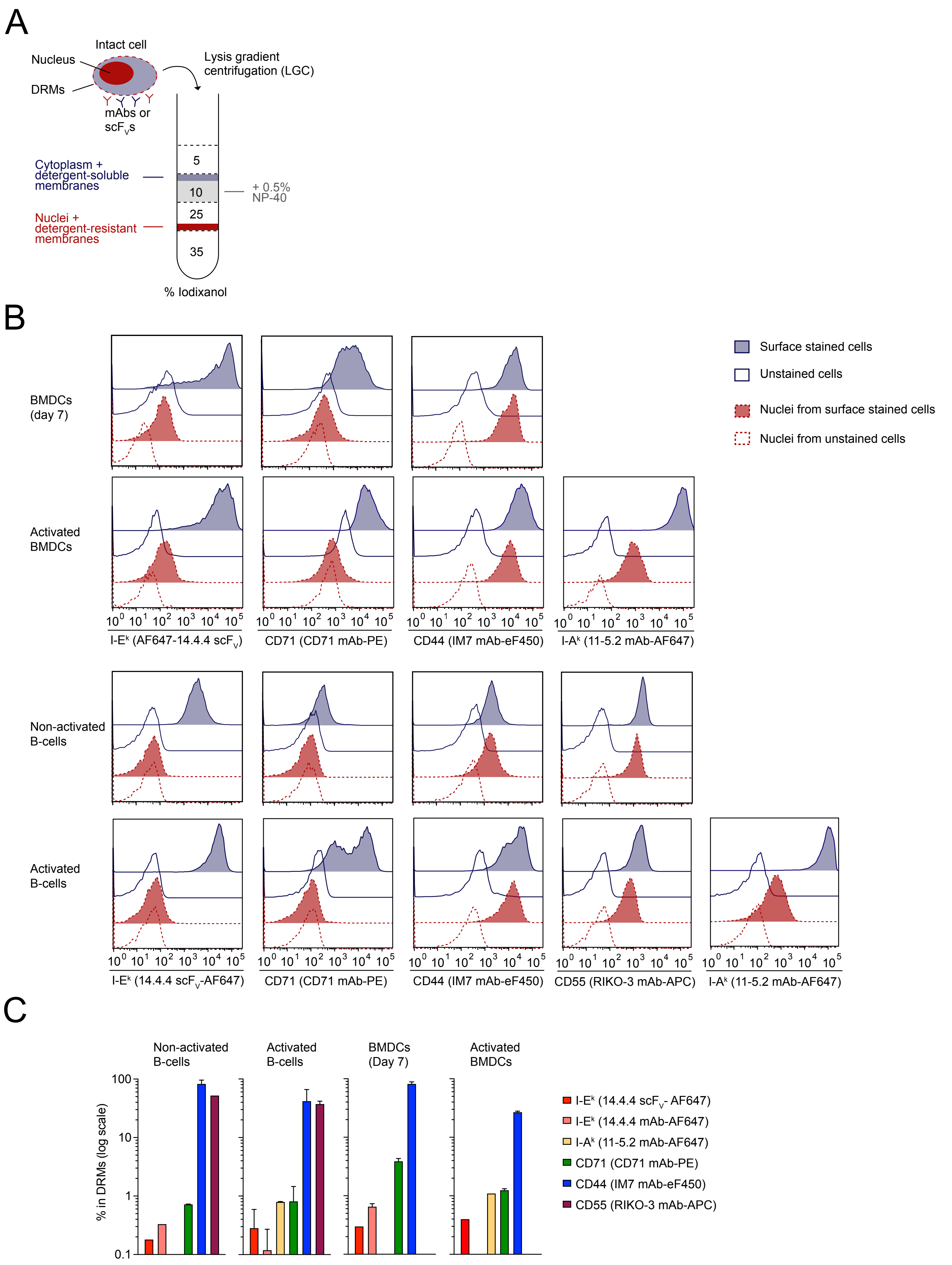

### Supplementary Figure 5

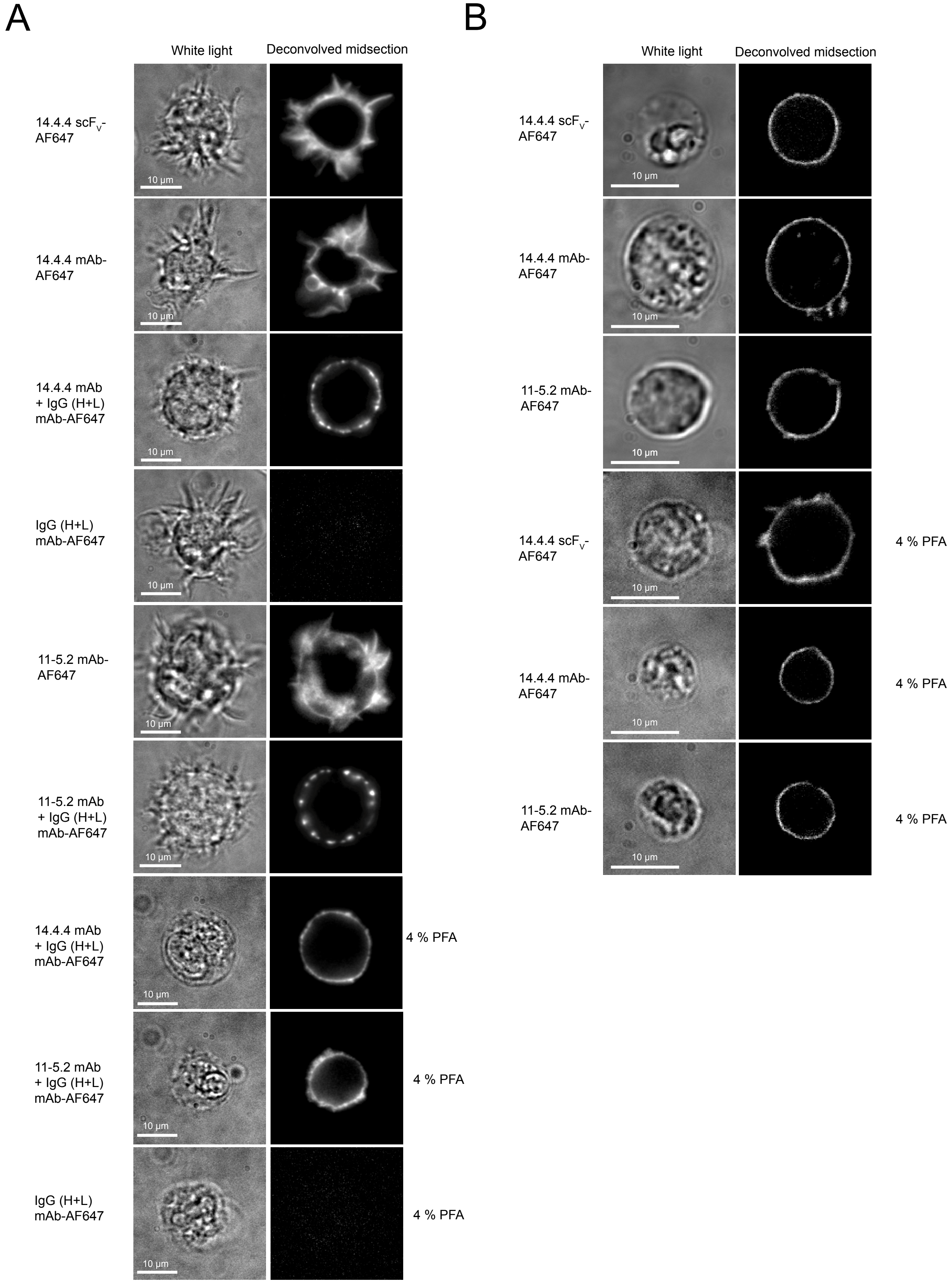

### Supplementary Figure 6

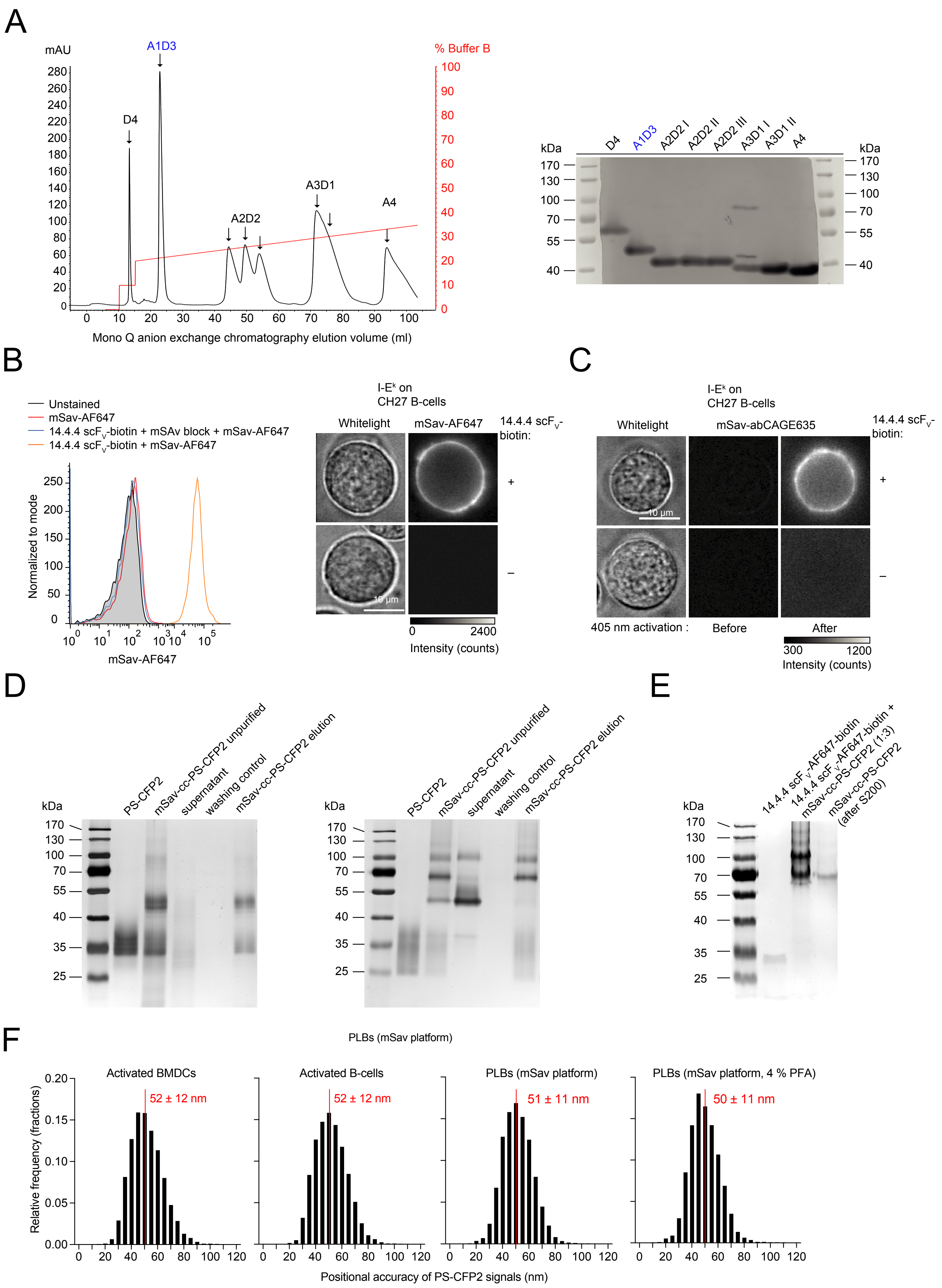

### Supplementary Figure 7

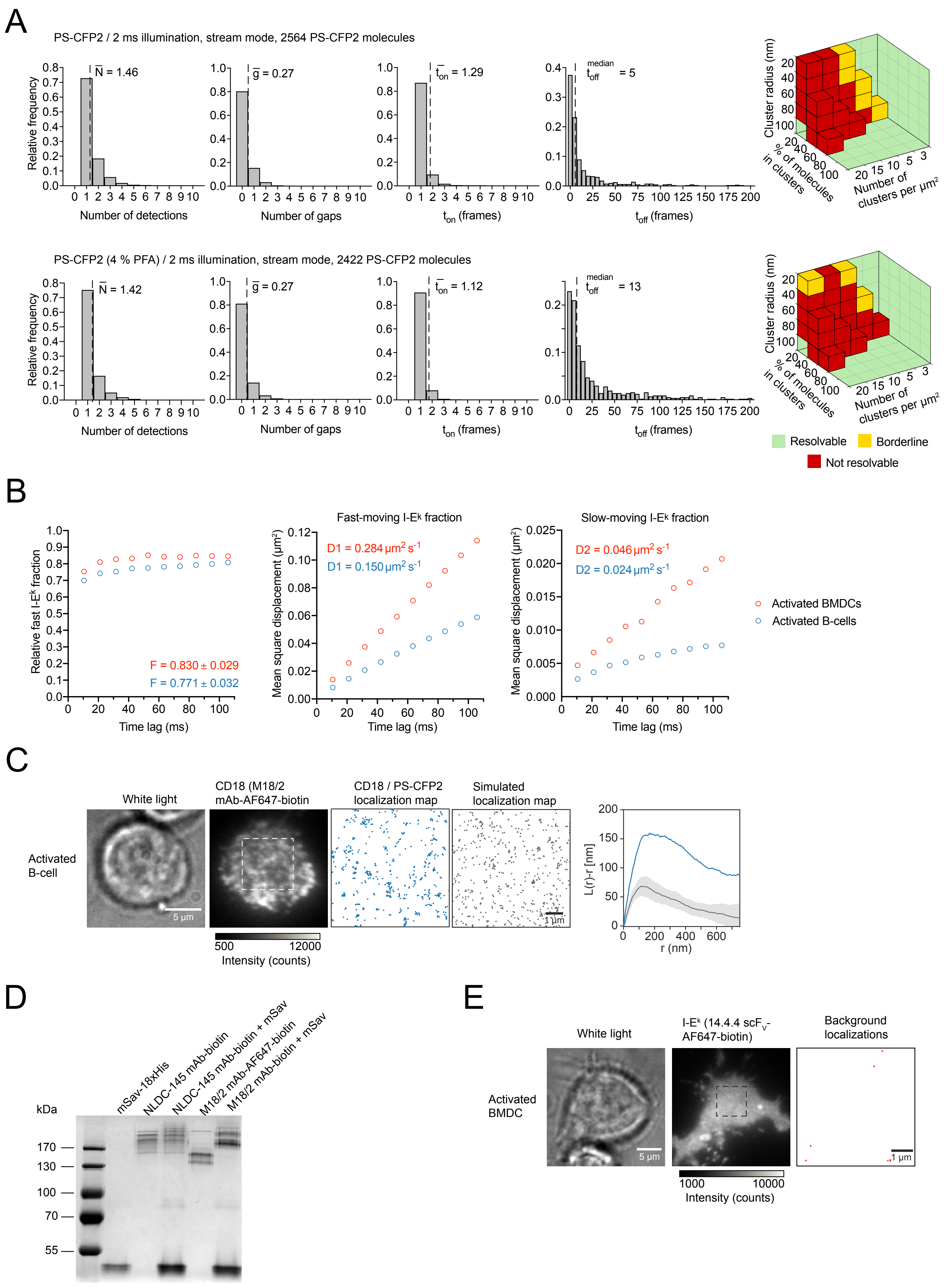

### Supplementary Figure 8

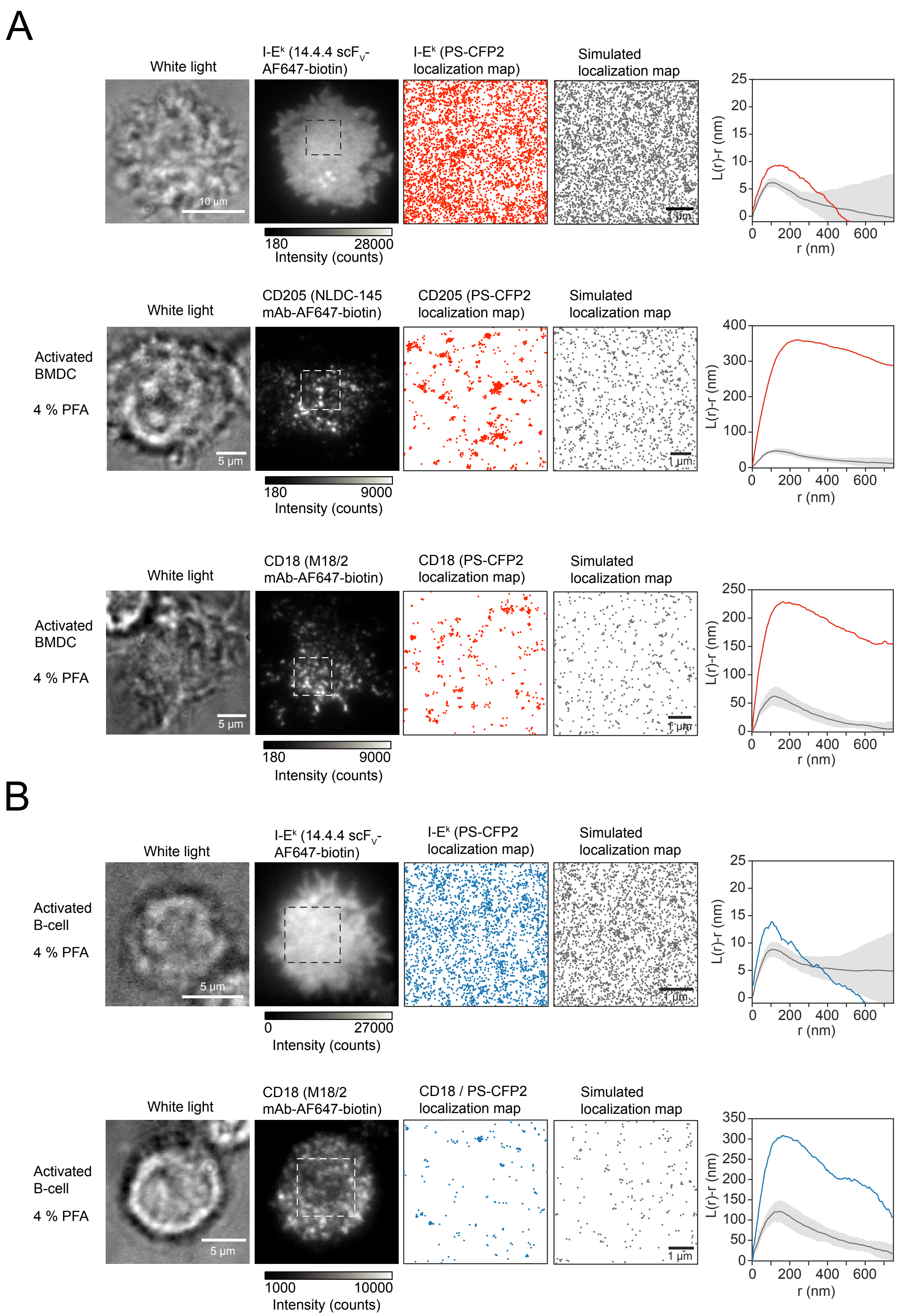

### Supplementary Figure 9

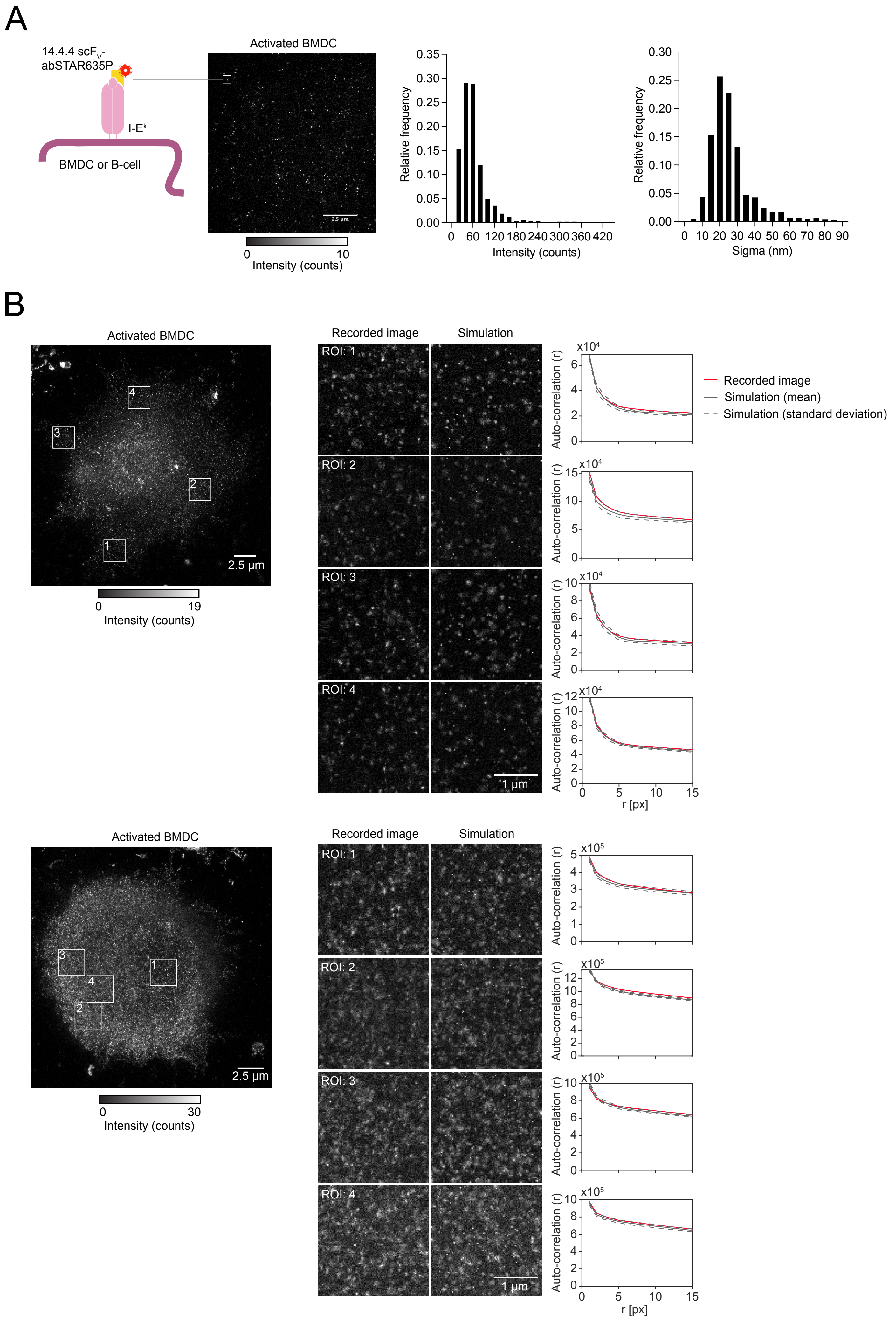

### Supplementary Figure 10

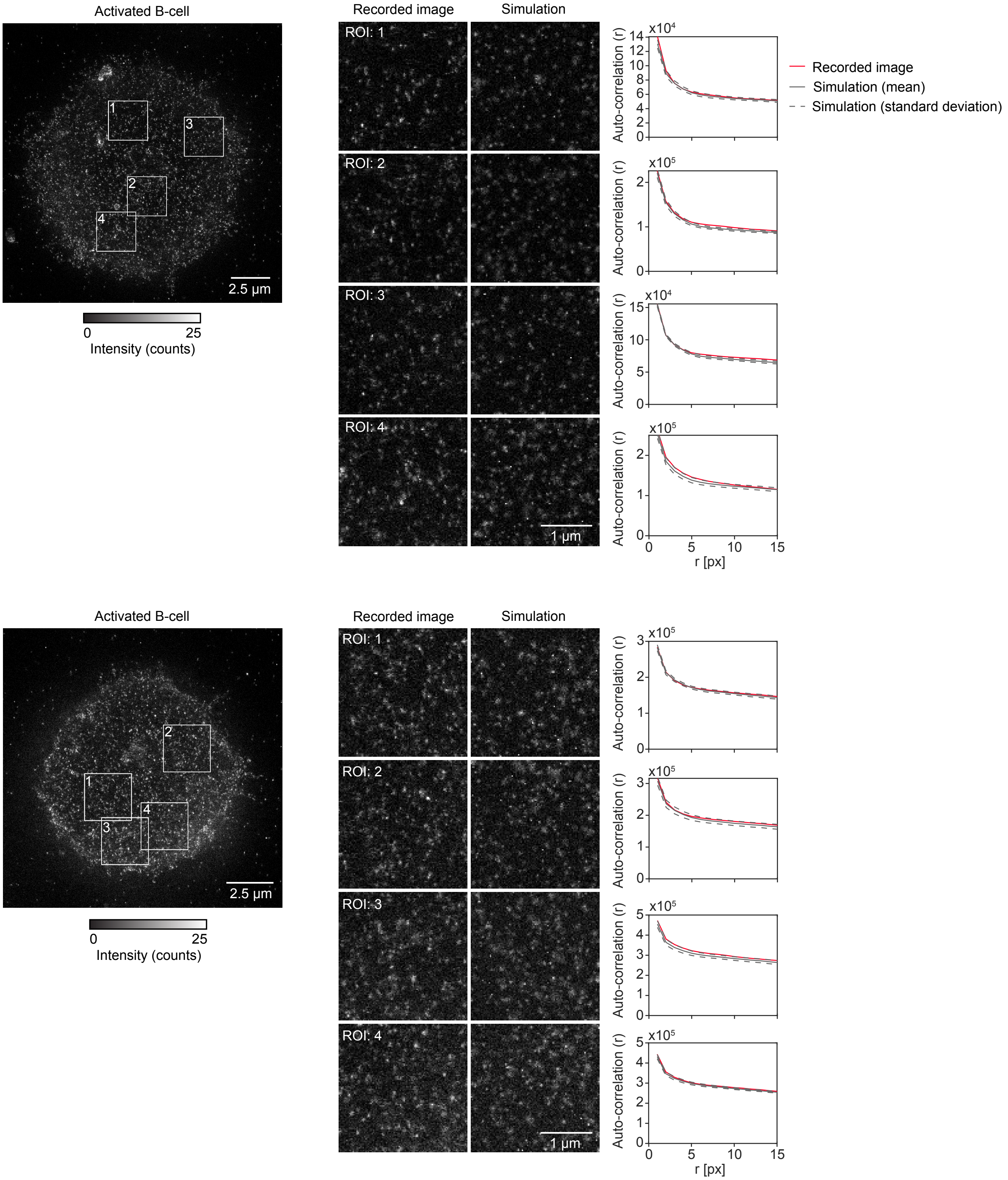

### Supplementary Figure 11

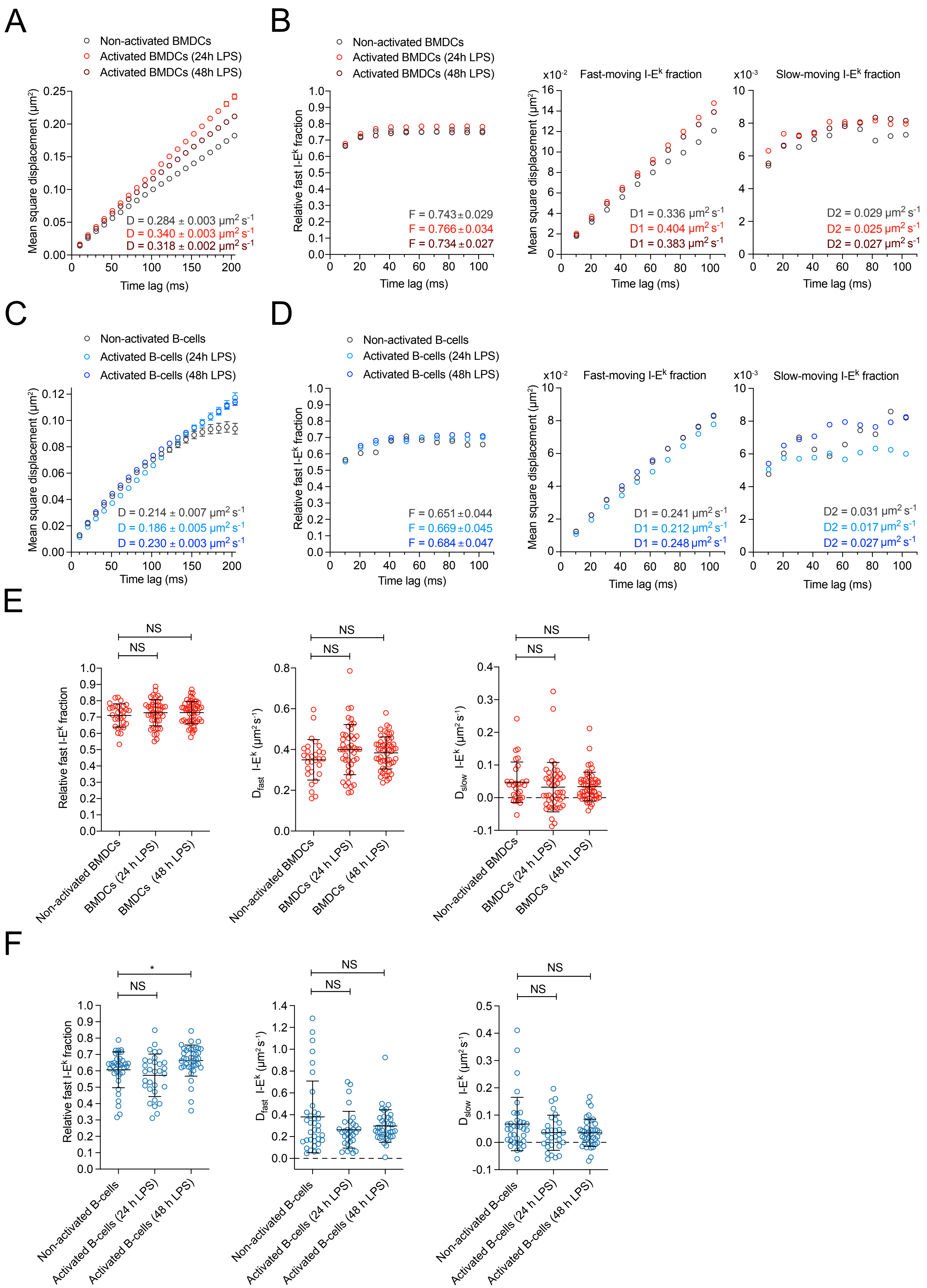

### Supplementary Figure 12

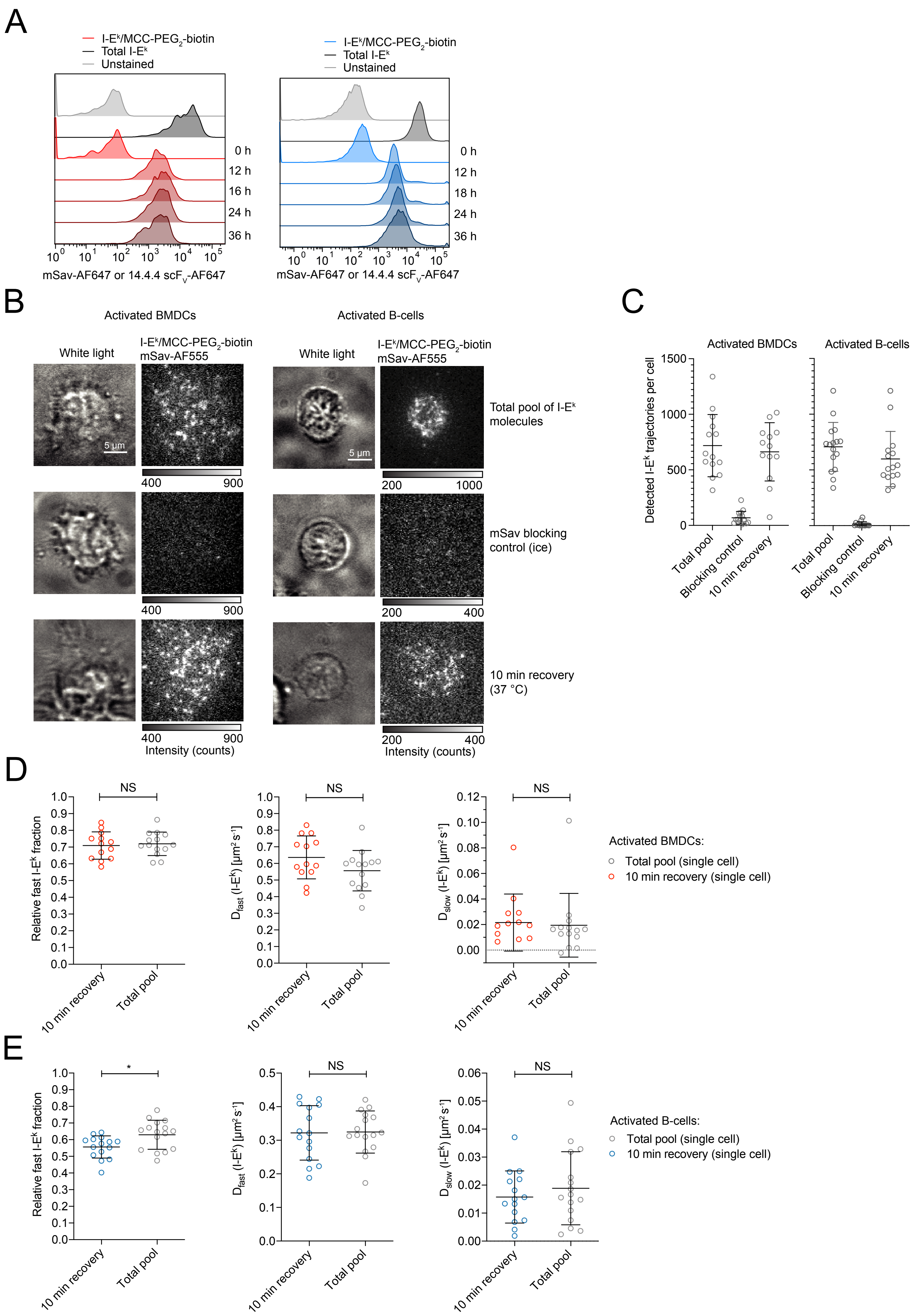

### Supplementary Figure 13

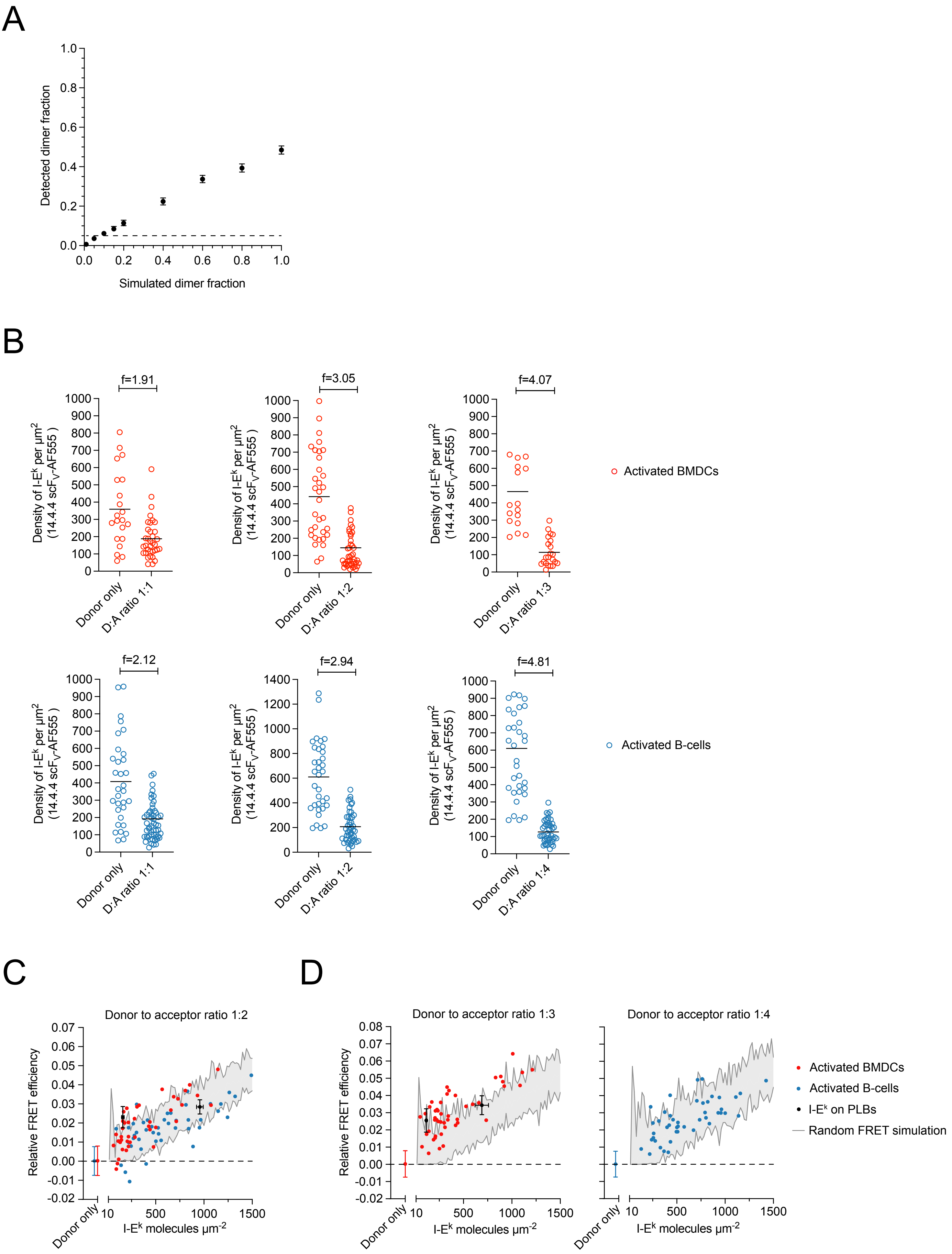

### Supplementary Figure 14

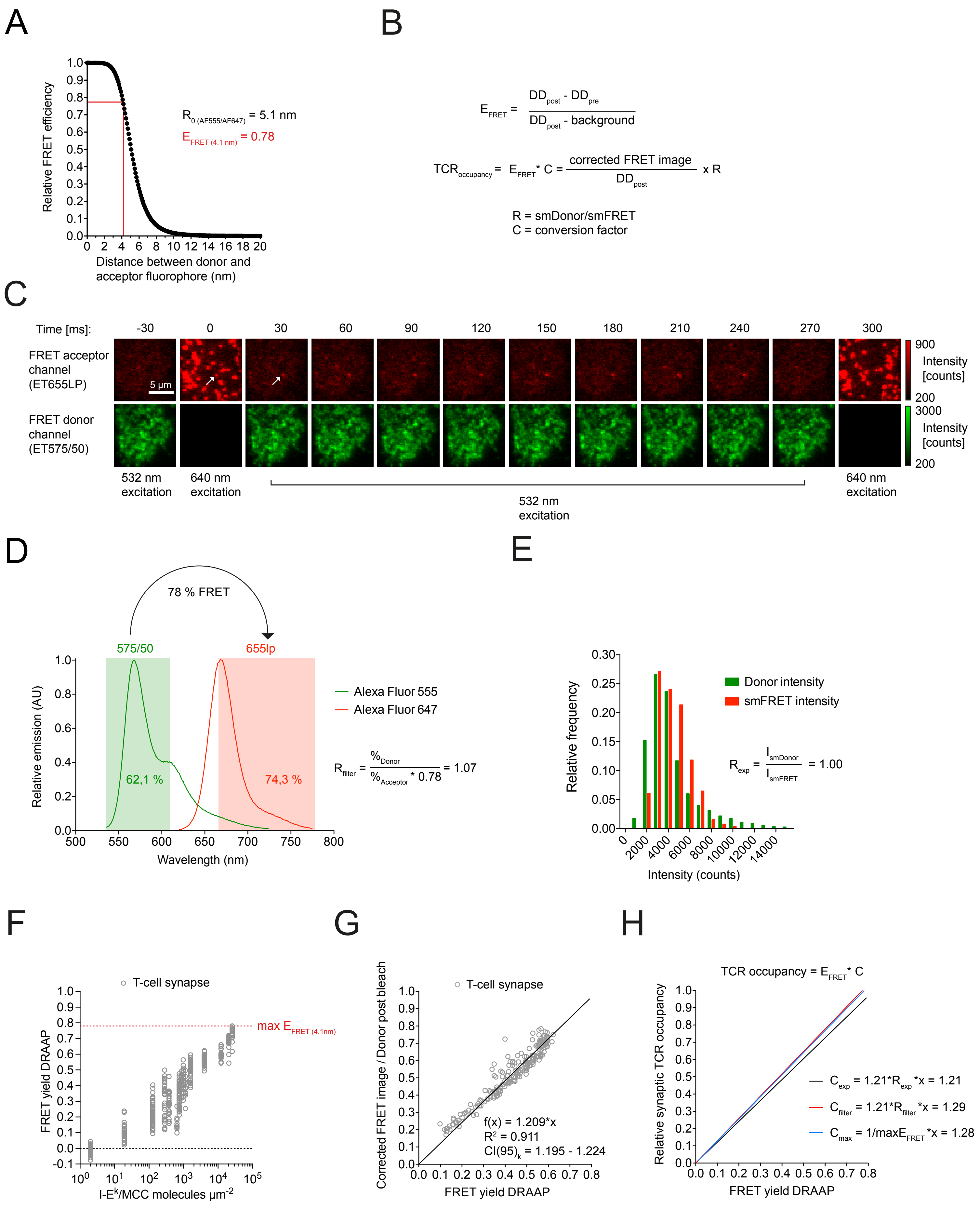

### Supplementary Figure 15

A

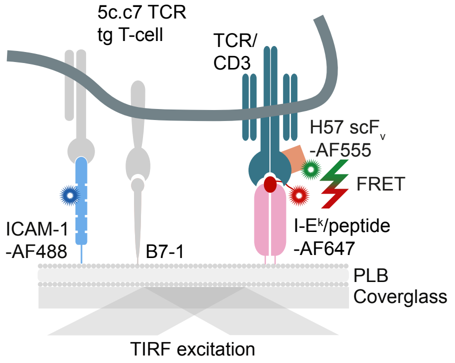


B

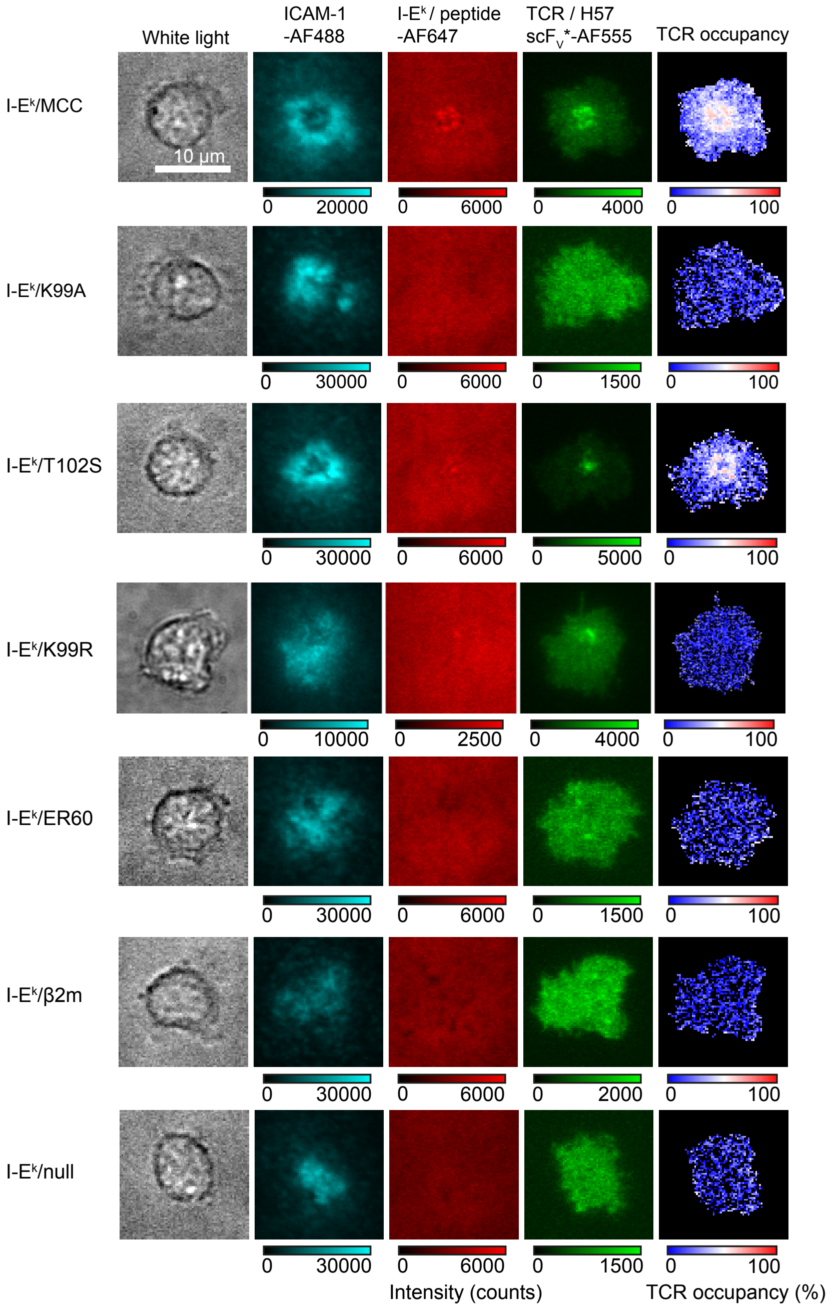

### Supplementary Figure 16

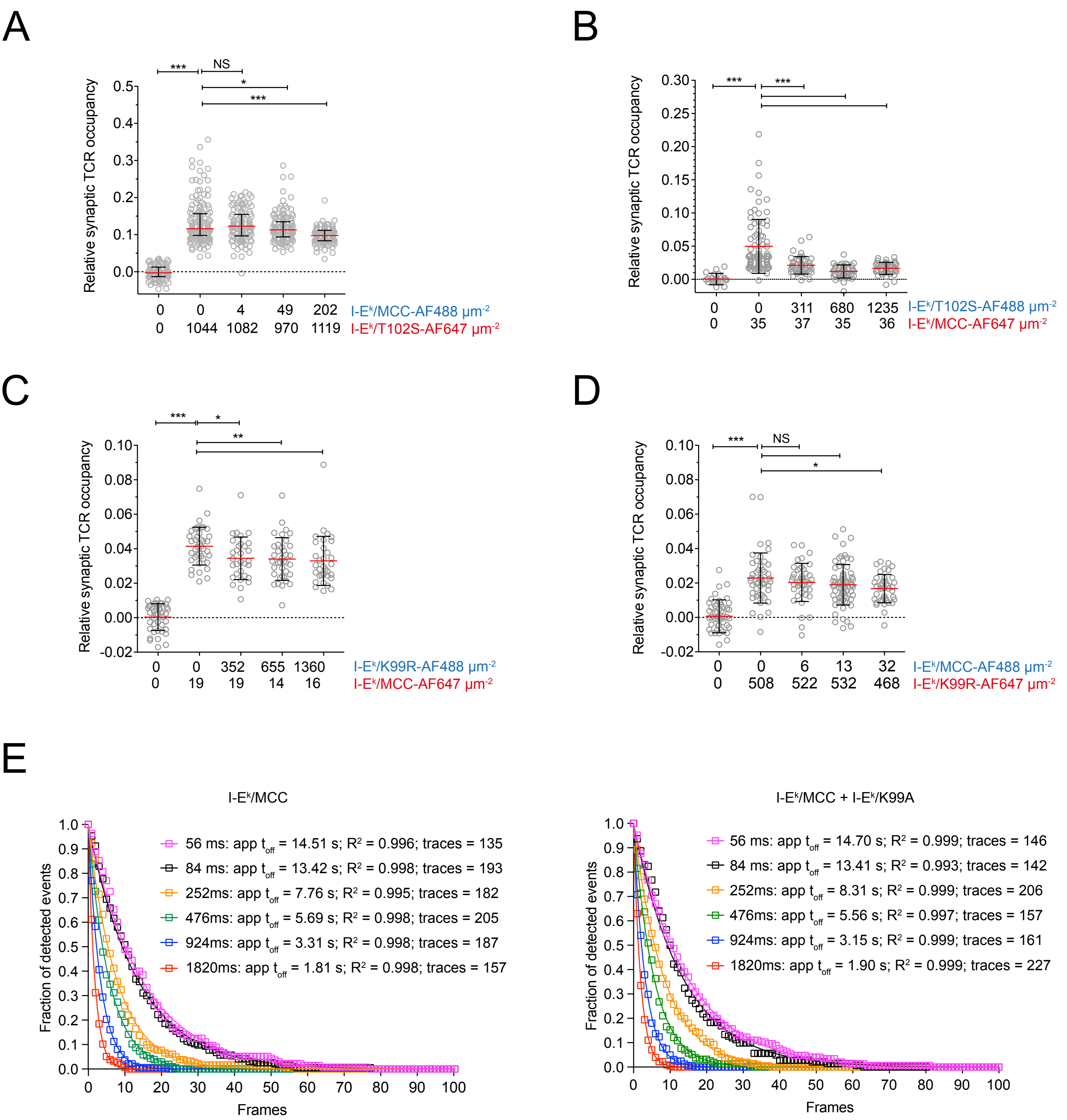

### Supplementary Figure 17

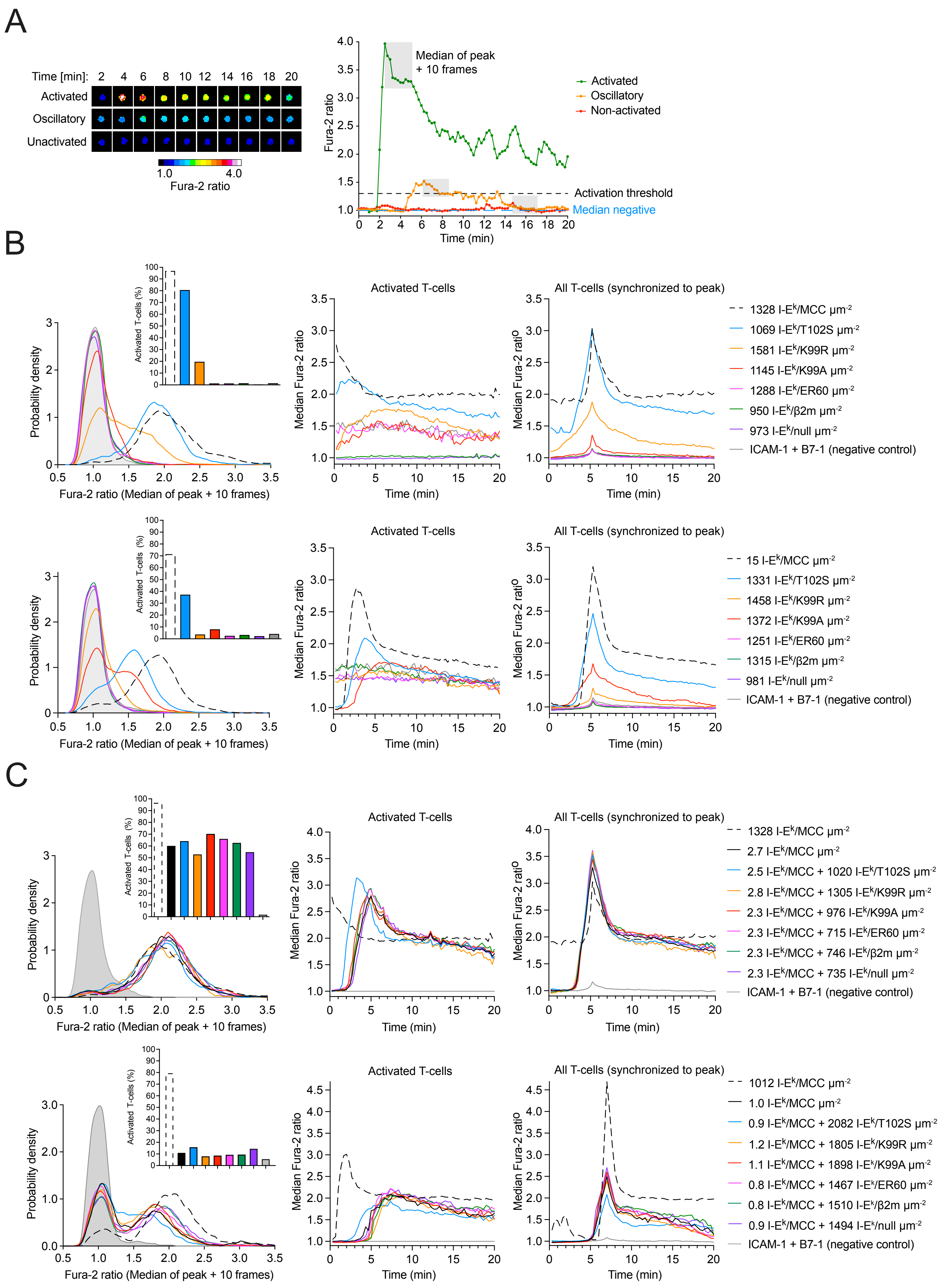

### Supplementary Figure 18

A

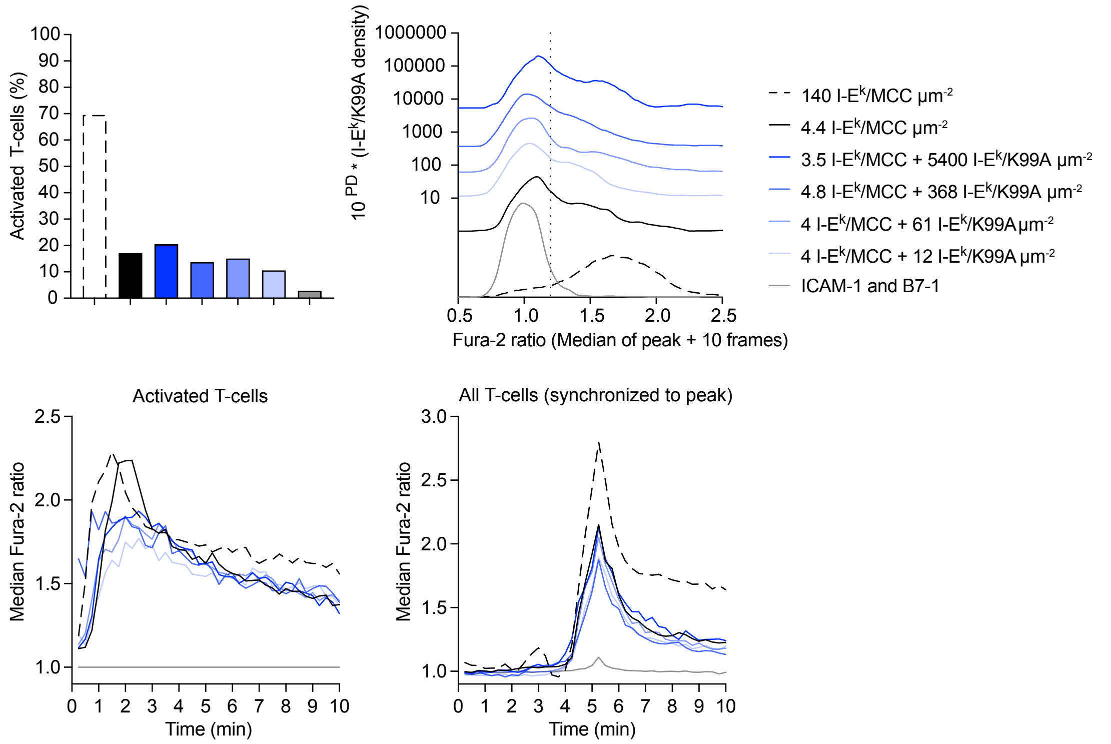


B

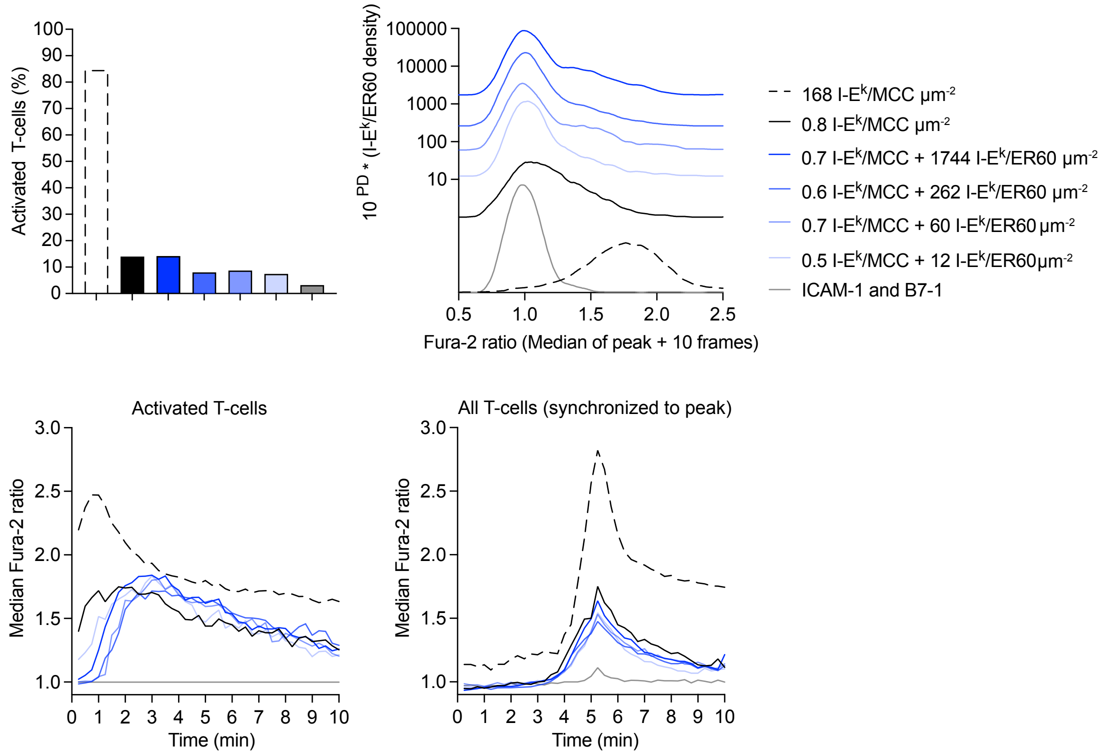

### Supplementary Figure 19

## A

```
             10         20         30         40         50         60         70
1    MQVQLQQSGP DLVKPGASVT ISCKASGYAF SSSWMSWLKQ RPGKGLEWIG WIFPRDGDTN YNGKFKGKAT
             80         90        100        110        120        130        140
71   LTADKSSSTA YMQLSSLTSE DSAVYFCARR GDYHYGMDYW GQGTSVTVSS AGGGGSGGGG SGGGGSDIVL
            150        160        170        180        190        200        210
141  TQSPASLAVS LGQRATISCR ASKSVSTSGY SYMHWYQQKP GQPPKLLIYL TSNLESGVPA RFSGSGSGTD
            220        230        240        249
211  FTLNIHPVEE EDAATYYCQH SRELPWTFGG GTKLEIKGS*--
             259       267
250  *GLNDIFEAQK IEWHEGSG--
```

## B

```
             10         20         30         40         50         60         70
1    CATATGCAGG TTCAGCTGCA GCAGTCTGGT CCGGACCTGG TTAAACCGGG TGCGTCTGTT ACCATCTCTT
             80         90        100        110        120        130        140
71   GCAAAGCGTC TGGTTACGCG TTCTCTTCTT CTTGGATGTC TTGGCTGAAA CAGCGTCCGG GTAAAGGTCT
            150        160        170        180        190        200        210
141  GGAATGGATC GGTTGGATCT TCCCGCGTGA CGGTGACACC AACTACAACG GTAAATTCAA AGGTAAAGCG
            220        230        240        250        260        270        280
211  ACCCTGACCG CGGACAAATC TTCTTCTACC GCGTACATGC AGCTGTCTTC TCTGACCTCT GAAGACTCTG
            290        300        310        320        330        340        350
281  CGGTTTACTT CTGCGCGCGT CGTGGTGACT ACCACTACGG CATGGACTAC TGGGGTCAGG GTACCTCTGT
            360        370        380        390        400        410        420
351  TACCGTTTCT TCTGCGGGTG GCGGCGGTTC GGGTGGCGGT GGCTCTGGTG GCGGCGGTTC TGACATCGTT
            430        440        450        460        470        480        490
421  CTGACCCAGT CTCCGGCGTC TCTGGCGGTT TCTCTGGGTC AGCGTGCGAC CATCTCTTGC CGTGCGTCTA
            500        510        520        530        540        550        560
491  AATCTGTTTC TACCTCTGGT TACTCTTACA TGCACTGGTA CCAGCAGAAA CCGGGTCAGC CGCCGAAACT
            570        580        590        600        610        620        630
561  GCTGATCTAC CTGACCTCTA ACCTGGAATC TGGTGTTCCG GCGCGTTTCT CTGGTTCTGG TTCTGGTACC
            640        650        660        670        680        690        700
631  GACTTCACCC TGAACATCCA CCCGGTTGAA GAAGAAGACG CGGCGACCTA CTACTGCCAG CACTCTCGTG
            710        720        730        740        750     760/814  762/816
701  AACTGCCGTG GACCTTCGGT GGTGGTACCA AACTGGAAAT CAAAGGATCC *TAATAAAAGC TT
             760        770        780        790        800   804
751  *GGCCTGAACG ATATTTTTGA AGCGCAGAAA ATTGAATGGC ATGAAGGTAG CGGC
```
